# Large scale analyses of genotype-phenotype relationships of glycine decarboxylase mutations and neurological disease severity

**DOI:** 10.1101/2019.12.20.884080

**Authors:** Joseph Farris, Barbara Calhoun, Md Suhail Alam, Shaun Lee, Kasturi Haldar

## Abstract

Monogenetic diseases provide unique opportunity for studying complex, clinical states that underlie neurological severity. Loss of glycine decarboxylase (*GLDC*) can severely impact neurological development as seen in non-ketotic hyperglycinemia (NKH). NKH is a neuro-metabolic disorder lacking quantitative predictors of disease states. It is characterized by elevation of glycine, seizures and failure to thrive, but glycine reduction often fails to confer neurological benefit, suggesting need for alternate tools to distinguish severe from attenuated disease. A major challenge has been that there are 255 unique disease-causing missense mutations in *GLDC*, of which 206 remain entirely uncharacterized. Here we report a Multiparametric Mutation Score (MMS) developed by combining *in silico* predictions of stability, evolutionary conservation and protein interaction models and suitable to assess 251 of 255 mutations. In addition, we created a quantitative scale of clinical disease severity comprising of four major disease domains (seizure, cognitive failure, muscular and motor control and brain-malformation) to comprehensively score patient symptoms identified in 131 clinical reports published over the last 15 years. The resulting patient Clinical Outcomes Scores (COS) were used to optimize the MMS for biological and clinical relevance and yield a patient Weighted Multiparametric Mutation Score (WMMS) that separates severe from attenuated neurological disease (p < 3.5e-5). Our study provides understanding for developing quantitative tools to predict clinical severity of neurological disease and a clinical scale that advances monitoring disease progression needed to evaluate new treatments for NKH.

## Introduction

Enzyme dysfunction underlies many pathologies, including a large number of neurological disorders, the metabolic consequences of which affect the central and/or peripheral nervous system. Clinical presentations of neuro-metabolic disorders include movement disorders[1], seizures, childhood epilepsies[2], and/or peripheral neuropathy[3]. Glycine decarboxylase (GLDC also known as P-protein) is an enzyme that catalyzes the cleavage of glycine, the first step of the mitochondrial glycine cleavage system (GCS). Other GCS components are aminomethyl transferase (*AMT*; T-protein), glycine cleavage system H-protein (*GCSH*), and dihydrolipoyl dehydrogenase (*DLD*; L-protein). Loss of GLDC-protein activity completely abrogates GCS function. Catabolism of glycine by the GCS is an essential metabolic process. Degradation of glycine feeds into one-carbon folate metabolism through the formation of 5,10-methylene THF[4], which in turn is utilized to synthesize nucleotides and proteins. Perturbations in the GCS are involved in a number of disease states, including cancer and neural tube defects (NTDs)[5–7]. Loss of function mutations in *GLDC* are the primary cause for the rare neuro-metabolic disorder non-ketotic hyperglycinemia (NKH), accounting for approximately 85% of NKH cases[8].

NKH affects approximately 1 in 76,000 births[8], although some populations, such as the Finnish, have a significantly higher rate due to founder mutations and consanguinity[9, 10]. A high incidence of NKH has also been reported amongst the Amish[11]. NKH results from loss of GLDC- or (to a lesser degree) T-protein activity. This causes an acute increase of glycine in plasma and cerebral spinal fluid (CSF)[12]. But plasma glycine is not predictive of clinical severity; furthermore, it is a challenge to continuously monitor glycine in the CSF. NKH is typically characterized as either severe or attenuated type [12, 13]. Severe NKH causes intractable seizures, failure to thrive, lack of developmental milestones and often premature death. Attenuated patients are often able to control seizures, go to school, and live into adulthood. But lack of mutation-based predictors of disease progression and a quantitative scale of disease severity scale impedes both management of NKH as well as development of therapies to treat and cure the disease.

In this study, we developed a multi-parametric mutation scale applicable to all but 4 of 255 missense NKH mutations across the *GLDC* gene and thereby assigned a multi-parametric mutation score (MMS) to 251 patient mutations. The MMS was further optimized against a newly developed patient-based clinical outcomes score that was based on major symptomatic domains extracted from a comprehensive review of clinical cases reported in the literature. Our findings yield a quantitative tool with high predictive value to support disease management and emerging treatments associated with >95% of known clinical NKH mutations.

## Results

### Positional distribution of NKH Missense Mutations

The most recently published comprehensive list of NKH missense mutations lists 171 unique mutations across the length of GLDC [8]. To this, we added 85 missense mutations based on additional literature and the Clinvar database. This yielded a list of 256 mutations ascribed to 213 unique residues (Table S1, Fig 1A). NKH missense mutations were found in the C-terminus, which houses most of the GLDC active site, as well as the N-terminus, suggesting substantial distribution throughout the protein. Of the twelve of the most frequently mutated residues (which give rise to 18 mutations; Fig 1B) five appear in the N-terminus and seven in the C-terminus, and are not concentrated in any specific domains (based on conservation analyses as well as domains predicted from primary and secondary structure; Fig 1C). One of the 256 mutations is located in the mitochondrial leader sequence (M1I). Of the remaining 255, only 49 mutations have been assessed for loss of enzymatic activity compared to the wild type. Of these, only 9 have been analyzed for their potential effect on GLDC-protein structure based on *in silico* 3D-modeling[14]. Together, the findings summarized in Fig 1A-C show the overall lack of annotation for NKH mutations and demonstrate the need for improved tools and analyses to better understand mutations across the gene, how they may cause deficiency in the encoded protein and thereby impact disease severity.

**Figure 1.**
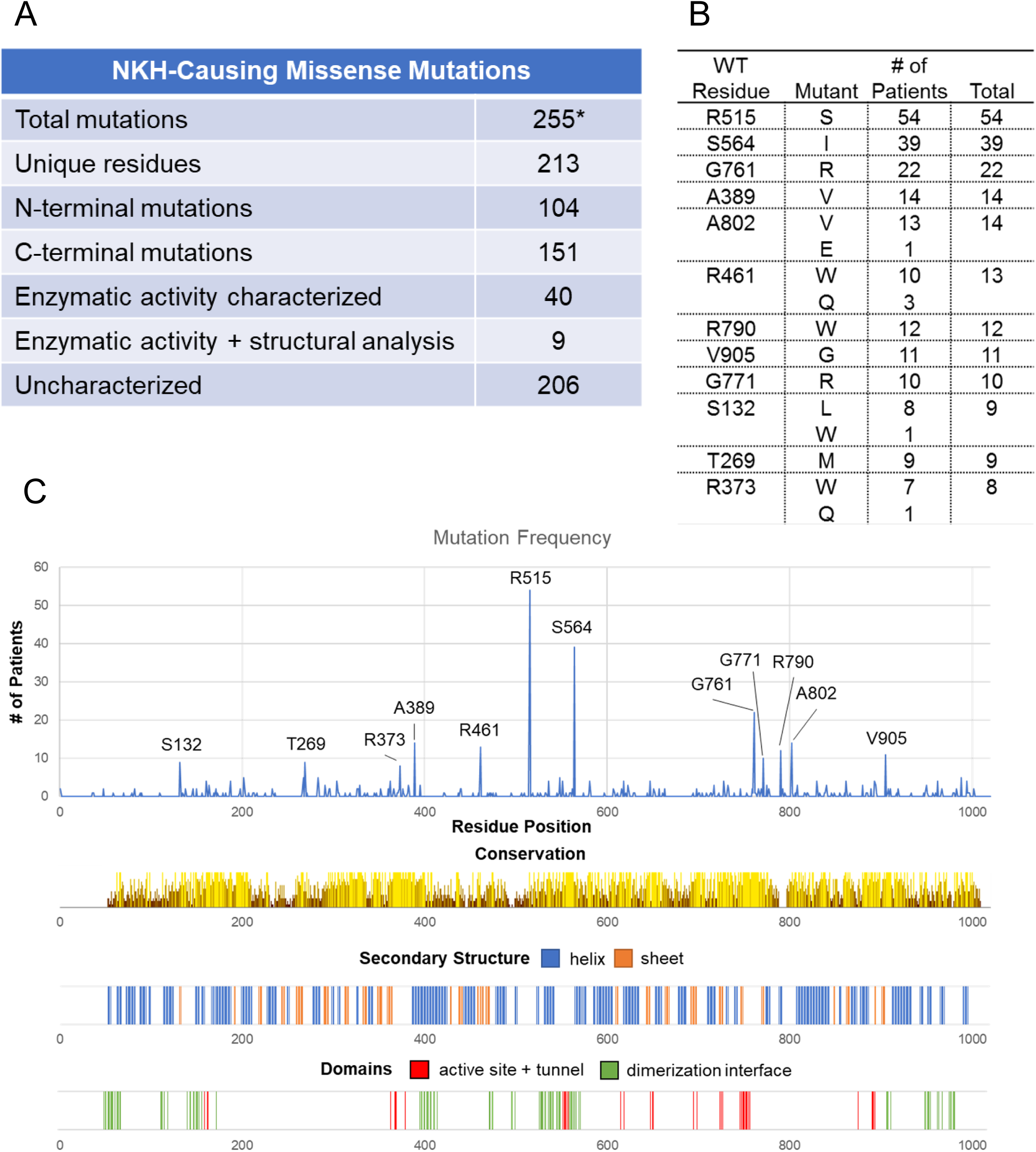
Survey of NKH-causing missense mutations. [A] Summary of NKH mutations. *255 unique missense mutations (barring the M1I mutation in the leader sequence), were compiled from the published clinical literature and the ClinVar database[13,14,53–62,29,63–68,46–52]. Only 49 mutations have been characterized (to varying degrees), leaving 206 uncharacterized. [B] The top ten most frequently observed mutated positions in GLDC (that account for 14 unique mutations). [C] Distribution of NKH-causing missense mutations across the length of the GLDC-protein. Conservation of amino acids and secondary structure, as well as the positions of the active site, active site-tunnel, and dimerization domains are shown.

### Structural Annotation of GLDC

#### i. Comparative structural analyses

Since comparative structural analysis provides a robust path to understanding functions of conserved domains, we generated a high-confidence homology model for human P-protein using *Synechocystis sp. 6803* PLP-bound glycine decarboxylase (PDB: 4LHC) as a template (Fig 2A). The model in Fig 2A shows a global mean quality estimate (GMQE) of 0.77 (where 1 is the highest possible score). Most of the uncertainty in this model comes from a flexible loop consisting of amino acids 360-384. This region is missing in the bacterial orthologue catalogued in the Protein Database because the region can exist in either a disulfide-bridge or open form, resulting in a low electron density[15]. SWISS-model predicted this region to be in the open conformation, which is the conformation seen in the holoenzyme[15].

**Figure 2.**
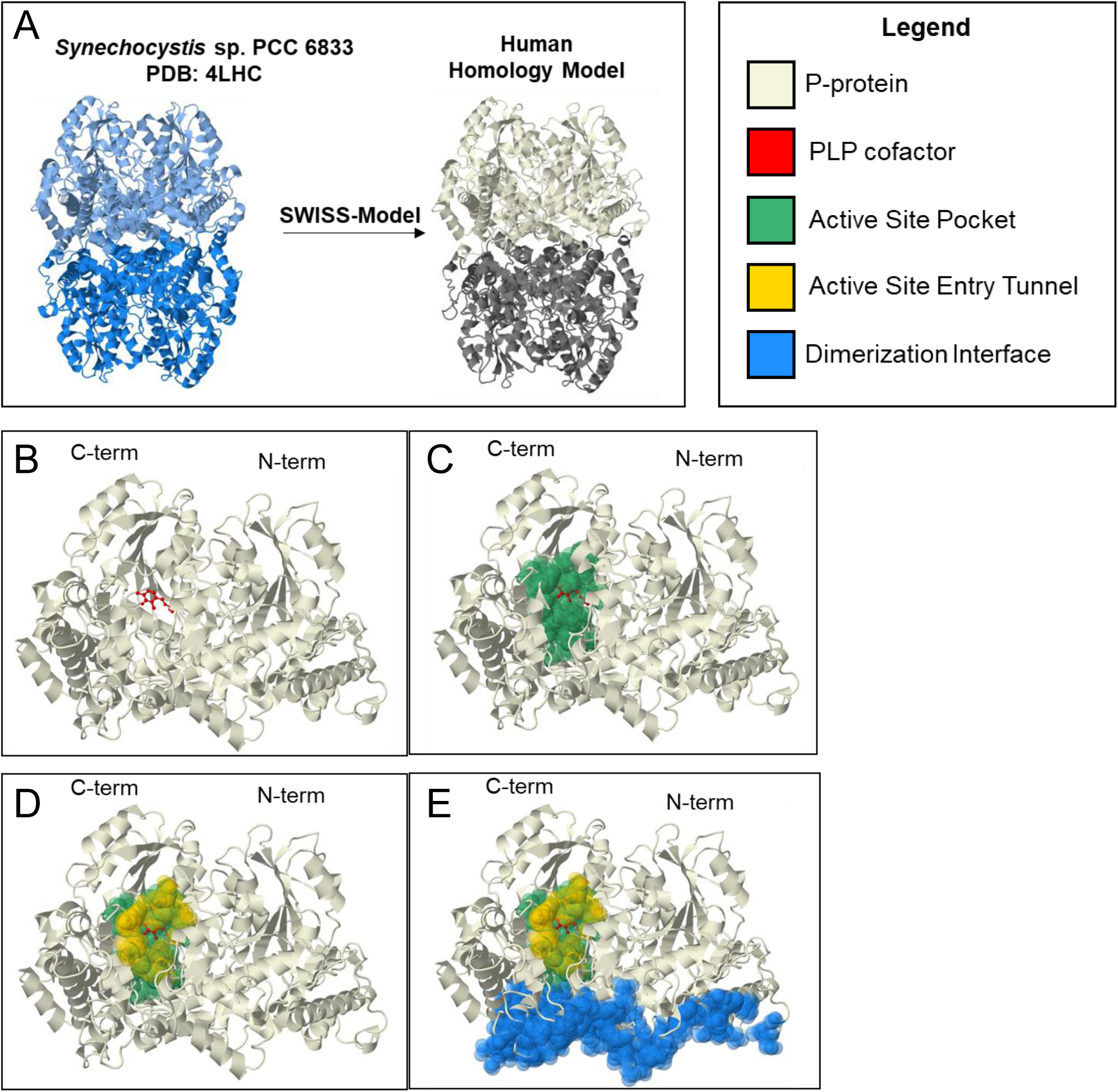
Human homology model of P-protein. [A]. A crystal structure of *Synechocystis* sp. PCC 6833 GLDC holoenzyme (PDB = 4LHC) (LHS) was the template used in SWISS-Model to generate the human homology model (RHS). Based on homology, human GLDC is predicted to form an α_2_ homodimer. α-subunit shown in beige. α’ shown in gray. [B – E] Evolutionary derived, functional regions of human GLDC model with. [B] cofactor PLP (shown in red); [C] active site pocket (green) defined as residues within 5 Angstroms of PLP containing active site lysine (K754), [D] a tunnel (shown in yellow) that opens at the surface when PLP is bound making the active site pocket accessible to lipoylated H-protein; [E] the dimerization interface of GLDC (blue).

GLDC’s enzymatic function is dependent on the binding of its cofactor pyridoxal phosphate (PLP) at Lys754 (Fig 2B). Active site residues (Fig 2C) were defined as any amino acids within 5 Angstroms of the PLP-bound Lys754 or substrate glycine (not shown). Active site tunnel residues (Fig 2D) were defined as residues equivalent to those constructing the active site tunnel observed in the bacterial structure (based on sequence alignment). The dimerization interface (Fig 2E) was defined as residues predicted to be within 5 Å of the GLDC-protein α’-subunit of the αα’ dimer (although there is no direct evidence that dimerization is required for enzymatic function). These three conserved, functional regions provide annotation for much of the C-terminus and they account for 40 NKH-causing mutations, only seven of which have been previously characterized.

#### ii. N-terminal Active Site Function

Although comparisons with the bacterial structure accounted for mutations at and around the active site, they failed to explain the presence of high density of pathogenic mutations in N-terminal regions, that appear to be devoid of functional annotation. We undertook additional evolutionary analyses to predict N-terminal domain function of GLDC. GLDC belongs to the PLP-enzyme Fold Type I family[16]. However, it is an unusual member as all other PLP Fold Type I enzymes in this family form α_2_ homodimers (where each α-subunit is ∼500 amino acids) while GLDC is either a single polypeptide over 1000 amino acids or, in some bacteria and archaea, an αβ heterodimer, with the α- and β-subunits corresponding to the N- and C-termini respectively (Fig 3A). In these αβ orthologs, there is structural and sequence similarity between the α and β subunits, leading to the suggestion that they came from the same evolutionary precursor[17]. We found that in human GLDC-protein, pBLAST alignment of the N- and C-termini shows a region of similarity (26% identity, 48% positives) between amino acids 238-349 and 572-785 (S1 Fig). Notably, the C-terminal region contains the active site, Lys754. In addition, large regions of the N-terminus (113-472) and C-terminus (531-906) are significantly structurally similar by rigid FATCAT alignment (p = 3.61e-13, RMSD = 2.59Å; Fig 3B). The structure of the C- and N-termini also both show similarity to other Fold Type I carboxylases. The defining feature of this fold is the 7-stranded β-sheet packed by α-helices[18], which is evident in both N- and C-termini (Fig 3C).

**Figure 3.**
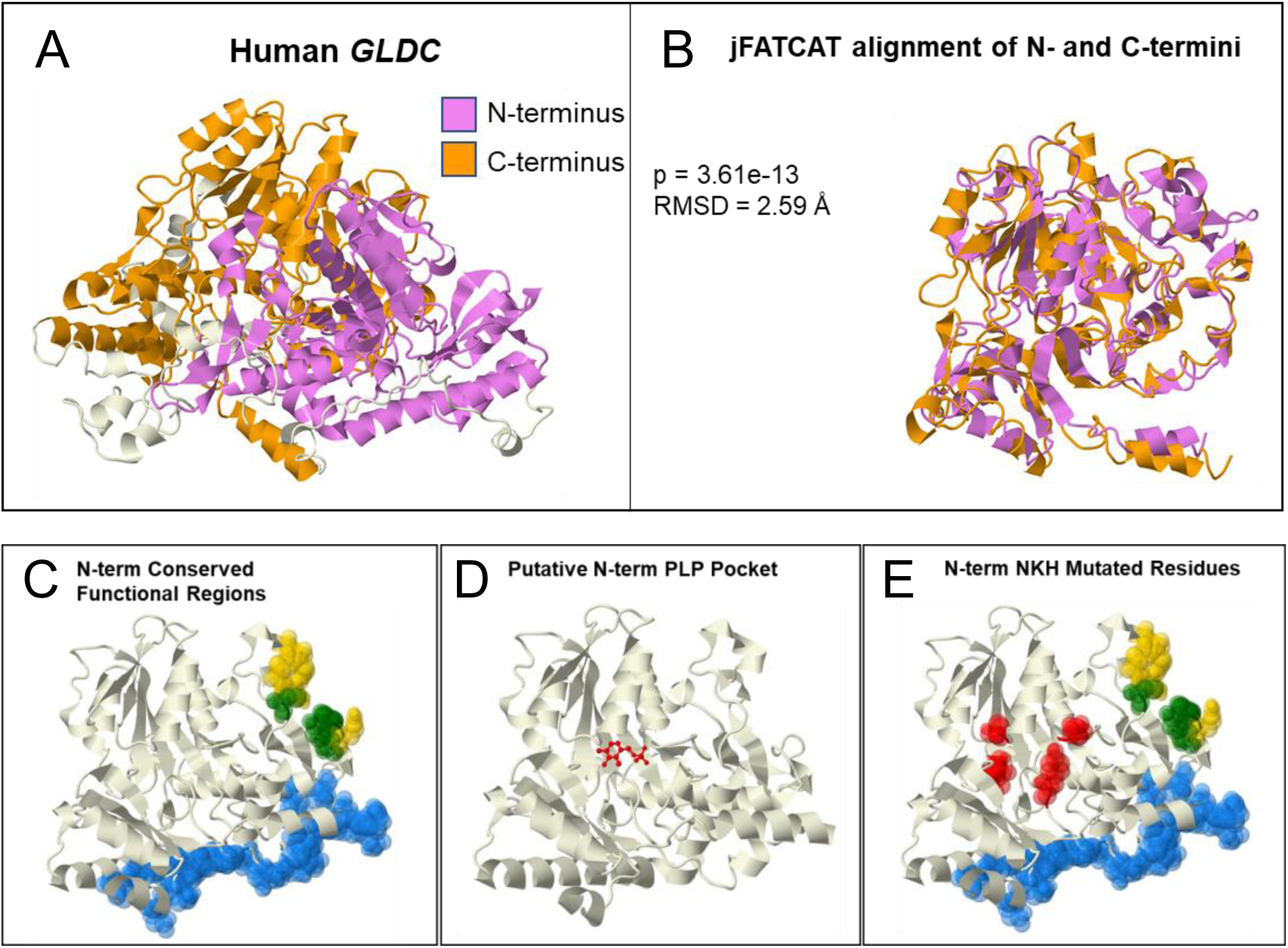
GLDC N-terminal Structural Homology and Function. [A] Structurally homologous regions of the N- and C-termini respectively shown in purple and orange, respectively. [B] Relative alignment of the N- and C-terminal regions shown in [A]. FATCAT alignment gives an RMSD of 2.59 Å and a p-value of 3.61e-13. [C] Functional regions predicted based on evolutionary conservation in the N-terminus: tunnel, yellow; active site, green, and the dimerization interface, blue. [D] I-Tasser COFACTOR predicted PLP binding site in the N-terminus (PLP shown in red), providing a rationale for the presence of [E] a high incidence of NKH disease mutations in region shown in red.

Taken together, these analyses suggested a conserved PLP-binding fold in the N-terminus. Accordingly, I-Tasser COFACTOR prediction for P-protein predicts the N-terminus to be a PLP-binding site based on structural similarity to other PLP-binding enzymes (Fig 3D), although with a low confidence score of 0.05 (range: 0-1). However, the known C-terminus PLP-binding site was not predicted by I-Tasser, enabling us to retain the N terminus projection (despite the low score). Intriguingly, however, the N-terminal PLP Fold is lacking an active site lysine, having instead a glutamine at that position. However, site-directed mutagenesis of another PLP Fold Type I enzyme’s active site lysine showed that lysine, while essential for catalysis, was not essential for PLP-binding[19]. Thus, we do not predict that this pocket has enzymatic activity. Rather, we predict that the N-terminus non-covalently binds PLP. It has been observed that PLP is needed for GLDC-protein to fold properly[20], suggesting a rationale for why 11 clinical NKH-mutations cluster in the fold-predicted PLP-binding region (Fig 3E; which has no other known function).

#### iii. Macromolecular Interaction of H-Protein with GLDC-protein

The interaction of GLDC with H-protein is essential for GLDC function, but the molecular coordinates of the interaction remain unknown. A lipoyllysine group on H-protein accepts the amino-methyl moiety produced by decarboxylation of glycine by GLDC-protein and transfers the moiety to T-protein[21]. We predicted that this lipoyllysine accesses the active site through the observed active site tunnel. To model this interaction, a homology model for H-protein was generated with SWISS-Model using a published crystal structure for bovine H-protein (98% sequence similarity to human H-protein) as the template. The ClusPro 2.0 server was used to model the interaction of H-Protein with the homology model for human GLDC-protein. ClusPro 2.0 produced 101 potential interaction models (S1 Appendix). To select the best model, a novel scoring system was developed based on conservation of the two proteins’ interacting residues and proximity of the modified lysine of H-protein to the entry of the active site tunnel in GLDC-protein. Conservation was chosen to select the best model because highly conserved protein surfaces are a predictor of a protein binding site[22]. Accordingly, GLDC-protein contains a highly conserved region at the site where H-protein should bind (Fig 4A). To further test our conservation parameters, we scored the interaction between human T-protein with H-protein (which are expected to interact) and T- and GLDC-protein (not expected to interact). Like the H-GLDC interaction, the H-T interactions had scores close to the max of 2 while the T-GLDC interaction did not (Fig 4B), confirming the utility of this method. To test the accuracy of all three scoring parameters, the interaction between *E. coli* T- and H-protein was modeled using ClusPro 2.0 (S1 Appendix) and scored (S1 Table). The highest-ranking computational docking interactions were compared to the previously published crystal structure of the docking interaction[23], and it was found that the second highest scoring model (score = 2.84 of 3) was significantly similar to the crystal structure (S2 Fig).

**Figure 4.**
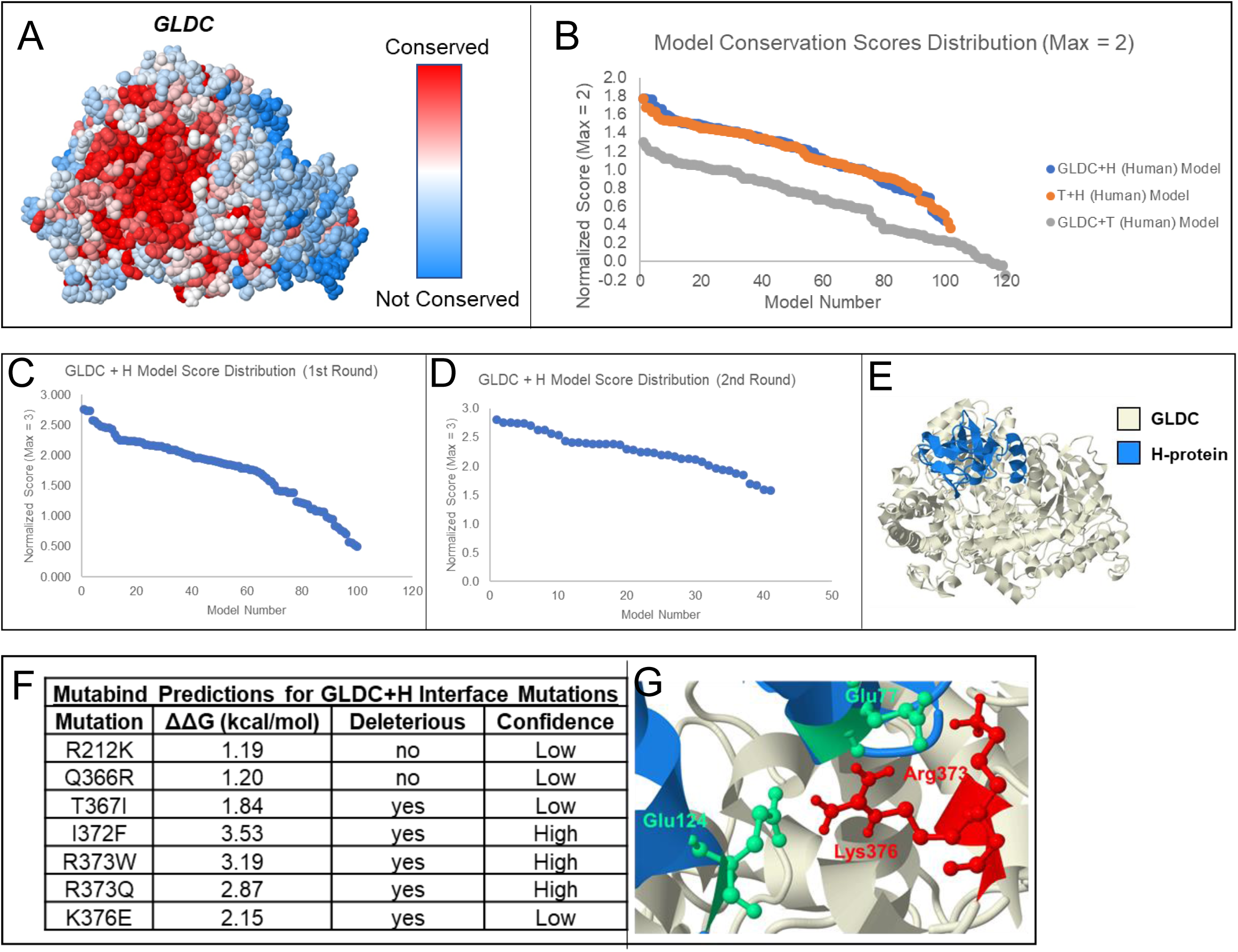
Model of the interaction between GLDC and H-Protein of the Glycine Cleavage System. [A] Model showing amino acid conservation scores in GLDC calculated using the Consurf server. Highly conserved amino acids are shown in red, while poorly conserved residues are shown in blue. The highly conserved surface (deep red) found at the entry of the active site tunnel suggests this region may be a binding site for H-protein. [B] Scoring of models generated for the GLDC + H interaction (blue), T + H interaction (orange), and the GLDC + T interaction (gray). Conservation of interacting surfaces was scored from 0-1, with the most conserved surface assigned a score of 1. GLDC and T-protein likely do not interact, thus their interaction was included as a negative control. Accordingly, the GLDC + T scores were lower than the GLDC + H and H + T interaction scores, validating our novel scoring method. [C] Scoring of GLDC + H-protein interaction models scored using the conservation of interacting amino acids with the distance between the H-protein and the entry to the GLDC-protein active site tunnel as an additional parameter. H-protein was allowed to bind to any location on the GLDC-protein surface. [D] H-protein was constrained to a region of GLDC protein based on the highest scoring results of the first round of scoring, and scoring was repeated for the resulting models. [E] The proposed H- (blue) and GLDC-protein (beige) docking interaction. [F] Seven known NKH mutations were found at the predicted H-GLDC interface. The ΔΔG caused by the mutation was estimated using Mutabind. 5 of 7 mutations are predicted to caused deleterious effects to the H-GLDC protein interaction, with 3 being high confidence predictions. [G] Salt bridges between Glu77 and Glu124 of H-protein and Arg373, and Lys376 of GLDC-protein. These GLDC-protein residues are known to cause NKH when mutated.

We performed two rounds of H- and GLDC-protein scoring to ensure that we obtained the highest-possible scoring model (S2 Table; Fig 4C-D) with a score of 2.81 out of 3 (Fig 4D). This model was selected as our predicted model of the interaction between human GLDC- and H-protein (Fig 4E). Finally, of seven known NKH-mutations at the predicted H-GLDC interface, five are predicted by Mutabind to negatively affect the interaction between H- and GLDC-protein (Fig 4F). These mutations point to the importance of the salt bridges formed between Arg373 and Lys376 of GLDC-protein and Glu77 and Glu124 of H-protein for stabilizing the interaction (Fig 4G). Thus, our interaction models provide crucial information for these six residues with NKH-mutations that were previously not understood.

As summarized in Table 1, the rates of mutation in the N-term PLP binding site (0.3) and H-GLDC interface (0.35) are higher than the baseline mutation rate in the protein (0.21) and approach those seen in the active site (0.33) (Table 1), further validating the functionality of these newly predicted regions.

**Table 1.** Mutation Rate in Predicted Structural Regions

| **Region** | **Whole Protein** | **Full Active Site** | **N-term PLP** | **H-interface** |
| --- | --- | --- | --- | --- |
| **Total Residues** | 1020 | 30 | 20 | 17 |
| **# Mutated** | 213 | 10 | 6 | 6 |
| **Residue Mutation Rate** | 0.209 | 0.333 | 0.300 | 0.353 |

#### iv. Large-scale analysis of disease mutations

Comparative structural and evolutionary analyses undertaken in i-iii, enabled annotation of 51 previously uncharacterized NKH mutations. Although this reflects a 100% increase in mutation annotation, a large number of mutations (∼150) remain in need of annotation. Since many missense mutations are expected to impact protein folding, we initiated large-scale studies that incorporate the assessment of the Gibbs free energy changes (ΔΔG) that arise as a consequence of mutation, as well as other changes in intrinsic properties of amino acids. ΔΔG provides a benchmark measure to predict the change in stability of a monomeric protein caused by a point mutation. Although many predictive online tools exist[24], we used CUPSAT because it makes fast and accurate ΔΔG predictions and is thus ideally suited for the large number of missense mutations seen in NKH and GLDC. We provided the SWISS-Model generated GLDC homology model as the input and defined destabilizing mutations as those with predicted ΔΔG < −1.5 kcal/mol (see Methods) to yield stability predictions for 251 of 255 missense mutations (see S3 Table). The remaining four mutations in a small, uncrystallized region at the beginning of the N-terminus could not be assessed and were not pursued further.

Fig 5A provides a pictographic representation that suggests that of the 251 mutations, 105 were predicted to be destabilizing (< −1.5 kcal/mol) with 42 being very destabilizing (< −5 kcal/mol). Destabilizing mutations were seen throughout the protein with 44 being found in the N-terminus and 61 being found in the C-terminus (S3 Table). The two most common NKH clinical mutations, both of which are known to cause severe disease are predicted to be destabilizing (R515S ΔΔG = −2.47 kcal/mol; S564I ΔΔG = −3.23 kcal/mol). In total, 4 (R515S, S561I, G771R, and V905G) of the top 10 most common missense mutations are predicted to be destabilizing. But the majority of mutations are predicted to have (i) negligible effect (ΔΔG = −1.5 to 1.5 kcal/mol; N = 96), (ii) stabilizing effect (ΔΔG = 1.5 to 5 kcal/mol; N = 35), or (iii) very stabilizing effect (ΔΔG > 5 kcal/mol; N = 15). This suggests that while ΔΔG provides valuable information on predicted stability for NKH missense mutations, it is not sufficient as a comprehensive parameter for the impact of these mutations.

**Figure 5.**
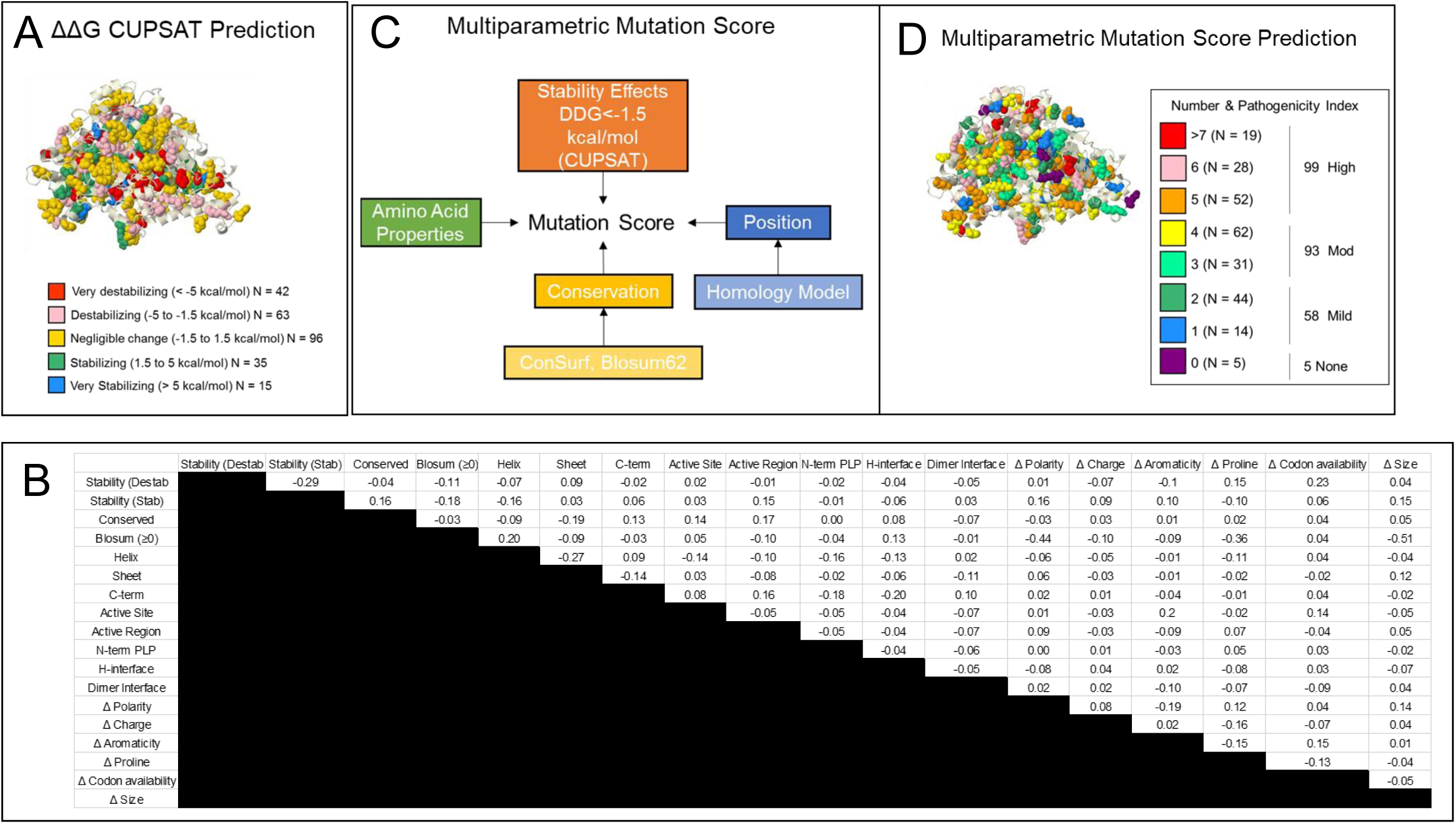
Multiparametric Mutation Score (MMS). [A] Stability effects. CUPSAT-predicted stability effect of NKH mutations on GLDC-protein assessed by ΔΔG. [B] Phi correlations of 18 parameters predicted to impact protein stability. Most parameters have no correlation with a phi-value of < |0.3|. Change in proline, size, and polarity have weak negative correlations with the conservation of amino acid change parameter, indicating that alterations in these amino acid properties are not well-tolerated through evolution. [C] Design for calculating MMS. In total, 18 parameters were used from 4 general categories of stability effects, mutation position, conservation, and change in amino acid properties (see Methods). [D] MMS was calculated from a summation of the 18 parameters, each with a weight of 1 except conservation of substitution, which was weighted −1. 3-dimensional distribution of MMS’s shown for all known NKH missense mutations.

Multiparametric mutation scores (MMS) that incorporate ΔΔG and other protein parameters, have previously been successfully used to predict missense mutation effect on protein dysfunction[25]. Therefore, we created an MMS that incorporated four broad categories of 1) stability effects, 2) conservation of mutated amino acid, 3) position of the mutated amino acid, and 4) change in amino acid properties caused by substitution (Fig 5B-C). Specifically, we defined two stability parameters (stabilizing and destabilizing substitutions), two conservation parameters (conservation of residue and of amino acid substitution), eight location-based parameters (sheet, helix, c-terminus, active site, active region, N-terminus PLP pocket, H-protein interface, and dimerization interface mutations), and six parameters based on change in amino acid properties (change in polarity, charge, aromaticity, codon availability, size, and to/from proline). Together these contributed eighteen distinct parameters which, when applied to 251 NKH causing-missense mutations, yielded an MMS for each mutation (S2 Table). Phi correlation analysis of the eighteen parameters demonstrated independence between all parameters. There were light negative correlations between conservation of amino acid substitution and change in polarity (φ = −0.44, with 1 being perfect correlation) and change in volume (φ = −0.51) (Fig 5B). However, these correlations are statistically weak and hence both parameters were retained. The distribution of the scores indicates a lack of distribution bias in scoring (Fig 5D). Only 4 (R212K, R377Q, E495Q, and N709S) of 251 mutations received a score of 0, confirming MMS yields value for the vast majority of known mutations. Mutations with scores of 1-2 were considered mild (N = 58), 3-4, moderate (N = 93), and ≥5 severe (N=99) (S4 Table), consistent with reports that NKH disease in the patient population is more likely to be moderate to severe (rather than mild) [12, 13].

### Clinical Outcomes Score to Assess Patient Disease Severity Status

NKH is a multi-system neuro-metabolic disorder. Hennermann et al [12] have classified disease on the basis of presence and absence of neurological/brain features, while Swanson et al [13] classified disease based on reaching developmental outcomes. But the lack of a disease severity scale based on a quantitative, dynamic progression of different NKH symptoms across severe and attenuated disease, has limited linking genotype to phenotype.

To develop such a clinical severity scale, we began by building a comprehensive list of symptoms associated with NKH. Prior studies utilized a list of 12 NKH symptoms [12]. We reviewed 131 patient records from 26 publications over the last 15 years, to identify fifty-eight unique symptoms (Table 2), that were identified and classified into eleven categories of hyperglycinemia, cognitive disorders, seizures, muscle/movement control, brain malformations/injury, respiration, hormonal disorders, hearing, eyesight, immune system and digestion. A category was considered a major disease domain if it was represented in at least of 30% of patients with recorded symptoms, with the exception of glycine elevation, because while it provides a diagnostic criterion, it does not correlate well to the severity of neurobehavioral disease[13]. As shown in Table 3, four major domains emerged, namely (and in order of frequency) cognitive disorders (81%), seizures (73%), muscle and movement dysfunctions (35%), and brain malformations (32%). They encompassed 46 of 58 (79% of) symptoms. Respiratory defects were seen in 17% of patients, which is likely a result of under-reporting of the symptom. Regardless, respiratory issues usually self-resolve (barring when intubation was removed because of the overall poor prognosis), and respiration was not included in symptomatic domains. Hearing, eyesight, immune system, hormonal and digestive disorders were each seen in less than 3% of cases and therefore not included. A Likert-like scale was used to assign major domain scores of 0-3 based on severity in each domain (Table 3). Cognitive disorders and muscle/movement control were assigned linearly from 0-3. The seizure domain was assigned a non-linear step increase of 1 to 3 corresponding to transition from controlled seizure activity to uncontrolled seizure activity (and capture the severity of intractable seizure compared to controlled seizure). The brain malformation domain was assigned a binary choice of 0 or 3, because any brain malformation is expected to seriously impact neurological disease. Summation of all four domains yielded a patient clinical outcome score (COS), with a maximal score of 12.

**Table 2.**
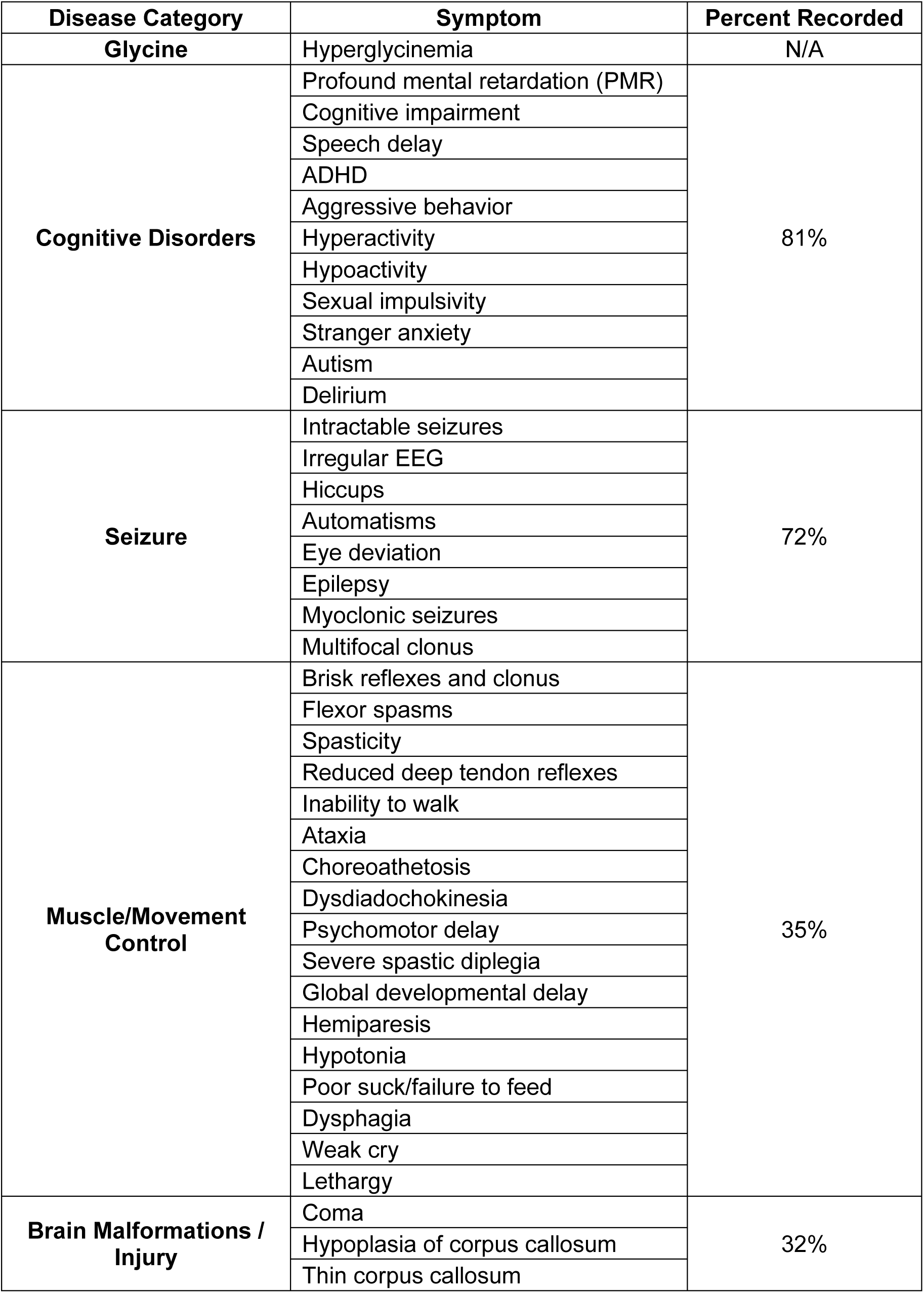
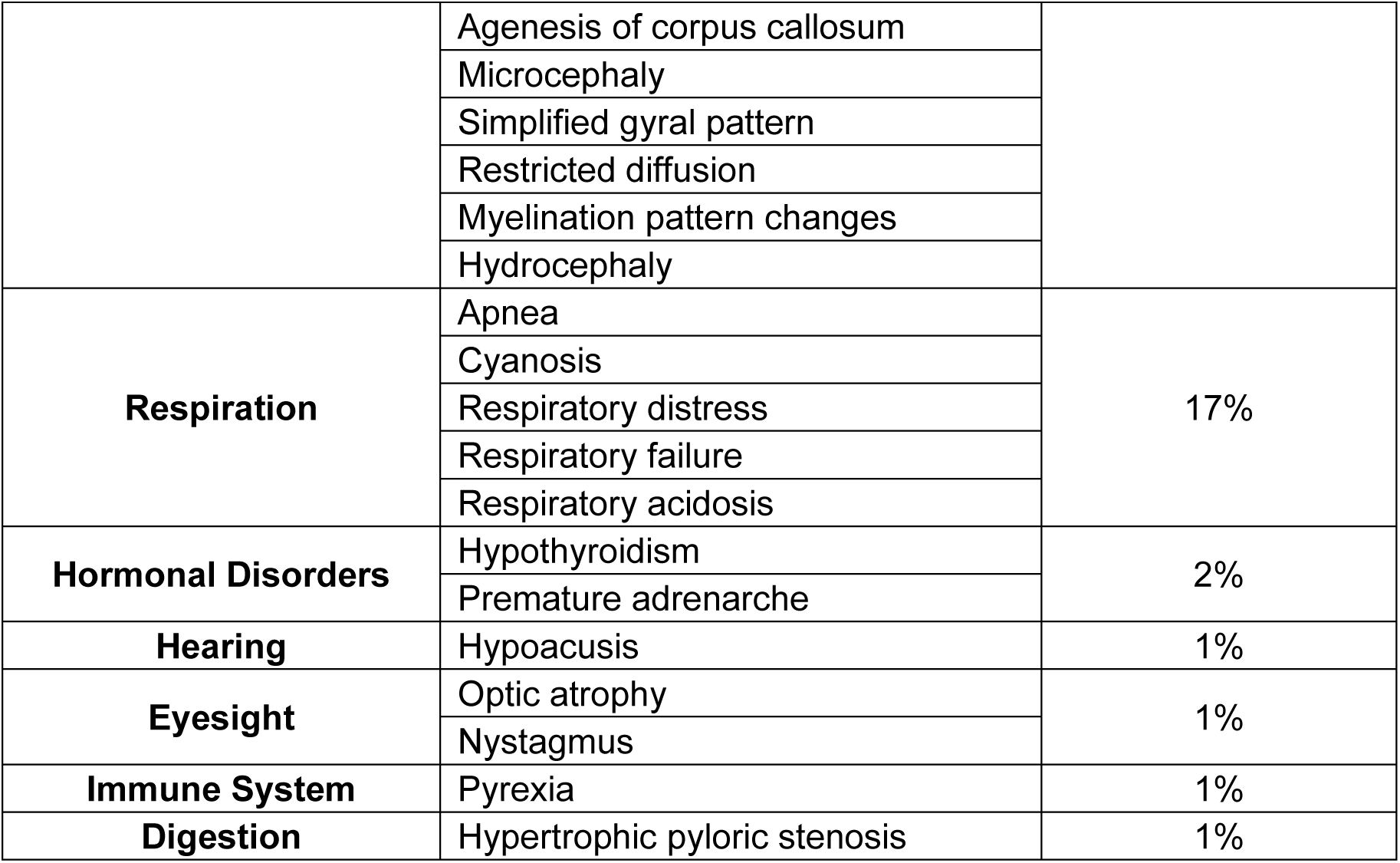
NKH Symptoms and Disease Categories.

**Table 3.** Quantitative severity scale for major NKH disease domains.

| **Domain and Associated Scoring Scale** |
| --- |
| **Cognitive Disorders** |
| 0 – No disease |
| 1 – Behavioral issues, learning disabilities, speech delay |
| 2 – Mental disability, some words, global delay in developmental markers |
| 3 – Severe mental disability, no cognitive abilities |
| **Seizures** |
| 0 – No disease |
| 1 – Hiccups (infants), Abnormal EEG, Seizures controlled by medication |
| 3 – Intractable seizures (> 2 AEDs) |
| **Muscle/Movement Control** |
| 0 – No disease |
| 1 – Assisted locomotion |
| 2 – Hypotonia, able to roll over or lift head, low muscle tone |
| 3 – Severe hypotonia, unable to roll over or lift head or severe global developmental delay |
| **Brain malformation** |
| 0 – No disease |
| 3 – Present |

EEG = electroencephalogram
AEDs = Anti-epileptic drugs

In the assessment of patient COS, we removed 45 records where patients had died, because death can occur due to a single acute event that may not reflect multi-symptom disease severity. An additional 12 patient records were excluded because they contained information in only 1 (of the 4) major domains. The remaining 74 patient records were quantitatively scored for their associated major disease domains (Tables 4-5, Table S2). The majority of records from patients were scored for 3 out of 4 major disease domains in both homozygous and compound heterozygotes (Table 4-5). Seizures and cognitive disorders were the two most common disease domains, although muscle/movement control and brain malformation were also often reported.

**Table 4.**
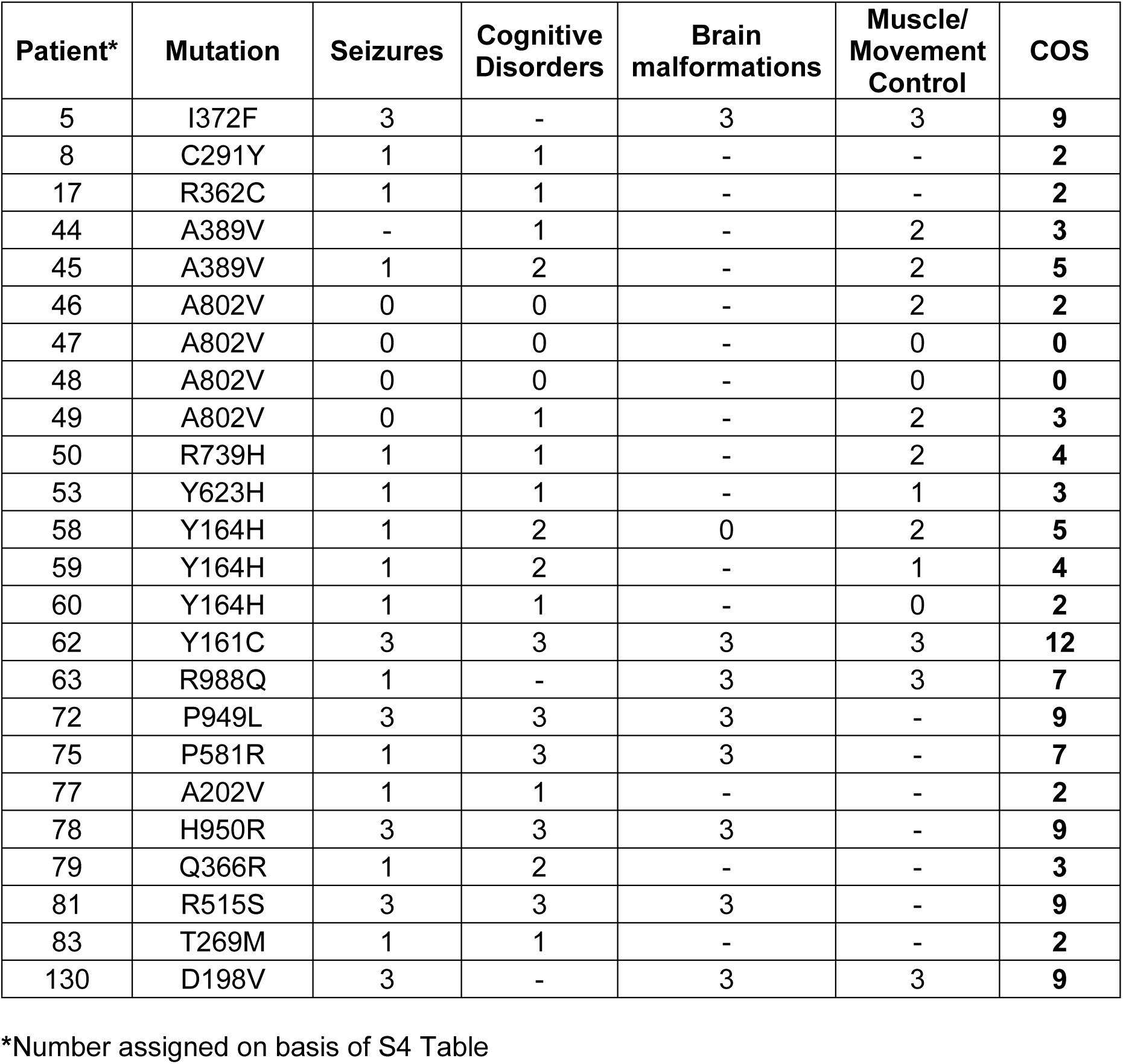
COS for patients with homozygous mutations.

| **Patient\*** | **Mutation** | **Seizures** | **Cognitive Disorders** | **Brain malformations** | **Muscle/ Movement Control** | **COS** |
| --- | --- | --- | --- | --- | --- | --- |
| 5 | I372F | 3 | - | 3 | 3 | **9** |
| 8 | C291Y | 1 | 1 | - | - | **2** |
| 17 | R362C | 1 | 1 | - | - | **2** |
| 44 | A389V | - | 1 | - | 2 | **3** |
| 45 | A389V | 1 | 2 | - | 2 | **5** |
| 46 | A802V | 0 | 0 | - | 2 | **2** |
| 47 | A802V | 0 | 0 | - | 0 | **0** |
| 48 | A802V | 0 | 0 | - | 0 | **0** |
| 49 | A802V | 0 | 1 | - | 2 | **3** |
| 50 | R739H | 1 | 1 | - | 2 | **4** |
| 53 | Y623H | 1 | 1 | - | 1 | **3** |
| 58 | Y164H | 1 | 2 | 0 | 2 | **5** |
| 59 | Y164H | 1 | 2 | - | 1 | **4** |
| 60 | Y164H | 1 | 1 | - | 0 | **2** |
| 62 | Y161C | 3 | 3 | 3 | 3 | **12** |
| 63 | R988Q | 1 | - | 3 | 3 | **7** |
| 72 | P949L | 3 | 3 | 3 | - | **9** |
| 75 | P581R | 1 | 3 | 3 | - | **7** |
| 77 | A202V | 1 | 1 | - | - | **2** |
| 78 | H950R | 3 | 3 | 3 | - | **9** |
| 79 | Q366R | 1 | 2 | - | - | **3** |
| 81 | R515S | 3 | 3 | 3 | - | **9** |
| 83 | T269M | 1 | 1 | - | - | **2** |
| 130 | D198V | 3 | - | 3 | 3 | **9** |

\*Number assigned on basis of S4 Table

**Table 5.** COS for patients with heterozygous mutations.

| **Patient\*** | **Mutation 1** | **Mutation 2** | **Seizures** | **Cognitive Disorders** | **Brain malformation** | **Muscle/ Movement Control** | **COS** |
| --- | --- | --- | --- | --- | --- | --- | --- |
| 1 | P509A | E597K | 1 | - | 3 | 2 | 6 |
| 3 | D295Y | R536Q | 3 | - | - | 3 | 6 |
| 9 | A202V | IVS22+1G>C | 1 | 1 | - | - | 2 |
| 10 | A202V | IVS22+1G>C | 1 | 1 | - | - | 2 |
| 11 | A802V | IVS22+1G>A | 1 | 1 | - | - | 2 |
| 12 | A802V | IVS22+1G>A | 1 | 1 | - | - | 2 |
| 13 | A389V | IVS12+2T>G | 1 | 1 | - | - | 2 |
| 14 | A389V | IVS12+2T>G | 1 | 1 | - | - | 2 |
| 26 | P267A | K376E | 1 | 2 | - | - | 3 |
| 27 | R373W | intronic | 1 | 1 | - | - | 2 |
| 28 | R373W | M1I | 1 | 1 | - | - | 2 |
| 31 | H371D | del GLDC | 1 | - | 3 | 3 | 7 |
| 51 | N150T | R790W | 1 | 3 | 0 | 2 | 6 |
| 52 | L82W | 607fs | 1 | 1 | 0 | 2 | 4 |
| 54 | C1002W | S419X | 3 | 2 | - | - | 5 |
| 55 | C1002W | S419X | - | 2 | - | 2 | 4 |
| 56 | Q620R | del Exon 3-9 | 3 | - | 0 | 3 | 6 |
| 57 | S132L | S86Vfs_119 | 3 | - | 3 | 2 | 8 |
| 68 | T894A | del Exon3 | 3 | 3 | - | 2 | 8 |
| 69 | Y839C | intronic | 3 | - | 3 | 3 | 9 |
| 73 | G771R | M552V | 1 | - | - | 2 | 3 |
| 74 | R739H | del Exon 1-2 | 1 | 1 | 0 | 1 | 3 |
| 76 | R461Q | del GLDC | 1 | 2 | 3 | 2 | 8 |
| 85 | G761R | Y632X | 3 | 3 | 3 | - | 9 |
| 86 | R515S | IVS19-1G>A | 1 | 3 | 3 | - | 7 |
| 87 | S132L | E167X | 3 | 3 | 3 | - | 9 |
| 89 | P907L | del Exon 1-24 | - | 3 | 3 | - | 6 |
| 90 | R515S | G618R | - | 3 | 3 | - | 6 |
| 91 | A733V | IVS19-1G>A | 1 | 3 | - | - | 4 |
| 95 | R515S | IVS19-1G>A | - | 3 | 3 | - | 6 |
| 97 | L885P | Y637X | - | 3 | 3 | - | 6 |
| 98 | G771R | IVS19-1G>A | - | 3 | 3 | - | 6 |
| 100 | F334L | del GLDC | - | 3 | 3 | - | 6 |
| 101 | A389V | R515S | - | 3 | 3 | - | 6 |
| 102 | L548V | del GLDC | 1 | 2 | - | - | 3 |
| 103 | A802V | IVS22+1G>C | 1 | 2 | - | - | 3 |
| 104 | A802V | R515S | 1 | 2 | - | - | 3 |
| 105 | A389V | R515S | 3 | 1 | - | - | 4 |
| 106 | I381T | R461Q | 1 | 1 | - | - | 2 |
| 107 | A283P | R461Q | 3 | 1 | - | - | **4** |
| 108 | A283P | R461Q | 1 | 1 | - | - | **2** |
| 109 | R461Q | IVS12+2T>G | 1 | 1 | - | - | **2** |
| 110 | Y161C | R347S | 1 | 1 | - | - | **2** |
| 111 | G156R | G728E | 1 | 1 | - | - | **2** |
| 112 | R630P | L548V | 1 | 1 | - | - | **2** |
| 113 | A802E | IVS19+2T>G | 1 | 1 | - | - | **2** |
| 114 | V905G | G728E | 1 | 1 | - | - | **2** |
| 115 | G652E | R373Q | 1 | 1 | - | - | **2** |
| 129 | L885P | W897C | 3 | 3 | 3 | 3 | **12** |
| 131 | A377V | A694Dfs | 3 | 3 | 0 | 2 | **8** |

\*Number assigned on basis of S4 Table

As shown in Fig 6A, the majority of patient COS’s ranged from 2-9, with two patients showing the maximal scores of 12. Severe patients scored above 5 (having a severe score of 3 in at least one domain and moderate score of 2 in another). In our cohort, there were 29 patients with severe disease. 43 patient records showed attenuated disease, and two individuals were asymptomatic (despite being homozygous for the pathogenic A802V mutation; Fig 6A). Of the 74 patients, 40 were male and 34 were female, and both genders showed similar range distributions of severe and attenuated COS’s (Fig 6B). Analyses for age suggest that severe disease was more prominent in children < 5 in both genders (Fig 6C).

**Figure 6.**
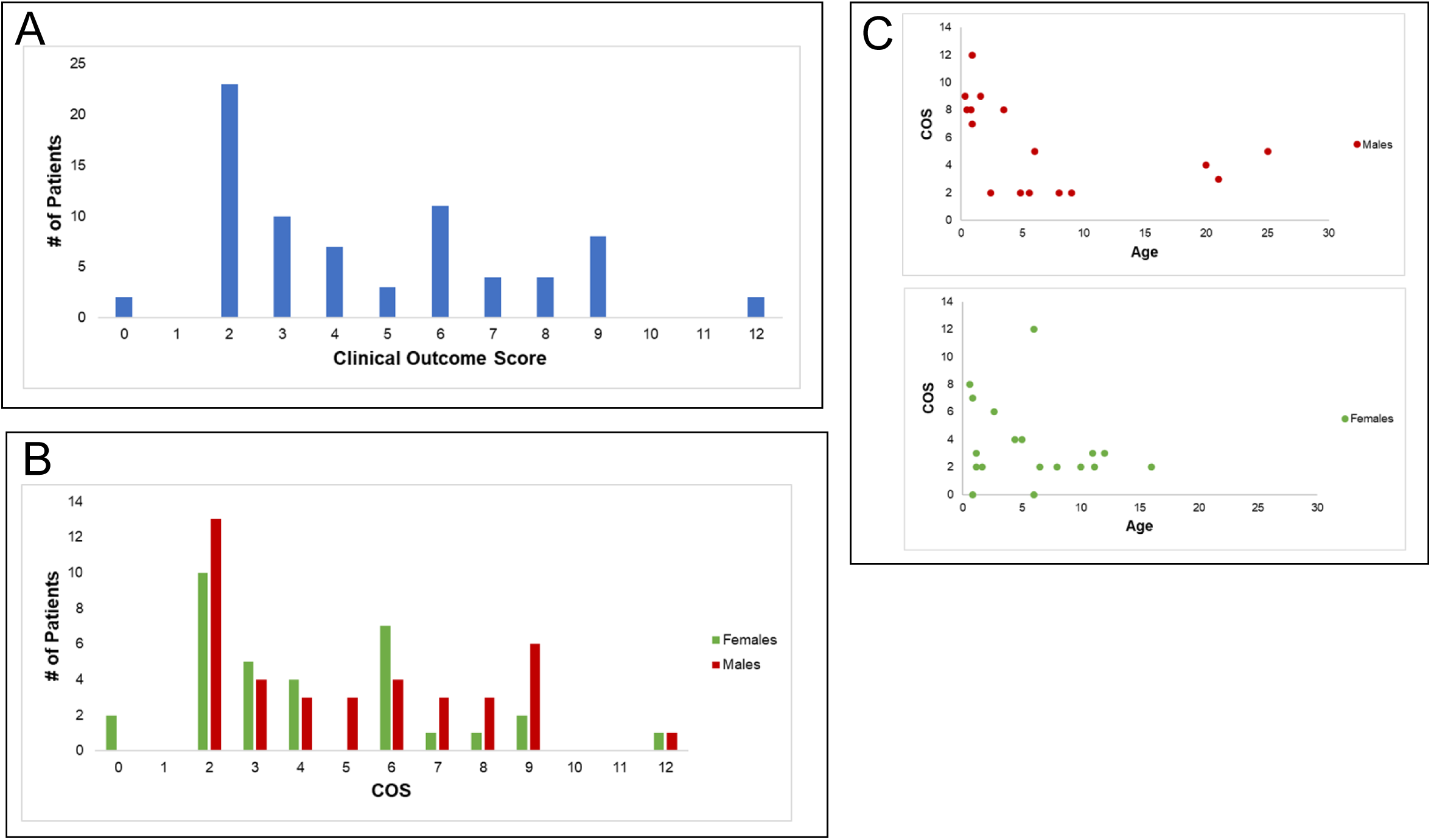
Clinical outcome scores based on symptom analyses of patient case reports. [A] Distribution of Clinical Outcomes Scores as a measure of disease severity in 74 patients based on quantitative determination of major symptomatic domains of seizures, muscle/movement control, cognition, and brain malformations as presented in Tables 4-5. The bulk of patient scores ranged from 2-9 although two patients showed scores of 12. Scores above five were designated severe, yielding 29 patients with severe disease, 43 with attenuated disease, and 2 who were asymptomatic. [B] Gender distribution of COS Scores. Of the 74 patients, 40 were male and 34 were female. Male and female patients had similar distributions of severe and attenuated scores, suggesting COS reveals no gender preference. [C] Age distribution. Examination of COS scores as a function of age for both males and females suggests that severe disease is dominant in patients < 5 years old.

We examined whether COS could be corroborated with known information about patient mutations for both homozygous and compound heterozygous mutations. First, for the 24 patients with homozygous mutations, each allele was assigned the same COS (see Table 4), which also served as the overall patient COS. When multiple patients were homozygous for the same mutation, an average COS was calculated. As shown in Table 6, four of the 18 homozygous mutations (R515S, T269M, A389V, A802V) are amongst the top 10 mutations found in clinical NKH. R515S received a COS of 9 (out of 12) consistent with its association with severe disease[26]. T269M, A389V, A802V, received COS’s of lower than 5, in keeping with their association with attenuated disease[27–29]. Overall, 14 out of 16 attenuated (COS ≤ 5) cases were associated with low MMS, while 4 of 8 severe cases (COS greater than 5) were associated with MMS ≥ 5. There was no overt positional bias of COS in GLDC-protein (Fig 7A).

**Figure 7.**
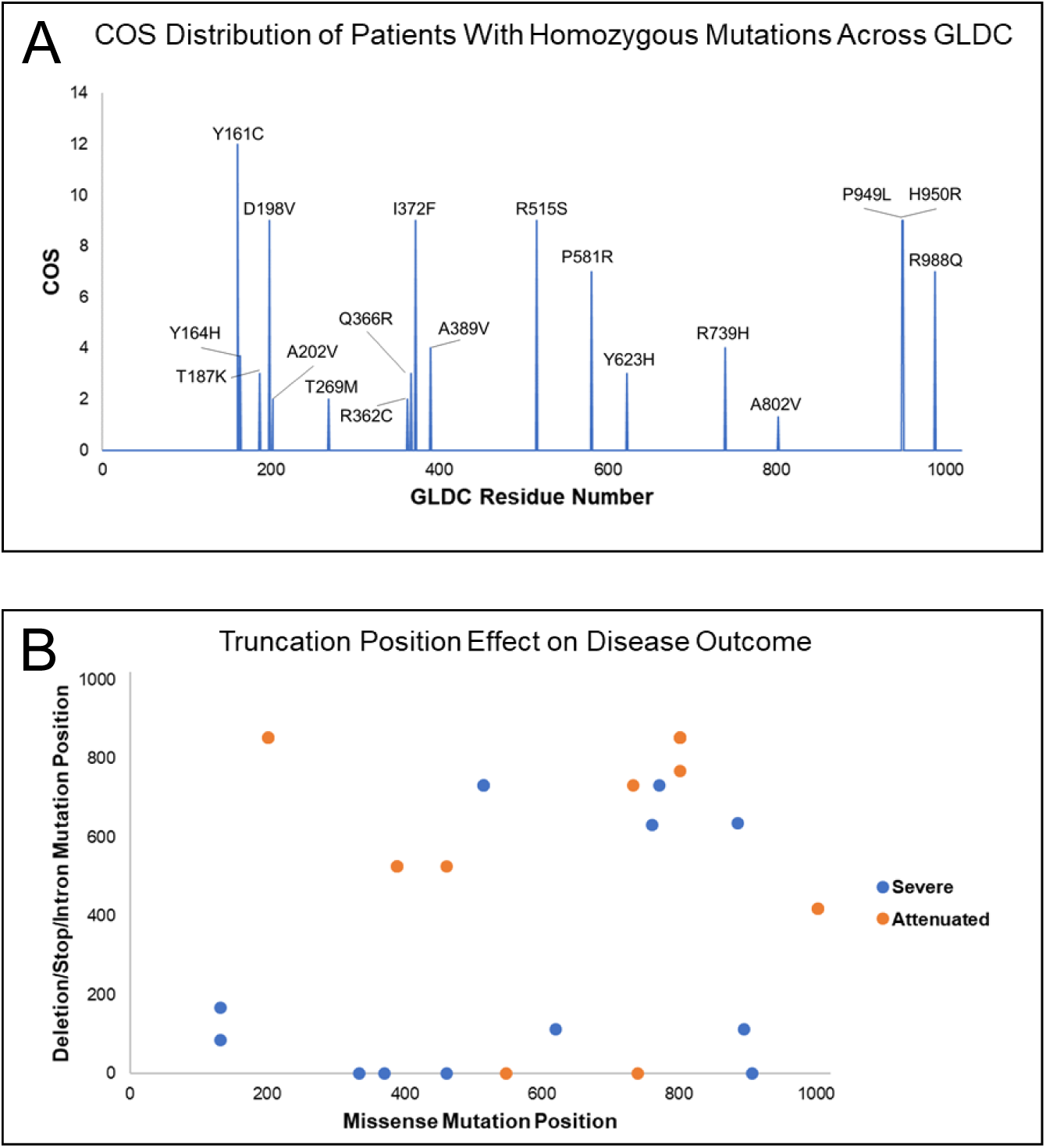
Distribution of Mutation and COS for homozygote and heterozygote patients. [A] Distribution of COS vs mutation position in GLDC-protein of homozygote patients shows no overt positional bias in COS. [B]. Positional distribution of deletion/truncation vs missense mutations in heterozygote patients is shown. Attenuated patientsare in orange; Severe patients in blue.

**Table 6.**
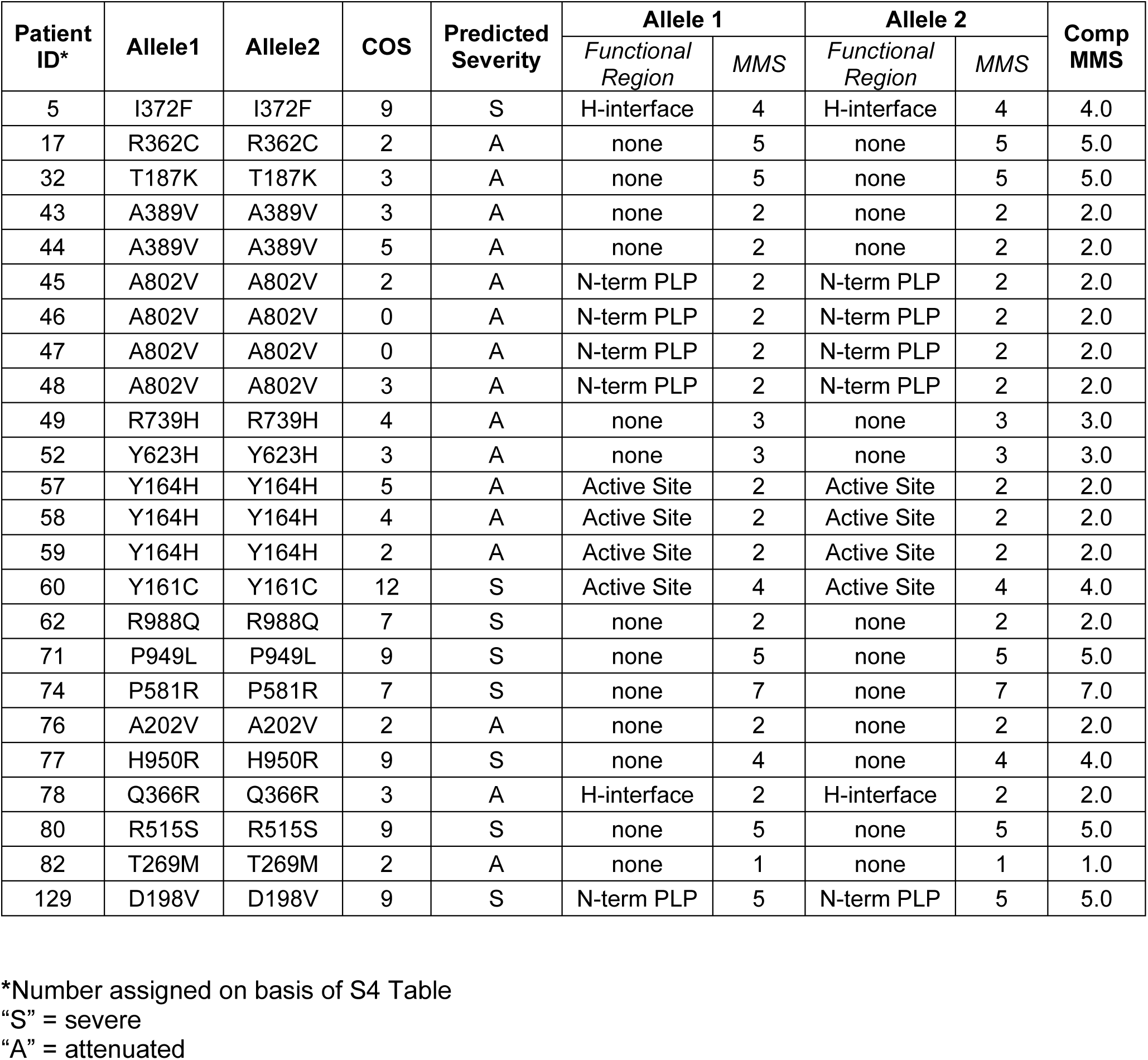
MMS for patients with homozygous mutations.

For compound heterozygotes, since this group had two alleles, the observed patient COS was considered to be the result of one half of the contribution of each allele. In cases where the second allele was a deletion or truncation, it was assigned an MMS of 5 (corresponding to the threshold of high pathogenicity index of MMS, see Fig 5D). As shown in Table 7, 26 of 29 (89.6%) of patients with attenuated disease (COS<5) showed lower MMS scores (<5). 8 of 21 (38%) patients with severe disease (COS>5) showed higher MMS scores (≥5). Notably of 11 patients with truncations/deletions and COS>5, nine showed MMS scores below 5. Examination of the location of missense mutation and deletions/truncations in each heterozygote suggested that deletions or truncations early in the gene maybe associated with severe disease (Fig 7B).

**Table 7.** MMS of patients with heterozygous mutations.

| Patient ID | Allele1 | Allele2 | COS | Predicted Severity | Allele 1 |  | Allele 2 |  | Comp MMS |
| --- | --- | --- | --- | --- | --- | --- | --- | --- | --- |
|  |  |  |  |  | Functional Region | MMS | Functional Region | MMS |  |
| 1 | P509A | E597K | 6 | S | none | 2 | none | 2 | 2.0 |
| 3 | D295Y | R536Q | 6 | S | none | 5 | dimerization | 4 | 4.5 |
| 26 | P267A | K376E | 3 | A | N PLP | 4 | H-interface | 2 | 3.0 |
| 51 | N150T | R790W | 6 | S | dimerization | 4 | none | 5 | 4.5 |
| 73 | G771R | M552V | 3 | A | none | 6 | dimerization | 3 | 4.5 |
| 90 | R515S | G618R | 6 | S | none | 5 | active site | 7 | 6.0 |
| 101 | A389V | R515S | 6 | S | none | 2 | none | 5 | 3.5 |
| 104 | A802V | R515S | 3 | A | N PLP | 2 | none | 5 | 3.5 |
| 105 | A389V | R515S | 4 | A | none | 2 | none | 5 | 3.5 |
| 106 | I381T | R461Q | 2 | A | none | 5 | none | 2 | 3.5 |
| 107 | A283P | R461Q | 4 | A | none | 2 | none | 2 | 2.0 |
| 108 | A283P | R461Q | 2 | A | none | 2 | none | 2 | 2.0 |
| 110 | Y161C | R347S | 2 | A | active site | 4 | none | 3 | 3.5 |
| 111 | G156R | G728E | 2 | A | none | 5 | none | 7 | 6.0 |
| 112 | R630P | L548V | 2 | A | none | 6 | dimerization | 3 | 4.5 |
| 114 | V905G | G728E | 2 | A | none | 4 | none | 7 | 5.5 |
| 115 | G652E | R373Q | 2 | A | none | 5 | H-interface | 2 | 3.5 |
| 129 | L885P | W897C | 12 | S | none | 6 | active site | 5 | 5.5 |
| 9 | A202V | IVS22+1G>C | 2 | A | none | 2 | N/A | 5 | 3.5 |
| 10 | A202V | IVS22+1G>C | 2 | A | none | 2 | N/A | 5 | 3.5 |
| 11 | A802V | IVS22+1G>A | 2 | A | N PLP | 2 | N/A | 5 | 3.5 |
| 12 | A802V | IVS22+1G>A | 2 | A | N PLP | 2 | N/A | 5 | 3.5 |
| 13 | A389V | IVS12+2T>G | 2 | A | none | 2 | N/A | 5 | 3.5 |
| 14 | A389V | IVS12+2T>G | 2 | A | none | 2 | N/A | 5 | 3.5 |
| 27 | R373W | splice site | 2 | A | H-interface | 5 | N/A | 5 | 5.0 |
| 69 | Y839C | splice site | 9 | S | none | 5 | N/A | 5 | 5.0 |
| 86 | R515S | IVS19-1G>A | 7 | S | none | 5 | N/A | 5 | 5.0 |
| 91 | A733V | IVS19-1G>A | 4 | A | none | 4 | N/A | 5 | 4.5 |
| 95 | R515S | IVS19-1G>A | 6 | S | none | 5 | N/A | 5 | 5.0 |
| 98 | G771R | IVS19-1G>A | 6 | S | none | 6 | N/A | 5 | 5.5 |
| 103 | A802V | IVS22+1G>C | 3 | A | N PLP | 2 | N/A | 5 | 3.5 |
| 109 | R461Q | IVS12+2T>G | 2 | A | none | 2 | N/A | 5 | 3.5 |
| 113 | A802E | IVS19+2T>G | 2 | A | N PLP | 4 | N/A | 5 | 4.5 |
| 31 | H371D | del GLDC | 7 | S | none | 3 | N/A | 5 | 4.0 |
| 56 | Q620R | del Exon 3-9 | 6 | S | none | 3 | N/A | 5 | 4.0 |
| 68 | T894A | del Exon 3 | 8 | S | active site | 4 | N/A | 5 | 4.5 |
| 74 | R739H | del Exon 1-2 | 3 | A | none | 3 | N/A | 5 | 4.0 |
| 76 | R461Q | del GLDC | 8 | S | none | 2 | N/A | 5 | 3.5 |
| 89 | P907L | del Exon 1-24 | 6 | S | none | 4 | N/A | 5 | 4.5 |
| 100 | F334L | del GLDC | **6** | S | none | 2 | N/A | 5 | 3.5 |
| 102 | L548V | del GLDC | **3** | A | dimerization | 3 | N/A | 5 | 4.0 |
| 52 | L82W | 607fs | **4** | A | none | 4 | N/A | 5 | 4.5 |
| 54 | C1002W | S419X | **5** | A | none | 4 | N/A | 5 | 4.5 |
| 55 | C1002W | S419X | **4** | A | none | 4 | N/A | 5 | 4.5 |
| 57 | S132L | S86Vfs_119 | **8** | S | none | 4 | N/A | 5 | 4.5 |
| 85 | G761R | Y632X | **9** | S | none | 6 | N/A | 5 | 5.5 |
| 87 | S132L | E167X | **9** | S | none | 4 | N/A | 5 | 4.5 |
| 97 | L885P | Y637X | **6** | S | none | 6 | N/A | 5 | 5.5 |
| 131 | A377V | A694Dfs | **8** | S | none | 1 | N/A | 5 | 3.0 |
| 28 | R373W | M1I | **2** | A | H-interface | 5 | mito leader<br>seq | 5 | 5.0 |

\*Number assigned on basis of S4 Table
“S” = severe
“A” = attenuated

### Weighted-Optimization of the MMS by the COS

For both homozygous and compound heterozygous mutations in a patient, the MMS is a better predictor of attenuated versus severe disease. This suggested that the MMS needed further optimization against clinical disease to more accurately capture the determinants of severe disease. It should also be noted that the MMS does not incorporate human factors such as genetic background which is known to play a prominent role in disease manifestation in genetic disorders.

#### i. Homozygous Mutations

The MMS was first applied to all 18 homozygous mutations associated with 24 clinical cases (summarized in Fig 8A). As shown in Table 6, every patient mutation was assigned an MMS score in addition to the previously determined COS. Ten variants that were found in homozygous form of GLDC in the Exome Aggregation Consortium (ExAC) database, hosted by the Broad Institute, (Table 8) were also scored. The ExAC database utilizes genomic data from healthy individuals; thus, homozygosity of these mutations indicates that they are non-pathogenic. They were included therefore as a non-pathogenic control group and were assigned a COS of zero.

**Figure 8.**
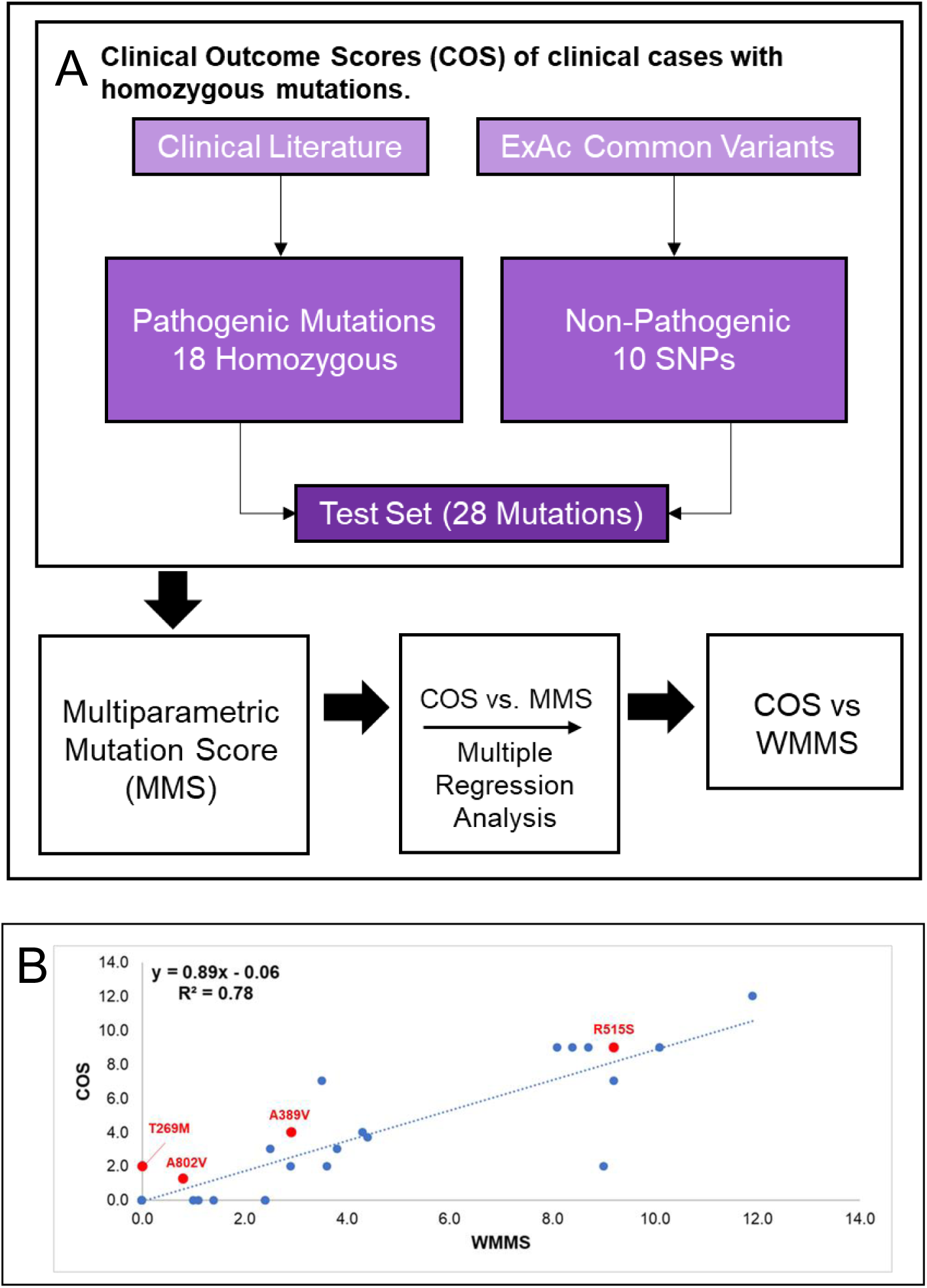
Weight Multiparametric Score (WMMS) for Homozygous Mutations. [A] A flowchart of the generation of weighted multiparametric scores (WMMS’s) for homozygous mutations. Data were assessed from twenty-four NKH patients with homozygous mutations and at least two scoreable major symptomatic domains. Control data was from ten variants that were found in homozygous form of GLDC in the Exome Aggregation Consortium (ExAC) database (Broad Institute) and assigned COS of 0. [B] MMS parameters were weighted to maximize the correlation R^2^-value and yield the COS vs WMMS plot shown for a test set of 28 mutations (18 pathogenic, 10 nonpathogenic from the ExAC database).

**Table 8.** ExAc Nonpathogenic Mutations

| **MUTATION** | **# of ExAc Homozygotes** | **COS** | **MMS** |
| --- | --- | --- | --- |
| M107V | 80 | 0.0 | 0.0 |
| R224H | 22 | 0.0 | 2.0 |
| C291G | 6 | 0.0 | 2.0 |
| R410K | 8 | 0.0 | 0.0 |
| L462V | 122 | 0.0 | 0.0 |
| E503A | 14 | 0.0 | 2.0 |
| A569T | 4 | 0.0 | 2.0 |
| V705M | 1 | 0.0 | 1.0 |
| V735L | 8 | 0.0 | 2.0 |
| A794T | 3 | 0.0 | 1.0 |

COS = Clinical Outcome Score
MMS = Multiparametric Mutation Score

The COS was applied as a function of the MMS of homozygous mutations (S3 Fig, Fig 8A), and this correlation was used to optimize parameter weights in order to yield a model with more biological and clinical value. Weighting was automated using Python. Each parameter weight was computationally allowed to range from 0.0 to 5.0 (or −5.0 to 0.0 for parameter 4) and the combination of parameter weights which yielded the highest R^2^-value was selected (see Methods and Table 8). The final correlation between COS and the weighted MMS (WMMS) yielded an R^2^-value of 0.78 (Fig 8B). The resulting WMMS range was slightly higher than MMS, but as shown in Fig. 8B, R515S, the severe and dominant NKH mutation, remained predicted to be pathogenic with a WMMS of 9.2. Less pathogenic but prevalent mutations A389V (WMMS = 2.9), and A802V (WMMS = 0.8) and T269M (WMMS = 0.0) are predicted to be attenuated.

R515S scored in five parameters. R515 is a highly evolutionary conserved, C-terminal residue, and the substitution to serine is destabilizing and causes a change in charge and volume (see S5 Table, S6 Table). A389V and A802V each scored in 4 parameters, including the negatively scoring conserved evolutionary substitution parameter. A389 is a conserved residue in an α-helix, while A802 is a C-terminal residue and part of a C-terminal loop that is within 5 Å of the predicted N-term PLP pocket. Substitution from alanine to valine causes a change in volume but is a well-tolerated substitution throughout evolution. T269M only scored in one parameter, as the mutation results in a change in polarity.

#### ii. Compound Heterozygous Mutations

The overall strategy for application of COS and MMS (S3 Fig) for compound heterozygous individuals is summarized in Fig 9A. Compound heterozygous scoring was complicated by the fact that each individual bears two different mutations, often with one of the mutations being a deletion, nonsense mutation, intronic mutation, or a mutation in the mitochondrial signal sequence. Of the 50 compound heterozygous patients in our cohort, 32 patients have one mutation that is not a missense mutation. MMS parameters were designed specifically for missense mutations; thus, to facilitate scoring of these 32 patients, deletions, nonsense mutations, intronic mutations, and mutations in mitochondrial signal sequence were each assigned scores of 5.0, corresponding to a moderate MMS score. We then calculated a composite score for each compound heterozygous patient according to the formula ½ Allele 1 Score + ½ Allele 2 Score, where the allele score for missense mutations was the MMS, and the score for other mutations was fixed at 5.0 (Table 7). We also scored variants from healthy individuals (obtained from dbGaP), which were included as non-pathogenic controls with a COS of zero (Table 9). MMS parameters and deletion, nonsense, intronic, and mitochondrial signal sequence mutation scores were optimized using the best fit line of the COS vs the composite score. The R^2^ of resulting best fit of COS vs composite score fit obtained with heterozygous mutations (Fig 9B) was slightly lower than that observed with homozygous mutations (0.69 vs 0.78) (Fig 8B). We suggest that with more robust clinical data, these differences, in addition to differences in weighting (Table 10), between homozygous and heterozygous-trained parameters would converge. Nonetheless, our analyses informed that COS is proportional to WMMS.

**Figure 9.**
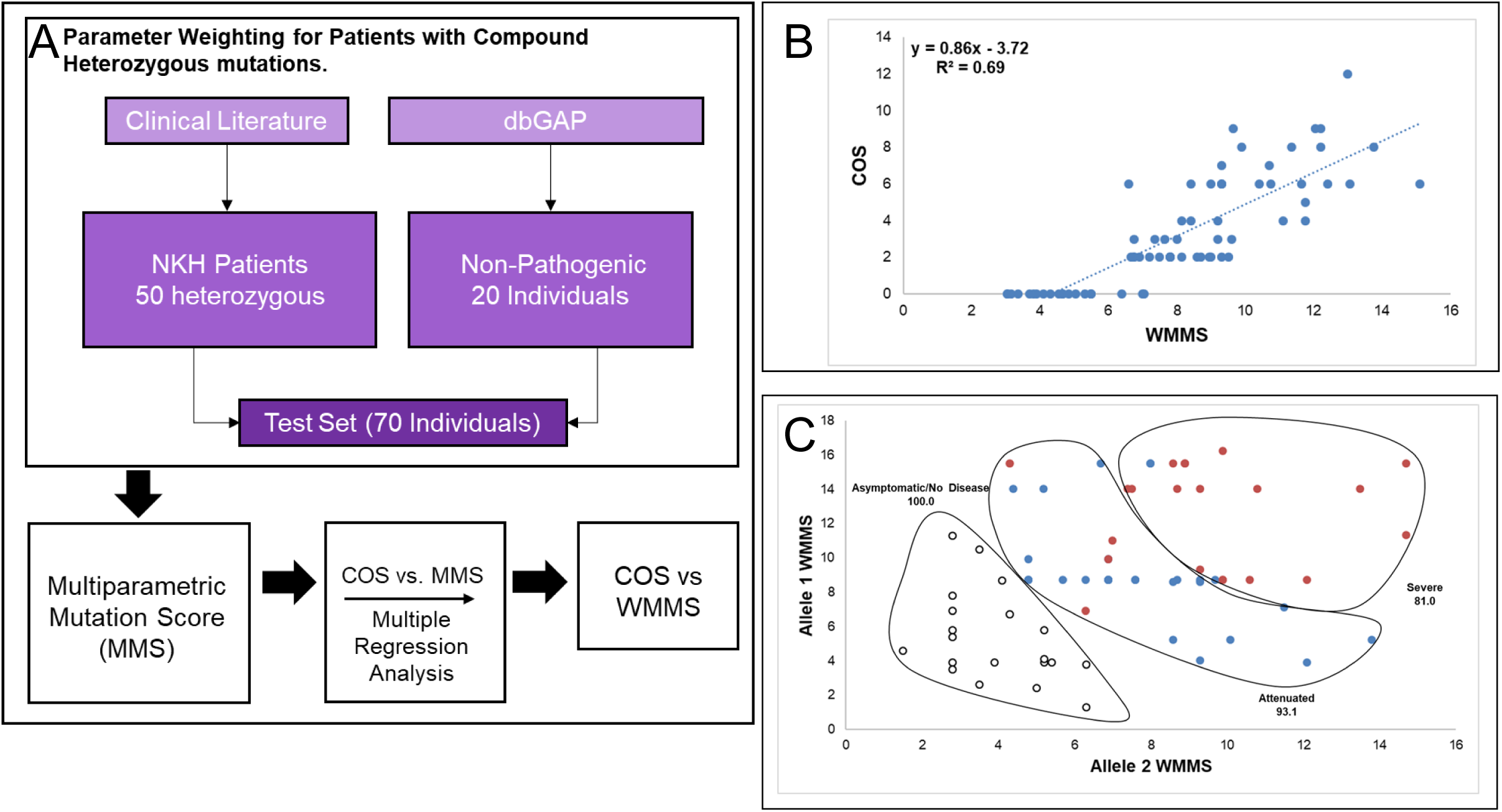
WMMS for Heterozygous Mutations. [A] A flowchart of the process used to generate WMMS’s for heterozygous mutations. Data were assessed from fifty NKH patients with compound heterozygous mutations and at least two scorable major symptomatic domains. Control data was from twenty healthy individuals with heterozygous variants obtained from the dbGaP database who were assigned COS of 0. [B] MMS parameters were weighted to maximize the correlation R^2^-value and yielded the COS vs WMMS plot shown. [C] Display of WMMS of Allele 1 versus Allele 2 for heterozygous patients yields characteristic of three different disease states namely asymptomatic/no disease (COS = 0; open dots), attenuated disease (COS = 1 – 5; blue dots), and severe disease (COS > 5; red dots) patients. Gating captured 100%, 93.1% and 81.0% of these respective populations, supporting the claim that WMMS scoring can clearly separate the different disease states.

**Table 9.** dbGaP Healthy Individuals’ Mutations and MMS’s

| **Allele 1** | **Allele 2** | **COS** | **Allele 1 MMS** | **Allele 2 MMS** | **Individual MMS** |
| --- | --- | --- | --- | --- | --- |
| E503A | V233A | 0 | 2.0 | 2.0 | 2.0 |
| E503A | T799S | 0 | 2.0 | 2.0 | 2.0 |
| E503A | R236Q | 0 | 2.0 | 2.0 | 2.0 |
| E503A | V705M | 0 | 2.0 | 1.0 | 1.5 |
| T799S | V747I | 0 | 2.0 | 4.0 | 3.0 |
| V800I | A414T | 0 | 2.0 | 2.0 | 2.0 |
| E503A | N533S | 0 | 2.0 | 3.0 | 2.5 |
| P509A | I301M | 0 | 2.0 | 0.0 | 1.0 |
| P509A | G137S | 0 | 2.0 | 2.0 | 2.0 |
| E503A | R596Q | 0 | 2.0 | 2.0 | 2.0 |
| E503A | S814F | 0 | 2.0 | 8.0 | 5.0 |
| V705M | A64S | 0 | 1.0 | 2.0 | 1.5 |
| A794T | L462V | 0 | 1.0 | 0.0 | 0.5 |
| L207V | N193S | 0 | 1.0 | 0.0 | 0.5 |
| E669K | N413Y | 0 | 1.0 | 4.0 | 2.5 |
| V735L | V233A | 0 | 2.0 | 2.0 | 2.0 |
| V735L | R66K | 0 | 2.0 | 2.0 | 2.0 |
| E278K | A64T | 0 | 1.0 | 2.0 | 1.5 |
| V735L | A569T | 0 | 2.0 | 3.0 | 2.5 |
| A569T | Mito Leader Sequence | 0 | 3.0 | 5.0 | 4.0 |

**Table 10.** Trained Parameter Weights. The eighteen parameters comprising the MMS were weighted twice, once on the homozygous individual dataset and once on the heterozygous individual dataset. There are two stability parameters (stabilizing, destabilizing substitutions), two conservation parameters (conservation of residue and amino acid substitution), seven location-based parameters (sheet, helix, c-terminus, active site, active region, N-terminus PLP, H-protein interface, and dimerization interface), and six parameters based on change in amino acid properties (change in polarity, charge, aromaticity, codon availability, size, and to/from proline). Parameters, except conservation of substitution, were allowed to range from 0 to 5.0. Conservation of substitution was allowed to range from −5.0 to 0.

| Parameters | Unweighted | Homozygous Trained | Heterozygous Trained |
| --- | --- | --- | --- |
| Stability (Stabilizing) | 1.0 | 0.0 | 0.0 |
| Stability (Destabilizing) | 1.0 | 0.0 | 1.3 |
| Conservation of Residue | 1.0 | 4.1 | 2.4 |
| Conservation of Substitution | -1.0 | -5.0 | 0.0 |
| Helix | 1.0 | 3.1 | 2.6 |
| Sheet | 1.0 | 0.0 | 4.6 |
| C-term | 1.0 | 3.3 | 1.5 |
| Active Site | 1.0 | 4.2 | 5.0 |
| Active Region | 1.0 | 0.0 | 0.0 |
| N-term PLP | 1.0 | 0.0 | 1.4 |
| H-interface | 1.0 | 3.6 | 0.0 |
| Dimerization Interface | 1.0 | 0.0 | 0.4 |
| $\Delta$ Polarity | 1.0 | 0.0 | 0.0 |
| $\Delta$ Charge | 1.0 | 1.1 | 2.8 |
| $\Delta$ Aromaticity | 1.0 | 2.9 | 2.2 |
| $\Delta$ Proline | 1.0 | 0.0 | 5.0 |
| $\Delta$ Codon availability | 1.0 | 3.1 | 0.7 |
| $\Delta$ Size | 1.0 | 0.7 | 1.6 |

We further examined cases where both mutations were heterozygous by plotting the score of one allele as a function of the score of the second allele (Fig 9C). Individuals that were asymptomatic (COS = 0), attenuated (COS = 1 – 5), and severe (COS > 5) could be separated based on gating these populations (Fig 9C). Overall, this plot shows that severe disease requires two severe mutations, while attenuated disease is often either a mixture of mild and severe or two moderate mutations. Based on the asymptomatic cohort clustering, healthy individuals can be compound heterozygous for *GLDC* variants if one of them is very mild. These results support the use of WMMS’s as a predictive tool for NKH outcome.

### Disease Prediction

As discussed earlier, NKH has been characterized as either severe or attenuated, but predicting disease state on the basis of mutation has been a challenge. Previous attempts at predicting severe vs attenuated outcomes for NKH mutations have been based solely on biochemical activity of recombinant mutant proteins and hindered by technical challenges and low throughput. Our data indicates that based on the relationship between COS and WMMS in Figures 8B and 9C, WMMS from both the homozygous and heterozygous datasets can separate severe and attenuated disease. To test this hypothesis, we determined the average WMMS value for asymptomatic individuals (COS = 0), attenuated NKH patients (COS = 1 – 5), and severe NKH patients (COS > 5). For both the homozygous-trained (Fig 10A) and heterozygous-trained (Fig 10B) WMMS’s, each disease category shows significant separation. Most importantly, severe disease can be significantly distinguished (by student’s t-test) from attenuated disease with a p-value of 3.5e-5 for homozygous patients and a p-value of 3.2e-6 for heterozygous patients. Average values in the heterozygous set are higher relative to the homozygous set, but this is a consequence of the heterozygous-trained parameters having higher values in general. These data support the use of WMMS’s as a predictive tool for the clinical outcome of NKH based on retrospective clinical analyses.

**Figure 10.**
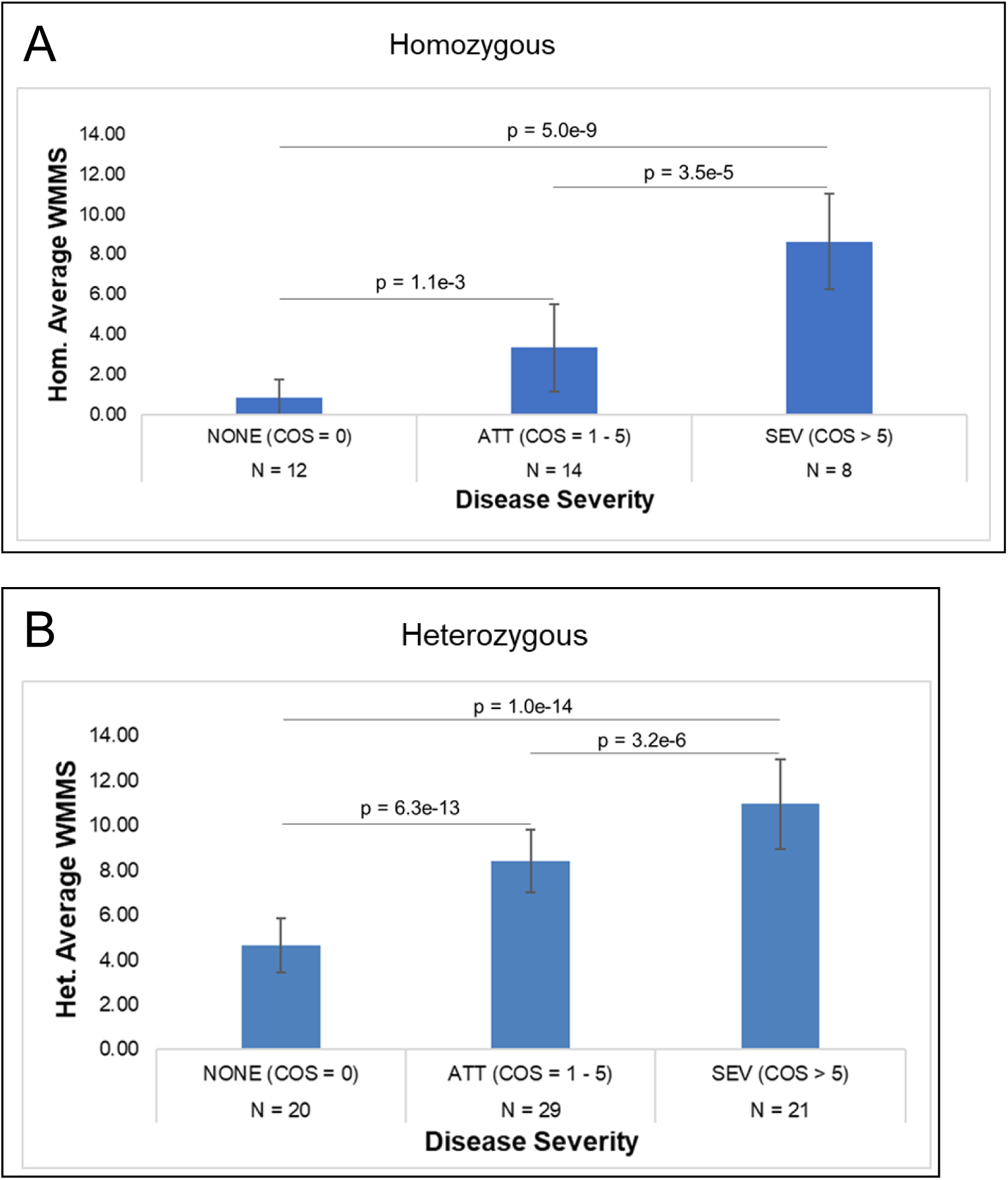
WMMS’s as a predictor of severe and attenuated disease. [A] Average homozygous-trained WMMS for homozygous patients by disease type. [B] Average heterozygous-trained WMMS’s for heterozygous NKH patients by disease type. The WMMS is a significant predictor of attenuated (ATT) and severe disease type (SEV) for disease caused by both homozygous and compound heterozygous missense mutations. Healthy controls with COS=0 are denoted as NONE.

## Discussion

We used computational approaches to undertake large scale, comparative and evolutionary analyses to enable multiparametric *in silico* assessment of 251 (of 255) NKH disease causing missense mutations based on structure-function properties intrinsic to GLDC protein. Further, our data ascribe either conserved, evolutionary function or clinical disease severity to 89 previously uncharacterized mutations. In contrast, prior studies have cumulatively reported on biochemical characterization of 49 mutations. Our evolutionary analyses support a new PLP binding function for the N-terminal PLP domain and predict residues that form the junction for H-GLDC interactions. Four of the top 10 clinical mutations (R515S, S564I G771R, and V905G) could only be annotated by the MMS.

The development and utilization of COS (rather than a biochemical activity) are particularly advantageous for a large gene like GLDC (1020 aa) with hundreds of disease-causing mutations. But as with many rare diseases, clinical symptom presentation of NKH disease is highly heterogeneous. We therefore based our clinical outcomes scale/score on multiple (∼50) symptoms that were, aggregated into four major disease domains. A major limitation of retrospective analyses of clinical records, is the variability in the published data content. This variability was somewhat decreased by excluding a minority of records that only contained one (of four) major symptomatic domains. While the removal of patients who died risked skew toward attenuated disease, the cohort analyzed nonetheless captured a dynamic range of disease outcomes, including severe disease in young patients.

In studies with compound heterozygotes, each mutation conferred 50% value to the final multi-parametric score. This may underestimate the contribution of dominant mutations and further adjustments may be needed to optimize the multi-parametric mutation score of compound heterozygotes. Further optimization is also needed for other non-missense mutations, such as considering the position of a deletion or premature stop codon in relation to the protein active site. Nonetheless we were able to score 50 compound heterozygotes and 24 homozygotes (total of 74 patient records) based on a comprehensive survey of the clinical literature and in future work could be rapidly scaled to all mutations, missense or otherwise.

In conjunction with pathogenic destabilizing mutations in target proteins, multiple additional factors influence disease progression in all genetic diseases. A nonsense or deletion early in the coding region is likely to be very severe, but much milder if located in the C-terminus after the protein active site. Intronic mutations that are not at the splice site will likely have less impact while those at the splice site can have differential effects depending on location. Finally, the overall genetic background has a profound influence on the emergence and progression of disease. While these parameters cannot be incorporated into the first step of developing a mutation-based score, weighting the MMS against the COS enables influence of parameters of ‘clinical’ relevance into the weighted MMS (WMMS). The WMMS was particularly important in analyses of compound heterozygous mutations in cases where the second mutation is a deletion whose location can dramatically alter severity.

In conclusion, our data suggest that WMMS is sufficiently robust to distinguish between severe and attenuated disease based on optimization using retrospective analyses of patient records. We therefore suggest that it presents a powerful tool to initiate and refine future analyses in larger prospective studies with active recruitment/review of patients and their medical records to strengthen management and prediction of disease course.

## Methods

### Homology modeling of P- and H-proteins

Human homology models were generated for GLDC and H-proteins using the SWISS-model (Swiss Institute of Bioinformatics), which generates homology models as described previously[30–32]. Human GLDC was modeled using as the template the solved crystal structure for *Synechocystis* sp. PCC 6833 P-protein holoenzyme (PDB ID = 4lhc; sequence identity with human = 56.8%). Human H-protein was modeled using as the template bovine H-protein crystal structure (PDB ID = 3wdn, sequence identity with human = 98.0%).

The active site, active site tunnel, and dimerization interface functional regions of human P-protein were inferred from sequence comparison to the *Synechocystis* holoenzyme crystal structure.

### Protein Imaging

All 3D protein images were generated using the free protein-modeling software Jmol.

### NKH Mutation Analysis

A comprehensive list of NKH-causing missense mutations was compiled through a literature search of previously published mutations and mining of missense mutations catalogued in the ClinVar database hosted by the National Center for Biotechnology Information (NCBI). Missense mutations in ClinVar reported as “Benign” or “Likely Benign” were excluded. Secondary structure location of missense mutations was determined using jMol. Allele frequencies, if reported, were extracted from the Exome Aggregation Consortium (ExAC) database (Broad Institute)[33].

### Ligand Prediction

Structure-based ligand predictions were done using the I-Tasser COFACTOR tool provided by the Zhang Lab[34, 35] (University of Michigan).

### Protein-protein interaction modeling

The human P- and H-protein docking models were generated using the ClusPro 2.0 server[36–38] (Boston University). Models were generated as previously described. Briefly, the interacting proteins were docked using the fast-fourier transform (FFT) method. Highly populated clusters were selected and screened by CHARMM minimization.

### Model ranking

ClusPro 2.0 models were ranked using three equally weighted parameters: conservation of P-protein interacting residues, conservation of H-protein interacting residues, and distance of the active site lipoylated lysine from H-protein from the active site entry tunnel of P-protein. Scores were normalized such that the highest scoring model in a particular parameter received a score of 1, and the lowest scoring model received a score of 0. Interacting residues were defined as residues within 4 Angstroms from the docking partner. Conservation scores were generated using the Consurf server[39–41], which assigns a conservation score based on a multiple sequence alignment (MSA) with 150 unique orthologs. Parameters were tested using a previously-published crystal structure of the interaction between *E.* coli T- and H-protein (PDB: 3A8K)[23]. The interaction was modeled using ClusPro 2.0, and the models were scored using the above parameters. Top models were compared to the crystal structure. The interaction between human T- and H-protein and P- and T-protein (as a negative control) were also modeled using the above methods. Human T-protein has a previously published crystal structure (PDB: 1WSV)[42].

### Mutation effect on protein stability

Multiple online tools for prediction of ΔΔG caused by point mutations were preliminarily assessed, including CUPSAT, MuPro, PopMusic, STRUM, mCSM, FoldX, Dynamut, I-Mutant 3.0, SDM, and DeepDDG. CUPSAT uses protein structure to make fast and accurate ΔΔG predictions from environment-specific atomic and torsion angle potentials. The high speed of CUPSAT predictions lends itself to analyses of diseases associated with a large number of missense mutations[43]. CUPSAT was also recently found to be the most accurate predictor for disease-causing mutations in *PIXT2* that were known to be destabilizing[44]. Because of these considerations and because predictions are based on structural, site-specific information about the mutated residue, predictions of the stability effects of NKH-causing mutations were generated using CUPSAT. The SWISS-Model generated *GLDC* homology model was provided as the input. We defined destabilizing mutations as mutations with a predicted ΔΔG < −1.5 kcal/mol because a study found that 45 missense mutations associated with protein loss-of-function with experimentally derived ΔΔG’s had an average value of −1.67 kcal/mol[45].

### Multi-Parametric Mutation Score (MMS)

An eighteen-parameter test based on stability effects, conservation, location, and amino acid properties was used to score each mutation. Parameters were defined as the following: **i**. Destabilizing ΔΔG as predicted by CUPSAT. **ii.** Highly stabilizing mutation (positive ΔΔG, because rigidity in the protein is known to cause a loss of function). **iii.** Evolutionary conservation of the mutated amino acid (based on 150 homologs from different species because high conservation indicates that the residue is vital to the function or fold of the protein) **iv.** Conservation of amino acid substitution (based on Blosum62 matrix which indicates how likely it is that one amino acid will be substituted for another), **v.** Location in secondary structure domains of α-helix or **vi.** β-sheet because secondary structures contribute to fold and function of proteins **vii**. location in C-terminus because the catalytic activity is primarily in the C terminus **viii.** Residue is part of the active site (directly involved in binding/stabilizing PLP or substrate glycine, because it is essential to the catalytic function of the protein). **ix**. Mutation is within 7 Angstroms of the active site (because mutations in 3D proximity to the active site can affect its fold) **x**. Residue is part of the N-term PLP pocket (this study, because this will affect stability and/or catalytic activity.) **xi**. Residue is at P-H interaction interface (because this too can affect stability and/or catalytic activity). **xii**. Residue is at the known dimerization interface (because dimerization may be important to function). Xii-xviii. The mutation results in a change of amino acid properties, including change in **xiii**. polarity, **xiv**. charge, **xv**. aromaticity, **xvi.** to/from proline, **xvii**. tRNA availability, and **xviii**. volume.

Phi correlation analysis demonstrated independence between all parameters, except for slight negative correlations between conservation of amino acid substitution and change in polarity (φ = −0.44, with 1 being perfect correlation) and change in volume (φ = −0.51) (Fig 5B). However, these correlations are statistically weak and therefore do not negate the validity of including each of these parameters.

These eighteen parameters were combined to create a multiparametric mutation score (MMS), which scores mutations based the broad categories 1) stability effects, 2) conservation of mutation amino acid, 3) position of the mutated amino acid, and 4) change in amino acid properties caused by substitution. Mutations with scores of 1-2 were considered mild, 3-4, moderate, and ≥5 severe.

### NKH Patient Clinical Outcome Scoring

Phenotypic data were collected from case studies for 131 patients in the literature where the genotype was known and the patient had at least one missense mutation [13,14,29,46–56]. A comprehensive list of NKH symptoms in the clinical data was created (Table 2), and we developed a clinical outcome scoring scale based on the four major symptomatic domains of 1) seizures, 2) cognitive disorders, 3) muscle/movement control and 4) brain malformations (Table 3). The cognitive disorders and muscle/movement control domains were assigned linear Likert-like scores from 0-3 which represent the observed severity gradation of these domains in NKH patients. The seizure domain has a non-linear increase from 1 to 3 for controlled seizure activity and uncontrolled seizure activity, respectively. This increase more accurately captures the severity of the intractable seizure phenotype. The brain malformation domain is a binary choice of 0 or 3, because unlike the other symptom domains which have gradations of severity, brain malformations are a presence/absence binary. We concluded that the seriousness of a structural brain malformations warranted a score of 3. Summation of all four domains yields a patient clinical outcome score (COS), with a maximal score of 12.

Patients who were deceased at the time of the case report were not scored, leaving 86 patients. Domains that were reported as asymptomatic in the case report were scored 0. Domains that were not mentioned were left blank. Patients for whom only one major domain was able to be scored (12 of the 86 patients) were excluded from further analyses, leaving a 74-patient cohort.

### Homozygous Mutation Clinical Outcome Scores

Twenty-four of the 74 patient cohort were homozygous for 18 NKH-causing missense mutations. In cases where the mutation was present in one individual, the mutation was assigned the same COS as the individual. In cases where more than one individual was homozygous for the same mutation, an average COS was taken (Table 4).

### Weighted Multiparametric Mutation Score (WMMS)

#### i. Homozygous Mutations

The MMS was first applied to all 18 homozygous mutations associated with 24 clinical cases. Every patient mutation was assigned an MMS score in addition to the previously determined COS. Ten variants of GLDC were found in homozygous form in the Exome Aggregation Consortium (ExAC) database, hosted by the Broad Institute were also scored. The ExAC database utilizes genomic data from healthy individuals; thus, homozygosity of these mutations indicates that they are non-pathogenic. They were included therefore as a non-pathogenic control group and were assigned a COS of zero.

The COS was plotted as a function of the MMS of homozygous mutations, and this correlation was used to optimize parameter weights in order to yield a model with more biological and clinical value. Weighting was automated using Python. For all but one parameter, we began with the selection of a random parameter that was given weights from 0 to 5.0 in 0.1 increments thereby enabling up to a fifty-fold difference in weight and sufficient range for each parameter. The exception, which was conserved amino acid substitution, was allowed to range from −5.0 to 0. This was to confer a negative value or zero for a conserved substitution (for example, an R→K substitution) and prevent inappropriate elevation of score. We did not want to further increase the weight limit or reduce the increment size because this greatly increases computation time.

#### ii. Compound Heterozygous Mutations

For compound heterozygous mutants, deletions, nonsense mutations, intronic mutations, and mutations in mitochondrial signal sequence were each assigned scores of 5.0, corresponding to a moderate MMS score. We then calculated a composite score for each compound heterozygous patient according to the formula ½ Allele 1 Score + ½ Allele 2 Score, where the allele score for missense mutations was the MMS, and the score for other mutations was fixed at 5.0. Variants from healthy individuals (obtained from dbGaP), were included as non-pathogenic controls with a COS of zero. MMS parameters were optimized as above using the best fit line of the COS vs the composite score. For deletion, nonsense, intronic, and mitochondrial signal sequence mutation scores, since severe missense mutations WMMS range from 12-15, we selected a range of 0-20 for the remaining categories of mutations. Therefore, each mutation category was allowed to range from 0 to 20 in 0.1 increments.

dbGaP Data was obtained from the following datasets: ATVB - MIGen Exome Sequencing: Italian Atherosclerosis Thrombosis and Vascular Biology, phs000814.v1.p1; PROMIS - MIGen Exome Sequencing: Pakistan Risk Of Myocardial Infarction Study, phs000917.v1.p1; IBD - Inflammatory Bowel Disease Exome Sequencing Study, phs001076.v1.p1; Ottawa - MIGen Exome Sequencing: Ottawa Heart, phs000806.v1.p1.

#### iii. For both homozygous and heterozygous mutations

Ideally, the homozygous-trained parameters and heterozygous-trained parameters would have approximately equal weights. The weight of the stabilizing mutations and the change in polarity parameters regressed to 0 for each dataset, indicating that these parameters have a negligible effect on disease severity. Active site mutations in each case were the most highly weighted parameter with weights of 4.2 for homozygous and 5.0 for heterozygous mutations. This indicates, as would be expected, that active site mutations severely affect protein function. For both the homozygous- and heterozygous-trained sets, the parameters helix mutations, Δ aromaticity, and Δ size are within weights of +/- 1. For the homozygous mutations, none of the mutations were in the active region or dimerization interface, and thus these parameters could not be optimized and were set to 0. The other 10 parameters have weight differences the larger than +/- 1, indicating either that the parameter weighting for these parameters is biased by the data that it’s trained on, or that these parameters have different degrees of importance for heterozygous and homozygous mutations.

## Acknowledgements

We thank Stefan Freed and Dr. Tobin Sosnic for their helpful suggestions and comments on this manuscript. We thank the Broad Institute for generating high-quality sequence data supported by NHGRI funds (grant # U54 HG003067) with Eric Lander as PI. dbGaP data was obtained at http://www.ncbi.nlm.nih.gov/gap through accession numbers phs000814.v1.p1, phs000917.v1.p1, phs001076.v1.p1, and phs000806.v1.p1.

## Supporting Information

**S1 Figure. Human GLDC N- and C-terminal alignment**

**S2 Figure. T- and H-protein interaction model**

**S3 Figure. Clinical outcome score vs multiparametric mutation score**

**S1 Table. *E. Coli* T- and H-protein interaction model scores.**

**S2 Table. Human GLDC and H-protein interaction model scores**

**S3 Table. CUPSAT ΔΔG predictions**

**S4 Table. NKH patient clinical outcome scores**

**S5 Table. NKH multiparametric mutation scores**

**S6 Table. Homozygous mutation weighted multiparametric mutation scores**

**S7 Table. Heterozygous mutation weighted multiparametric mutation scores**

**S1 Appendix. ClusPro 2.0 interaction model PDB files**

**S1 Figure.**
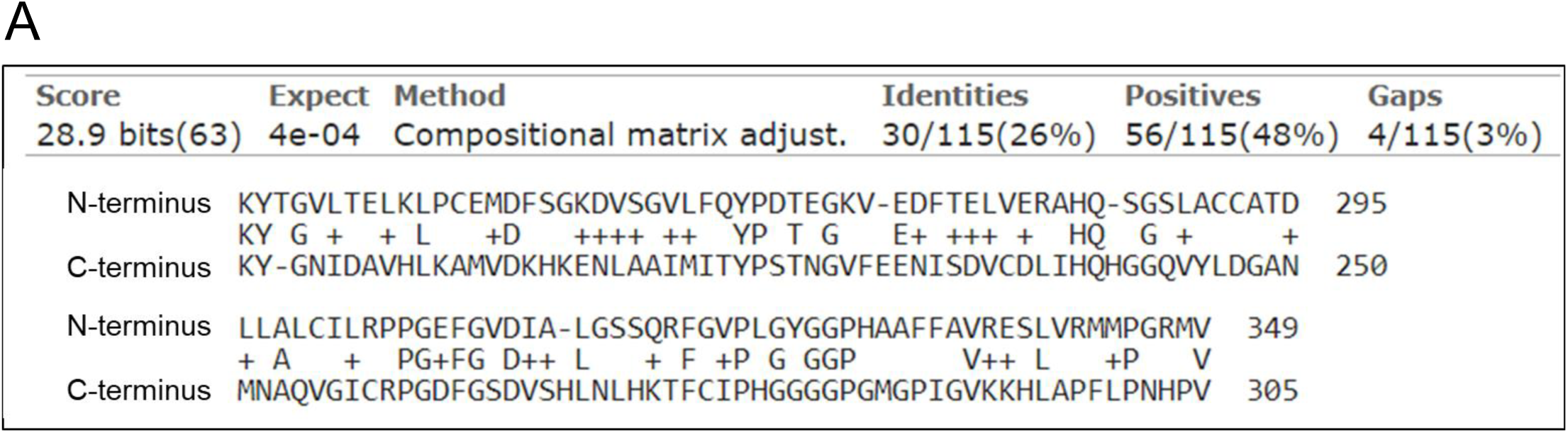
pBlast Alignment between the N- and C-termini PLP subunits of human GLDC. Alignment reveals a region of similarity with 26% identity and 48% similarity.

**S2 Figure.**
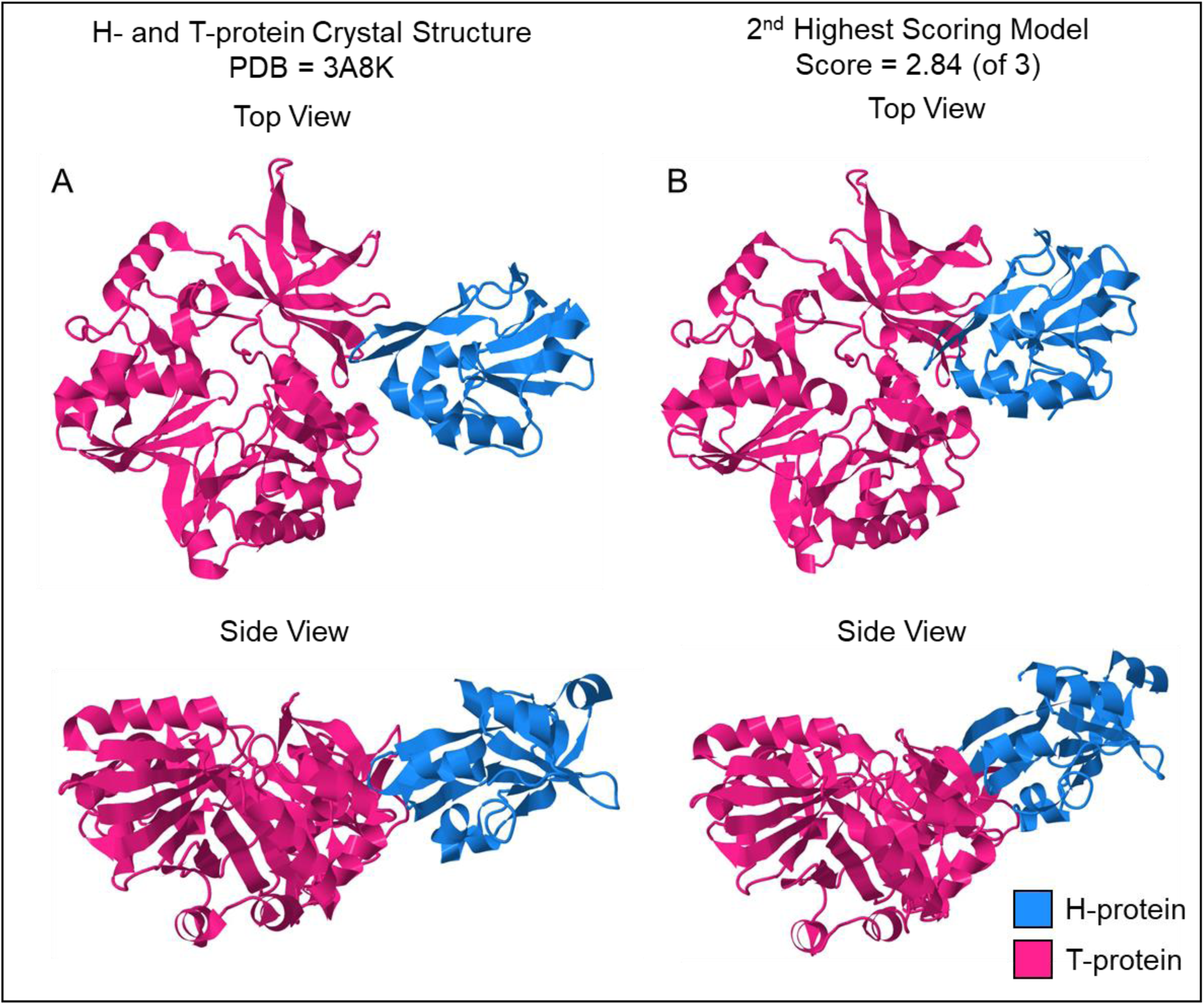
Verification of novel interaction model parameters. [A] Crystal structure of the interaction between T- and H-protein (PDB: 3A8K). [B] The second highest scoring modeled interaction between T- and H-protein with a score of 2.64. The highest-ranking model (score = 2.67) had H-protein flipped but docking at the same site on T-protein.

**S3 Figure.**
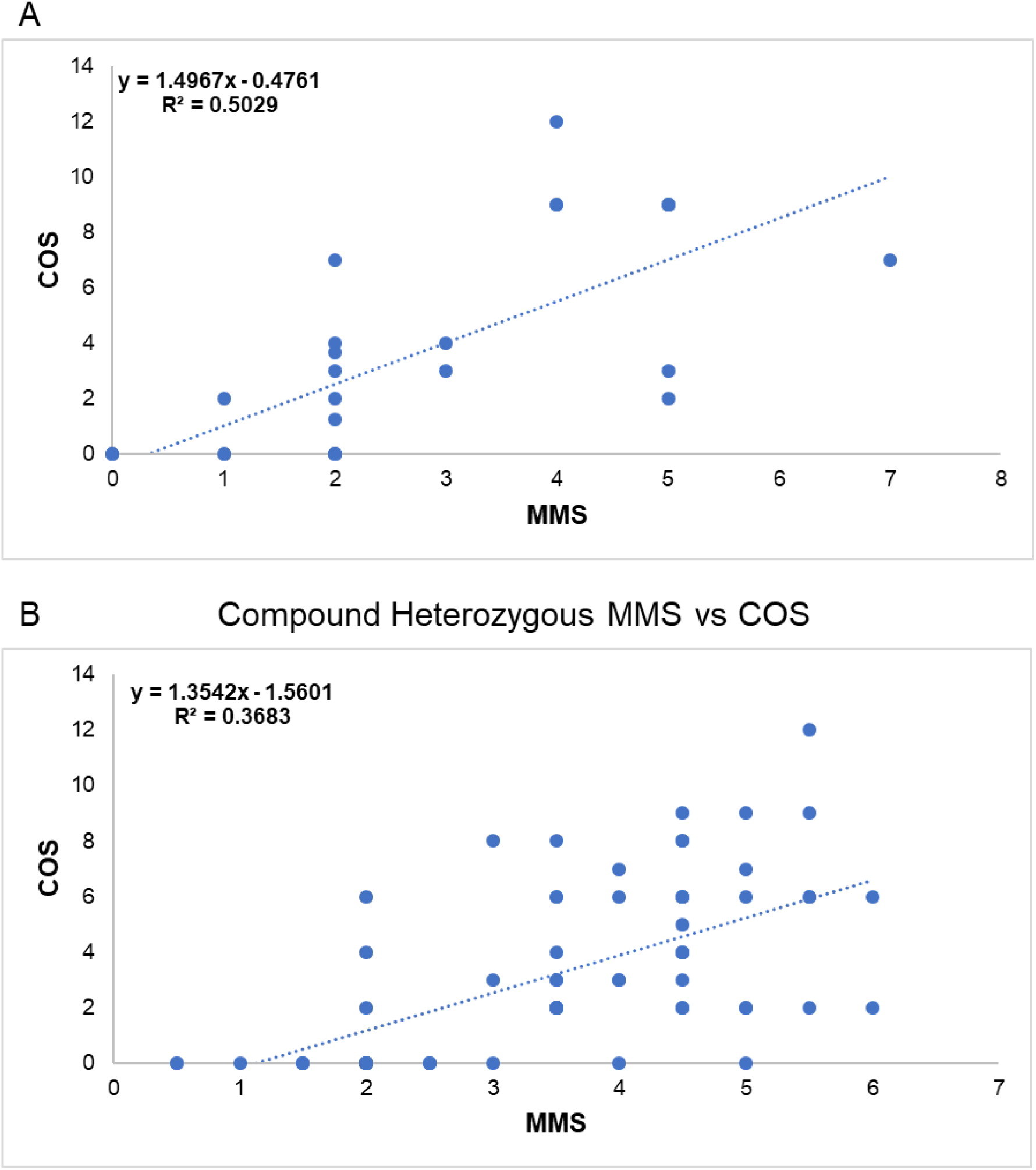
[A] MMS vs COS for homozygous patients and nonpathogenic controls. There is a positive correlation (R2 = 0.50) between the MMS and COS. [B] MMS vs COS for compound heterozygous patients and nonpathogenic controls. There is a slight positive correlation (R2 = 0.37) between the MMS and COS.

