## Supplemental Table 1 for "Large scale analyses of genotype-phenotype relationships of glycine decarboxylase mutations and neurological disease severity"

| T + H-protein E. Coli Scoring | | | | | | | |
|---|---|---|---|---|---|---|---|
| Parameter | **T-protein Conservation** | | **H-protein Conservation** | | **H lipoyllysine distance** | | **TOTAL** |
| *Model ##* | *Raw Score* | *Normalized* | *Raw Score* | *Normalized* | *Raw Score* | *Normalized* | *Score* |
| model.004.15 | -0.585 | 0.966 | -0.510 | 0.885 | 9.123 | 0.819 | 2.671 |
| model.004.04 | -0.628 | 1.000 | -0.464 | 0.850 | 9.433 | 0.792 | 2.642 |
| model.000.11 | -0.596 | 0.975 | -0.289 | 0.718 | 8.330 | 0.897 | 2.590 |
| model.004.13 | -0.517 | 0.912 | -0.267 | 0.702 | 8.353 | 0.895 | 2.509 |
| model.000.14 | -0.297 | 0.736 | -0.336 | 0.754 | 7.473 | 1.000 | 2.490 |
| model.004.06 | -0.490 | 0.890 | -0.238 | 0.680 | 9.478 | 0.788 | 2.358 |
| model.000.08 | -0.413 | 0.829 | -0.197 | 0.649 | 8.818 | 0.847 | 2.326 |
| model.004.08 | -0.212 | 0.669 | -0.254 | 0.692 | 8.400 | 0.890 | 2.250 |
| model.002.01 | -0.530 | 0.922 | -0.515 | 0.889 | 18.740 | 0.399 | 2.210 |
| model.004.10 | -0.382 | 0.804 | -0.439 | 0.832 | 13.465 | 0.555 | 2.191 |
| model.004.14 | -0.462 | 0.868 | -0.442 | 0.834 | 15.523 | 0.481 | 2.183 |
| model.000.04 | -0.530 | 0.922 | -0.515 | 0.889 | 22.497 | 0.332 | 2.143 |
| model.004.00 | -0.429 | 0.841 | -0.413 | 0.812 | 18.645 | 0.401 | 2.054 |
| model.004.05 | -0.422 | 0.836 | -0.297 | 0.725 | 16.568 | 0.451 | 2.012 |
| model.004.02 | -0.308 | 0.745 | -0.454 | 0.843 | 18.795 | 0.398 | 1.986 |
| model.004.01 | -0.484 | 0.886 | -0.330 | 0.749 | 21.313 | 0.351 | 1.985 |
| model.002.07 | -0.259 | 0.706 | -0.399 | 0.801 | 16.273 | 0.459 | 1.967 |
| model.000.01 | -0.224 | 0.679 | -0.409 | 0.809 | 16.475 | 0.454 | 1.941 |
| model.002.00 | -0.035 | 0.528 | -0.662 | 1.000 | 18.178 | 0.411 | 1.939 |
| model.000.15 | -0.469 | 0.874 | 0.001 | 0.499 | 13.595 | 0.550 | 1.923 |
| model.004.03 | -0.468 | 0.873 | 0.000 | 0.500 | 13.893 | 0.538 | 1.911 |
| model.000.00 | -0.033 | 0.526 | -0.566 | 0.927 | 16.920 | 0.442 | 1.895 |
| model.004.18 | -0.431 | 0.843 | -0.179 | 0.635 | 19.103 | 0.391 | 1.869 |
| model.004.16 | -0.050 | 0.540 | -0.526 | 0.898 | 18.185 | 0.411 | 1.848 |
| model.004.07 | -0.092 | 0.573 | -0.415 | 0.814 | 16.350 | 0.457 | 1.844 |
| model.002.13 | -0.318 | 0.753 | -0.257 | 0.694 | 19.470 | 0.384 | 1.831 |
| model.000.13 | -0.596 | 0.975 | 0.008 | 0.494 | 20.823 | 0.359 | 1.828 |
| model.006.18 | -0.028 | 0.522 | -0.571 | 0.932 | 20.823 | 0.359 | 1.813 |
| model.000.03 | -0.361 | 0.787 | -0.220 | 0.666 | 21.433 | 0.349 | 1.802 |
| model.002.09 | -0.110 | 0.588 | -0.429 | 0.824 | 19.495 | 0.383 | 1.795 |
| model.002.15 | -0.084 | 0.567 | -0.355 | 0.768 | 17.475 | 0.428 | 1.763 |
| model.002.02 | -0.048 | 0.538 | -0.398 | 0.800 | 18.120 | 0.412 | 1.751 |
| model.004.17 | -0.164 | 0.630 | -0.003 | 0.503 | 12.470 | 0.599 | 1.732 |
| model.000.07 | -0.478 | 0.881 | -0.045 | 0.534 | 26.748 | 0.279 | 1.694 |
| model.002.18 | -0.075 | 0.560 | -0.415 | 0.814 | 23.443 | 0.319 | 1.692 |
| model.002.06 | 0.071 | 0.424 | -0.479 | 0.862 | 19.313 | 0.387 | 1.672 |
| model.000.06 | -0.365 | 0.791 | -0.042 | 0.531 | 22.285 | 0.335 | 1.657 |
| model.006.21 | 0.135 | 0.354 | -0.532 | 0.902 | 21.120 | 0.354 | 1.609 |
| model.006.15 | 0.199 | 0.285 | -0.604 | 0.957 | 25.233 | 0.296 | 1.538 |
| model.000.10 | -0.517 | 0.912 | 0.250 | 0.336 | 25.923 | 0.288 | 1.536 |
| model.000.20 | 0.140 | 0.349 | -0.335 | 0.753 | 19.043 | 0.392 | 1.494 |
| model.004.12 | -0.324 | 0.758 | 0.008 | 0.495 | 31.145 | 0.240 | 1.493 |
| model.004.09 | -0.232 | 0.685 | -0.078 | 0.559 | 30.685 | 0.244 | 1.487 |
| model.000.18 | -0.223 | 0.678 | -0.134 | 0.602 | 38.658 | 0.193 | 1.473 |

| | | | | | | | |
|---|---|---|---|---|---|---|---|
| model.000.22 | -0.304 | 0.742 | 0.122 | 0.420 | 26.800 | 0.279 | 1.441 |
| model.006.06 | 0.182 | 0.303 | -0.328 | 0.747 | 21.728 | 0.344 | 1.394 |
| model.000.21 | 0.042 | 0.455 | -0.243 | 0.684 | 30.475 | 0.245 | 1.384 |
| model.000.19 | -0.293 | 0.733 | 0.130 | 0.415 | 32.395 | 0.231 | 1.379 |
| model.000.12 | -0.418 | 0.833 | 0.318 | 0.292 | 30.425 | 0.246 | 1.370 |
| model.006.22 | 0.024 | 0.474 | -0.087 | 0.566 | 23.038 | 0.324 | 1.364 |
| model.000.16 | -0.285 | 0.727 | 0.106 | 0.430 | 36.595 | 0.204 | 1.362 |
| model.006.01 | 0.151 | 0.336 | -0.274 | 0.707 | 26.688 | 0.280 | 1.323 |
| model.006.20 | 0.203 | 0.280 | -0.326 | 0.747 | 29.333 | 0.255 | 1.282 |
| model.000.05 | -0.016 | 0.513 | 0.007 | 0.495 | 29.473 | 0.254 | 1.262 |
| model.002.03 | -0.016 | 0.513 | 0.007 | 0.495 | 29.473 | 0.254 | 1.262 |
| model.002.17 | -0.079 | 0.563 | 0.038 | 0.475 | 34.745 | 0.215 | 1.253 |
| model.002.04 | 0.019 | 0.479 | 0.022 | 0.486 | 29.648 | 0.252 | 1.217 |
| model.006.09 | -0.022 | 0.518 | 0.063 | 0.459 | 32.175 | 0.232 | 1.209 |
| model.006.13 | 0.135 | 0.354 | -0.090 | 0.568 | 26.705 | 0.280 | 1.202 |
| model.002.12 | 0.034 | 0.464 | -0.033 | 0.525 | 35.300 | 0.212 | 1.201 |
| model.002.16 | 0.048 | 0.448 | -0.058 | 0.543 | 36.548 | 0.204 | 1.196 |
| model.002.08 | -0.067 | 0.553 | 0.110 | 0.428 | 35.998 | 0.208 | 1.189 |
| model.002.14 | 0.083 | 0.410 | -0.012 | 0.509 | 28.560 | 0.262 | 1.181 |
| model.000.17 | 0.211 | 0.272 | -0.246 | 0.686 | 34.045 | 0.219 | 1.177 |
| model.006.05 | 0.155 | 0.333 | -0.111 | 0.584 | 30.238 | 0.247 | 1.164 |
| model.002.05 | 0.175 | 0.310 | -0.083 | 0.563 | 26.773 | 0.279 | 1.152 |
| model.000.09 | 0.153 | 0.334 | -0.032 | 0.524 | 29.073 | 0.257 | 1.116 |
| model.006.00 | 0.130 | 0.359 | 0.099 | 0.435 | 30.733 | 0.243 | 1.038 |
| model.006.23 | 0.163 | 0.324 | 0.066 | 0.457 | 31.353 | 0.238 | 1.019 |
| model.002.10 | 0.058 | 0.437 | 0.233 | 0.348 | 32.408 | 0.231 | 1.015 |
| model.000.02 | -0.084 | 0.567 | 0.472 | 0.191 | 34.095 | 0.219 | 0.977 |
| model.004.11 | 0.462 | 0.000 | -0.259 | 0.696 | 31.763 | 0.235 | 0.931 |
| model.006.08 | 0.134 | 0.355 | 0.263 | 0.328 | 34.053 | 0.219 | 0.902 |
| model.006.17 | 0.191 | 0.293 | 0.172 | 0.387 | 35.268 | 0.212 | 0.892 |
| model.006.19 | 0.207 | 0.276 | 0.140 | 0.408 | 36.760 | 0.203 | 0.887 |
| model.006.12 | 0.156 | 0.331 | 0.263 | 0.327 | 37.295 | 0.200 | 0.859 |
| model.006.07 | 0.192 | 0.292 | 0.230 | 0.349 | 40.775 | 0.183 | 0.825 |
| model.006.16 | 0.220 | 0.262 | 0.251 | 0.335 | 33.055 | 0.226 | 0.823 |
| model.006.03 | 0.264 | 0.215 | 0.177 | 0.384 | 42.125 | 0.177 | 0.776 |
| model.006.14 | 0.401 | 0.066 | 0.048 | 0.469 | 31.085 | 0.240 | 0.775 |
| model.006.02 | 0.328 | 0.145 | 0.095 | 0.438 | 43.578 | 0.171 | 0.755 |
| model.002.11 | 0.008 | 0.491 | 0.666 | 0.063 | 49.423 | 0.151 | 0.706 |
| model.006.10 | 0.002 | 0.498 | 0.763 | 0.000 | 51.260 | 0.146 | 0.644 |
| model.006.11 | 0.227 | 0.255 | 0.740 | 0.015 | 44.355 | 0.168 | 0.438 |
| model.006.04 | 0.449 | 0.014 | 0.450 | 0.205 | 38.315 | 0.195 | 0.414 |
