## Supplemental Table 2 for "Large scale analyses of genotype-phenotype relationships of glycine decarboxylase mutations and neurological disease severity"

| GLDC + H-protein Round 1 Scoring | | | | | | | |
|---|---|---|---|---|---|---|---|
| Parameter | **GLDC Conservation** | | **H-protein Conservation** | | **H lipoyllysine distance** | | **TOTAL** |
| *Model ##* | *Raw Score* | *Normalized* | *Raw Score* | *Normalized* | *Raw Score* | *Normalized* | *Score* |
| model.000.24 | -0.568 | 0.942 | -0.589 | 0.827 | 6.775 | 0.992 | 2.761 |
| model.002.03 | -0.536 | 0.918 | -0.631 | 0.851 | 8.553 | 0.965 | 2.734 |
| model.002.23 | -0.576 | 0.948 | -0.585 | 0.825 | 8.988 | 0.959 | 2.732 |
| model.002.06 | -0.548 | 0.927 | -0.424 | 0.735 | 12.130 | 0.912 | 2.574 |
| model.002.07 | -0.563 | 0.938 | -0.407 | 0.726 | 13.123 | 0.897 | 2.561 |
| model.002.26 | -0.610 | 0.975 | -0.514 | 0.785 | 22.658 | 0.754 | 2.515 |
| model.000.25 | -0.381 | 0.797 | -0.399 | 0.721 | 8.820 | 0.961 | 2.480 |
| model.000.19 | -0.425 | 0.831 | -0.289 | 0.660 | 7.743 | 0.977 | 2.469 |
| model.006.07 | -0.300 | 0.734 | -0.665 | 0.869 | 16.183 | 0.851 | 2.454 |
| model.002.02 | -0.326 | 0.754 | -0.775 | 0.930 | 21.718 | 0.769 | 2.453 |
| model.002.13 | -0.581 | 0.953 | -0.150 | 0.583 | 13.188 | 0.896 | 2.432 |
| model.002.12 | -0.184 | 0.643 | -0.641 | 0.856 | 16.030 | 0.854 | 2.353 |
| model.006.10 | -0.599 | 0.966 | -0.235 | 0.630 | 26.560 | 0.696 | 2.293 |
| model.000.29 | -0.566 | 0.941 | -0.235 | 0.630 | 27.485 | 0.682 | 2.254 |
| model.002.17 | -0.501 | 0.890 | -0.197 | 0.610 | 22.868 | 0.751 | 2.251 |
| model.002.01 | -0.374 | 0.791 | -0.392 | 0.718 | 23.783 | 0.738 | 2.247 |
| model.002.04 | -0.053 | 0.542 | -0.671 | 0.873 | 17.888 | 0.826 | 2.240 |
| model.002.05 | -0.229 | 0.679 | -0.385 | 0.714 | 16.583 | 0.845 | 2.237 |
| model.006.14 | -0.184 | 0.644 | -0.637 | 0.854 | 23.705 | 0.739 | 2.236 |
| model.000.16 | -0.435 | 0.839 | -0.281 | 0.656 | 24.258 | 0.731 | 2.226 |
| model.002.22 | -0.642 | 1.000 | -0.077 | 0.543 | 27.638 | 0.680 | 2.223 |
| model.000.26 | -0.225 | 0.675 | -0.469 | 0.760 | 21.145 | 0.777 | 2.212 |
| model.002.19 | -0.064 | 0.550 | -0.605 | 0.836 | 20.143 | 0.792 | 2.178 |
| model.002.14 | -0.359 | 0.779 | -0.245 | 0.636 | 22.380 | 0.759 | 2.174 |
| model.006.01 | -0.313 | 0.744 | -0.317 | 0.676 | 22.965 | 0.750 | 2.170 |
| model.006.24 | -0.262 | 0.704 | -0.438 | 0.743 | 25.333 | 0.714 | 2.162 |
| model.006.23 | -0.484 | 0.877 | -0.072 | 0.540 | 23.560 | 0.741 | 2.158 |
| model.002.00 | 0.140 | 0.456 | -0.738 | 0.910 | 20.220 | 0.791 | 2.157 |
| model.002.11 | -0.603 | 0.969 | 0.026 | 0.479 | 26.093 | 0.703 | 2.152 |
| model.002.24 | -0.242 | 0.689 | -0.533 | 0.796 | 29.638 | 0.650 | 2.135 |
| model.006.26 | -0.156 | 0.621 | -0.626 | 0.848 | 28.680 | 0.664 | 2.133 |
| model.000.00 | -0.115 | 0.590 | -0.403 | 0.724 | 20.703 | 0.784 | 2.097 |
| model.006.13 | -0.291 | 0.727 | -0.188 | 0.604 | 21.950 | 0.765 | 2.096 |
| model.006.15 | -0.312 | 0.743 | 0.024 | 0.480 | 15.005 | 0.869 | 2.092 |
| model.000.10 | -0.437 | 0.840 | 0.043 | 0.465 | 22.075 | 0.763 | 2.069 |
| model.006.18 | -0.378 | 0.795 | -0.281 | 0.656 | 32.890 | 0.601 | 2.052 |
| model.006.19 | -0.220 | 0.672 | -0.479 | 0.766 | 33.123 | 0.598 | 2.035 |
| model.000.11 | -0.302 | 0.735 | -0.012 | 0.507 | 20.913 | 0.781 | 2.023 |
| model.002.25 | -0.305 | 0.737 | -0.290 | 0.661 | 32.510 | 0.607 | 2.005 |
| model.006.11 | -0.428 | 0.834 | 0.029 | 0.476 | 27.538 | 0.681 | 1.992 |
| model.006.22 | -0.362 | 0.782 | -0.059 | 0.533 | 29.555 | 0.651 | 1.966 |
| model.002.08 | -0.461 | 0.859 | -0.018 | 0.510 | 33.503 | 0.592 | 1.962 |
| model.006.05 | -0.281 | 0.719 | -0.039 | 0.522 | 25.238 | 0.716 | 1.956 |
| model.002.20 | 0.159 | 0.450 | -0.684 | 0.880 | 31.540 | 0.622 | 1.952 |

| | | | | | | | |
|---|---|---|---|---|---|---|---|
| model.000.20 | -0.117 | 0.591 | -0.750 | 0.916 | 44.683 | 0.425 | 1.932 |
| model.006.12 | -0.419 | 0.826 | -0.050 | 0.528 | 35.033 | 0.569 | 1.924 |
| model.006.27 | -0.391 | 0.805 | 0.111 | 0.411 | 26.135 | 0.702 | 1.918 |
| model.002.09 | -0.183 | 0.643 | -0.269 | 0.649 | 32.085 | 0.613 | 1.906 |
| model.000.18 | -0.299 | 0.733 | -0.246 | 0.637 | 37.358 | 0.535 | 1.904 |
| model.002.16 | 0.042 | 0.487 | -0.421 | 0.734 | 28.578 | 0.666 | 1.886 |
| model.006.20 | -0.301 | 0.734 | 0.197 | 0.341 | 19.985 | 0.794 | 1.870 |
| model.000.27 | 0.215 | 0.433 | -0.551 | 0.806 | 31.400 | 0.624 | 1.862 |
| model.000.17 | 0.124 | 0.461 | -0.634 | 0.852 | 36.793 | 0.543 | 1.856 |
| model.006.04 | 0.024 | 0.492 | -0.157 | 0.587 | 21.615 | 0.770 | 1.849 |
| model.000.23 | -0.213 | 0.666 | -0.044 | 0.525 | 29.843 | 0.647 | 1.837 |
| model.006.06 | -0.257 | 0.700 | 0.363 | 0.208 | 11.970 | 0.914 | 1.823 |
| model.006.09 | 0.046 | 0.486 | -0.270 | 0.650 | 27.253 | 0.686 | 1.821 |
| model.006.02 | 0.261 | 0.418 | -0.205 | 0.614 | 22.143 | 0.762 | 1.794 |
| model.004.10 | 0.282 | 0.412 | -0.666 | 0.870 | 39.365 | 0.505 | 1.786 |
| model.000.05 | -0.035 | 0.528 | -0.785 | 0.936 | 51.598 | 0.322 | 1.785 |
| model.000.15 | 0.139 | 0.457 | -0.181 | 0.601 | 25.633 | 0.710 | 1.767 |
| model.006.03 | -0.022 | 0.517 | 0.018 | 0.485 | 22.373 | 0.759 | 1.761 |
| model.000.09 | 0.049 | 0.485 | -0.564 | 0.813 | 42.300 | 0.461 | 1.759 |
| model.006.25 | -0.432 | 0.836 | 0.383 | 0.192 | 26.125 | 0.703 | 1.731 |
| model.002.18 | -0.181 | 0.641 | 0.136 | 0.391 | 27.915 | 0.676 | 1.707 |
| model.000.01 | 0.325 | 0.398 | -0.517 | 0.787 | 40.775 | 0.484 | 1.669 |
| model.006.16 | 0.015 | 0.495 | -0.159 | 0.588 | 36.350 | 0.550 | 1.633 |
| model.006.21 | -0.291 | 0.727 | 0.305 | 0.255 | 33.163 | 0.597 | 1.579 |
| model.006.00 | 0.003 | 0.499 | -0.224 | 0.624 | 42.765 | 0.454 | 1.577 |
| model.000.28 | -0.106 | 0.582 | -0.084 | 0.547 | 47.518 | 0.383 | 1.512 |
| model.006.17 | 0.001 | 0.500 | -0.115 | 0.564 | 48.488 | 0.368 | 1.432 |
| model.004.04 | 1.164 | 0.136 | -0.708 | 0.893 | 46.995 | 0.391 | 1.419 |
| model.000.03 | 1.164 | 0.136 | -0.685 | 0.880 | 46.555 | 0.397 | 1.413 |
| model.006.08 | -0.143 | 0.611 | 0.206 | 0.335 | 41.995 | 0.465 | 1.411 |
| model.002.21 | 1.252 | 0.108 | -0.671 | 0.873 | 45.758 | 0.409 | 1.390 |
| model.000.08 | 0.416 | 0.370 | -0.241 | 0.634 | 47.403 | 0.384 | 1.388 |
| model.002.15 | 0.038 | 0.488 | -0.042 | 0.523 | 47.920 | 0.377 | 1.388 |
| model.004.00 | 1.481 | 0.037 | -0.609 | 0.838 | 48.313 | 0.371 | 1.246 |
| model.004.12 | 1.417 | 0.057 | -0.730 | 0.905 | 54.818 | 0.274 | 1.235 |
| model.004.13 | 1.503 | 0.030 | -0.566 | 0.814 | 48.108 | 0.374 | 1.218 |
| model.004.11 | 1.184 | 0.129 | -0.381 | 0.711 | 48.828 | 0.363 | 1.204 |
| model.000.12 | 1.532 | 0.020 | -0.604 | 0.835 | 50.100 | 0.344 | 1.200 |
| model.000.02 | 1.449 | 0.046 | -0.453 | 0.751 | 51.498 | 0.323 | 1.121 |
| model.004.02 | 1.377 | 0.069 | -0.421 | 0.734 | 51.875 | 0.318 | 1.120 |
| model.004.08 | 1.598 | 0.000 | -0.418 | 0.732 | 49.450 | 0.354 | 1.086 |
| model.000.06 | 1.230 | 0.115 | -0.295 | 0.664 | 52.890 | 0.302 | 1.081 |
| model.000.21 | 1.144 | 0.142 | -0.145 | 0.581 | 49.305 | 0.356 | 1.078 |
| model.004.07 | 1.242 | 0.111 | -0.281 | 0.656 | 52.513 | 0.308 | 1.075 |
| model.000.22 | 1.280 | 0.099 | -0.272 | 0.651 | 57.375 | 0.235 | 0.986 |
| model.004.01 | 1.503 | 0.030 | -0.301 | 0.667 | 55.160 | 0.268 | 0.965 |
| model.000.14 | 1.025 | 0.179 | -0.069 | 0.538 | 57.018 | 0.241 | 0.958 |

| | | | | | | | |
|---|---|---|---|---|---|---|---|
| model.004.03 | 1.503 | 0.030 | -0.555 | 0.808 | 72.420 | 0.010 | 0.848 |
| model.004.14 | 1.460 | 0.043 | -0.421 | 0.734 | 70.190 | 0.044 | 0.820 |
| model.000.07 | 1.578 | 0.006 | -0.458 | 0.754 | 72.888 | 0.003 | 0.764 |
| model.000.13 | 0.161 | 0.450 | 0.622 | 0.000 | 53.568 | 0.292 | 0.742 |
| model.004.09 | 1.511 | 0.027 | -0.321 | 0.678 | 73.115 | 0.000 | 0.706 |
| model.004.06 | 1.131 | 0.146 | 0.147 | 0.382 | 70.183 | 0.044 | 0.571 |
| model.002.10 | 1.057 | 0.169 | 0.140 | 0.387 | 72.345 | 0.012 | 0.568 |
| model.004.05 | 1.598 | 0.000 | 0.001 | 0.500 | 71.203 | 0.029 | 0.528 |
| model.000.04 | 1.084 | 0.161 | 0.231 | 0.314 | 71.623 | 0.022 | 0.497 |

| GLDC + H-protein Round 2 Scoring | | | | | | | |
|---|---|---|---|---|---|---|---|
| Parameter | **GLDC Conservation** | | **H-protein Conservation** | | **H lipoyllysine distance** | | **TOTAL** |
| *Model ##* | *Raw Score* | *Normalized* | *Raw Score* | *Normalized* | *Raw Score* | *Normalized* | *Score* |
| model.004.01 | -0.486 | 0.878 | -0.786 | 0.936 | 6.858 | 0.991 | 2.805 |
| model.000.01 | -0.440 | 0.843 | -0.746 | 0.914 | 6.235 | 1.000 | 2.757 |
| model.002.01 | -0.440 | 0.843 | -0.746 | 0.914 | 6.235 | 1.000 | 2.757 |
| model.000.07 | -0.462 | 0.860 | -0.708 | 0.893 | 6.278 | 0.999 | 2.753 |
| model.006.09 | -0.600 | 0.967 | -0.668 | 0.871 | 12.708 | 0.903 | 2.741 |
| model.004.07 | -0.555 | 0.932 | -0.886 | 0.992 | 20.700 | 0.784 | 2.708 |
| model.006.04 | -0.513 | 0.899 | -0.494 | 0.774 | 9.238 | 0.955 | 2.628 |
| model.002.10 | -0.370 | 0.788 | -0.614 | 0.841 | 6.318 | 0.999 | 2.628 |
| model.004.00 | -0.436 | 0.840 | -0.901 | 1.000 | 24.323 | 0.730 | 2.569 |
| model.004.06 | -0.304 | 0.737 | -0.646 | 0.859 | 9.685 | 0.948 | 2.544 |
| model.002.00 | -0.445 | 0.847 | -0.660 | 0.867 | 24.283 | 0.730 | 2.443 |
| model.002.06 | -0.451 | 0.851 | -0.305 | 0.669 | 13.650 | 0.889 | 2.409 |
| model.002.03 | -0.558 | 0.935 | -0.197 | 0.609 | 15.365 | 0.863 | 2.408 |
| model.000.00 | -0.547 | 0.926 | -0.458 | 0.754 | 24.563 | 0.726 | 2.406 |
| model.004.05 | -0.606 | 0.972 | -0.047 | 0.526 | 13.323 | 0.894 | 2.393 |
| model.000.02 | -0.268 | 0.708 | -0.418 | 0.732 | 9.985 | 0.944 | 2.384 |
| model.002.04 | -0.268 | 0.708 | -0.418 | 0.732 | 9.985 | 0.944 | 2.384 |
| model.000.05 | -0.514 | 0.900 | -0.191 | 0.606 | 14.425 | 0.878 | 2.384 |
| model.000.03 | -0.327 | 0.755 | -0.390 | 0.717 | 12.715 | 0.903 | 2.375 |
| model.006.13 | -0.402 | 0.813 | -0.495 | 0.775 | 25.455 | 0.713 | 2.301 |
| model.004.03 | -0.349 | 0.772 | -0.201 | 0.612 | 12.733 | 0.903 | 2.287 |
| model.000.04 | -0.219 | 0.671 | -0.673 | 0.874 | 25.893 | 0.706 | 2.251 |
| model.006.02 | -0.517 | 0.903 | -0.036 | 0.520 | 18.155 | 0.822 | 2.245 |
| model.006.03 | -0.420 | 0.827 | -0.343 | 0.690 | 24.888 | 0.721 | 2.239 |
| model.006.08 | -0.161 | 0.625 | -0.673 | 0.874 | 26.088 | 0.703 | 2.202 |
| model.002.08 | -0.368 | 0.786 | -0.310 | 0.672 | 23.495 | 0.742 | 2.200 |
| model.004.04 | -0.179 | 0.639 | -0.549 | 0.805 | 24.648 | 0.725 | 2.169 |
| model.006.05 | -0.272 | 0.712 | -0.294 | 0.663 | 22.218 | 0.761 | 2.136 |
| model.002.07 | -0.277 | 0.716 | -0.258 | 0.643 | 21.550 | 0.771 | 2.130 |
| model.002.05 | -0.107 | 0.583 | -0.487 | 0.770 | 21.863 | 0.766 | 2.120 |
| model.002.02 | -0.307 | 0.739 | -0.225 | 0.625 | 25.388 | 0.714 | 2.077 |
| model.004.02 | -0.225 | 0.675 | -0.190 | 0.606 | 24.245 | 0.731 | 2.011 |
| model.006.10 | -0.142 | 0.611 | -0.178 | 0.599 | 22.525 | 0.756 | 1.966 |
| model.000.06 | -0.267 | 0.708 | 0.000 | 0.500 | 24.175 | 0.732 | 1.939 |
| model.006.06 | -0.363 | 0.783 | 0.085 | 0.432 | 25.650 | 0.710 | 1.924 |
| model.006.00 | -0.341 | 0.766 | 0.278 | 0.277 | 17.080 | 0.838 | 1.880 |
| model.006.12 | -0.567 | 0.942 | 0.319 | 0.244 | 28.535 | 0.667 | 1.852 |
| model.002.09 | -0.050 | 0.539 | 0.109 | 0.413 | 23.438 | 0.743 | 1.695 |
| model.006.07 | -0.066 | 0.551 | 0.218 | 0.325 | 19.800 | 0.797 | 1.673 |
| model.006.01 | -0.250 | 0.695 | 0.460 | 0.130 | 21.443 | 0.773 | 1.597 |
| model.006.11 | -0.011 | 0.509 | 0.162 | 0.370 | 25.913 | 0.706 | 1.584 |
