## Supplemental Table 3 for "Large scale analyses of genotype-phenotype relationships of glycine decarboxylase mutations and neurological disease severity"

| Mutation | Mut Ref | ddG (CUPSAT) |
|---|---|---|
| D36H | ClinVar - Unpub | N/A |
| S37N | ClinVar - Unpub | N/A |
| G48W | ClinVar - Unpub | N/A |
| R59T | Kure (2006) | -2.72 |
| P70R | Coughlin (2017) | 5.06 |
| L79P | Coughlin (2017) | -3.23 |
| L82S | Swanson (2015) | -1.81 |
| L82W | Kure (2004) | -0.76 |
| D88N | ClinVar - Unpub | 0.20 |
| L90F | ClinVar - Unpub | -1.85 |
| P110H | ClinVar - Unpub | -2.39 |
| S132L | Swanson (2015) | 0.66 |
| S132W | Coughlin (2017) | 3.25 |
| G135A | Coughlin (2017) | -6.02 |
| Y138D | Coughlin (2017) | 2.99 |
| Y138F | Conter (2006) | -0.77 |
| T146K | Coughlin (2017) | -5.57 |
| N150T | Kure (2004) | -1.76 |
| G156R | Coughlin (2017) | -2.34 |
| Y161C | Korman (2006) | -0.70 |
| Y164H | Khraim (2017) | 0.92 |
| S169P | Coughlin (2017) | -4.76 |
| G171A | Coughlin (2017) | 2.83 |
| G171W | Conter (2006) | 8.68 |
| L173P | Coughlin (2017) | -6.35 |
| Y179N | Coughlin (2017) | -6.65 |
| M182V | ClinVar - Unpub | -2.43 |
| T187K | Conter (2006) | -5.66 |
| L197V | ClinVar - Unpub | -1.52 |
| D198E | Coughlin (2017) | -2.19 |
| D198V | Yilmaz (2015) | 0.08 |
| A202V | Brunel-Guitton (2011) | -0.31 |
| A203V | Coughlin (2017) | 0.82 |
| L207V | ClinVar - Unpub | -0.41 |
| R212K | Conter (2006) | 0.76 |
| H213N | ClinVar - Unpub | 1.09 |
| R216G | ClinVar - Unpub | -0.26 |
| C225R | Azize (2014) | 4.38 |
| V233A | ClinVar - Unpub | -4.45 |
| Q234R | ClinVar - Unpub | -1.64 |
| R236P | Coughlin (2017) | 1.10 |
| Y266D | Coughlin (2017) | -6.74 |
| P267A | Coughlin (2017) | -4.39 |
| P267L | Coughlin (2017) | -1.34 |
| P267S | Swanson (2015) | -2.21 |
| T269M | Swanson (2015) | 2.01 |

| | | |
|---|---|---|
| G271R | Swanson (2015) | -5.96 |
| A283P | Applegarth (2001) | -0.89 |
| H284P | Coughlin (2017) | -2.01 |
| C291R | Swanson (2015) | -5.35 |
| C291Y | Coughlin (2017) | -7.17 |
| D295Y | Azize (2014) | 3.31 |
| L296F | Coughlin (2017) | -1.69 |
| L296R | Kure (2006) | -0.99 |
| P304L | Swanson (2015) | 0.59 |
| P305T | ClinVar - Unpub | 0.73 |
| G306R | Coughlin (2017) | 2.18 |
| A313P | Toone (2002) | -11.12 |
| Q318E | Swanson (2015) | 2.85 |
| Y326D | Coughlin (2017) | -5.19 |
| G327W | Coughlin (2017) | 1.26 |
| P329L | Coughlin (2017) | 0.93 |
| P329T | Applegarth (2001) | -1.43 |
| F334L | Swanson (2015) | -0.12 |
| R337Q | ClinVar - Unpub | -0.93 |
| P345T | Coughlin (2017) | -6.08 |
| R347S | Swanson (2015) | 0.19 |
| T352R | Coughlin (2017) | -3.14 |
| V360L | Swanson (2015) | -0.57 |
| R362C | Coughlin (2017) | -11.42 |
| R362P | Coughlin (2017) | 6.50 |
| Q366R | Swanson (2015) | -0.49 |
| T367I | ClinVar - Unpub | -1.76 |
| Q370K | Coughlin (2017) | -3.17 |
| H371D | Chang (2008) | 0.64 |
| I372F | Azize (2014) | -1.92 |
| R373Q | Conter (2006) | -0.94 |
| R373W | Swanson (2015) | 0.18 |
| K376E | Coughlin (2017) | -0.16 |
| A377V | Yoon (2012) | -0.54 |
| I381T | Swanson (2015) | -4.86 |
| C382Y | Swanson (2015) | 4.07 |
| T383I | Coughlin (2017) | 2.13 |
| A389V | Dinopoulos (2005) | 0.01 |
| F395I | Coughlin (2017) | -2.04 |
| F395L | Coughlin (2017) | -2.28 |
| G421V | Coughlin (2017) | -2.25 |
| L422I | Coughlin (2017) | -0.82 |
| T437I | Swanson (2015) | 4.86 |
| K439N | Swanson (2015) | -0.48 |
| I440N | Conter (2006) | -4.25 |
| I440T | Nykamp (2017) | -5.44 |
| A453S | ClinVar - Unpub | -0.94 |

| | | |
|---|---|---|
| N459D | Coughlin (2017) | 0.45 |
| R461Q | Swanson (2015) | -1.92 |
| R461W | Swanson (2015) | -1.13 |
| G469D | Coughlin (2017) | 5.24 |
| S471P | Coughlin (2017) | -1.29 |
| D473H | Coughlin (2017) | -0.55 |
| T475I | ClinVar - Unpub | -2.58 |
| E478A | Nykamp (2017) | -1.24 |
| E495Q | ClinVar - Unpub | 0.34 |
| R506T | ClinVar - Unpub | -3.22 |
| P509A | Azize (2014) | -4.87 |
| R515S | Toone (2000) | -2.47 |
| P518Q | ClinVar - Unpub | -2.60 |
| V524L | Narisawa (2012) | -0.37 |
| S527T | Nykamp (2017) | -1.90 |
| T532R | Kure (2006) | 1.11 |
| N533Y | Conter (2006) | 0.83 |
| R536Q | Coughlin (2017) | 0.96 |
| R536W | Coughlin (2017) | 2.52 |
| D545H | Coughlin (2017) | 4.39 |
| S547C | Nykamp (2017) | -6.43 |
| L548P | Swanson (2015) | 0.54 |
| L548V | Bravo-Alonso (2017) | -5.79 |
| S551I | Coughlin (2017) | 2.78 |
| M552V | Meyer (2010) | 3.06 |
| L555R | Coughlin (2017) | -5.64 |
| T559N | Coughlin (2017) | -2.35 |
| M560V | Swanson (2015) | 0.46 |
| L562P | Coughlin (2017) | -3.95 |
| S564I | Kure (1992) | -3.23 |
| K574N | ClinVar - Unpub | -1.09 |
| H580D | Coughlin (2017) | -8.11 |
| H580Y | Swanson (2015) | -10.43 |
| P581R | Coughlin (2017) | -2.53 |
| E597K | Azize (2014) | -1.40 |
| G607S | Kruszka (2014) | 0.60 |
| Y608D | Coughlin (2017) | 5.42 |
| V611G | Kure (2006) | -2.59 |
| P615L | Swanson (2015) | -0.92 |
| N616D | Coughlin (2017) | -1.78 |
| N616Y | Coughlin (2017) | -1.29 |
| G618R | Swanson (2015) | -1.48 |
| Q620R | Coughlin (2017) | 1.62 |
| Y623C | Coughlin (2017) | 0.16 |
| Y623H | Genc (2018) | -10.71 |
| A624D | Coughlin (2017) | -5.21 |
| I629T | Coughlin (2017) | -4.33 |

| | | |
|---|---|---|
| R630P | Swanson (2015) | 1.24 |
| C644F | Conter (2006) | -3.65 |
| P647L | Coughlin (2017) | -3.96 |
| P647R | Loviglio (2016) | -2.20 |
| H651R | Conter (2006) | -6.36 |
| G652E | Swanson (2015) | 0.15 |
| S657N | Swanson (2015) | 0.05 |
| S657R | Coughlin (2017) | 2.75 |
| A661P | Coughlin (2017) | 2.64 |
| M663K | Coughlin (2017) | -4.47 |
| A694P | Kure (2006) | -12.12 |
| T698I | Coughlin (2017) | 4.27 |
| P700A | Toone (2002) | -6.15 |
| S701F | Kure (2006) | 6.50 |
| G704V | Coughlin (2017) | -10.97 |
| N709S | ClinVar - Unpub | -0.78 |
| V713M | Nykamp (2017) | -0.21 |
| C714R | Coughlin (2017) | -8.57 |
| L716H | Coughlin (2017) | -0.97 |
| I717N | Coughlin (2017) | -0.93 |
| I717V | ClinVar - Unpub | -3.97 |
| G722R | Coughlin (2017) | -0.02 |
| G728E | Coughlin (2017) | 7.34 |
| G728R | Swanson (2015) | 6.07 |
| N732K | Conter (2006) | -9.83 |
| A733D | Coughlin (2017) | 4.01 |
| A733V | Swanson (2015) | -2.56 |
| R739H | Dinopoulos (2005) | 4.42 |
| D746E | Coughlin (2017) | 1.92 |
| D746H | Coughlin (2017) | 5.30 |
| H753P | Kure (2006) | -5.76 |
| P759R | Coughlin (2017) | -8.54 |
| H760P | Coughlin (2017) | -1.54 |
| H760Q | Coughlin (2017) | -11.51 |
| H760R | Azize (2014) | -5.55 |
| G761R | Kure (1999) | 5.43 |
| G762R | Toone (2002) | -8.50 |
| G763D | Bravo-Alonso (2017) | -4.03 |
| G763R | Coughlin (2017) | -3.07 |
| P765S | Kure (2006) | -6.88 |
| G766C | Love (2014) | 13.85 |
| G766V | Coughlin (2017) | 11.14 |
| G768E | Coughlin (2017) | 6.64 |
| P769L | Coughlin (2017) | -4.95 |
| P769T | Kure (2006) | -1.42 |
| G771R | Swanson (2015) | -7.05 |
| H775P | Coughlin (2017) | 2.58 |

| | | |
|---|---|---|
| H775R | Conter (2006) | 0.26 |
| H775Y | Conter (2006) | -1.37 |
| A777P | Coughlin (2017) | 0.83 |
| P778A | Nykamp (2017) | -1.83 |
| R790W | Kure (2004) | -0.03 |
| E792D | Coughlin (2017) | -0.18 |
| C795S | ClinVar - Unpub | -1.75 |
| A802E | Korman (2004) | 2.87 |
| A802V | Swanson (2015) | 1.71 |
| A803V | Coughlin (2017) | 2.02 |
| G806R | Swanson (2015) | -2.64 |
| S808I | Swanson (2015) | -5.76 |
| S808R | Swanson (2015) | 9.34 |
| A816T | ClinVar - Unpub | -5.07 |
| Y817C | Swanson (2015) | 0.59 |
| K819E | Coughlin (2017) | 0.34 |
| M820V | Coughlin (2017) | 0.63 |
| T830K | Coughlin (2017) | -10.82 |
| T830M | Conter (2006) | 4.45 |
| A833V | Swanson (2015) | -0.16 |
| N838D | Coughlin (2017) | 0.13 |
| Y839C | Liu (2017) | -0.62 |
| M840K | Nykamp (2017) | -0.86 |
| M840V | Kure (2006) | -10.62 |
| A841P | Conter (2006) | -2.73 |
| T846I | ClinVar - Unpub | -8.43 |
| R849G | ClinVar - Unpub | -4.37 |
| A855V | ClinVar - Unpub | 0.65 |
| G860R | Coughlin (2017) | -3.18 |
| E862K | Swanson (2015) | 2.70 |
| D866H | Coughlin (2017) | 0.31 |
| D880V | Conter (2006) | 3.78 |
| R884G | ClinVar - Unpub | 1.73 |
| L885P | Swanson (2015) | -8.11 |
| G889V | Coughlin (2017) | -4.51 |
| P893L | Swanson (2015) | -4.63 |
| T894A | Lin (2018) | 1.91 |
| M895T | ClinVar - Unpub | 0.58 |
| W897C | Tsuyusaki (2012) | -1.08 |
| V905G | Swanson (2015) | -5.31 |
| P907L | Beijer (2012) | -0.17 |
| T908P | Coughlin (2017) | 2.65 |
| S910L | Coughlin (2017) | 3.60 |
| D917E | Coughlin (2017) | 0.12 |
| F919I | Coughlin (2017) | -0.49 |
| I933F | Swanson (2015) | -3.44 |
| I933N | ClinVar - Unpub | -0.45 |

| | | |
|---|---|---|
| I933T | Coughlin (2017) | -2.68 |
| M947I | Coughlin (2017) | 2.18 |
| P949L | Coughlin (2017) | -8.62 |
| H950R | Swanson (2015) | -3.00 |
| S951Y | Coughlin (2017) | 2.95 |
| S957P | Conter (2006) | 0.16 |
| W960C | Coughlin (2017) | -0.52 |
| R962P | Coughlin (2017) | 0.45 |
| R962Q | Coughlin (2017) | -1.98 |
| R966G | Conter (2006) | -3.52 |
| E967D | Coughlin (2017) | -1.36 |
| E979A | ClinVar - Unpub | -0.57 |
| N980D | Karaca (2015) | -0.69 |
| W983R | ClinVar - Unpub | 5.91 |
| R988Q | Coughlin (2017) | -0.53 |
| R988W | Coughlin (2017) | 0.10 |
| G994R | Swanson (2015) | -3.82 |
| D995N | Coughlin (2017) | -0.93 |
| Q996H | ClinVar - Unpub | -0.27 |
| H997Q | Nykamp (2017) | -0.52 |
| C1002W | Jiang (2018) | 0.51 |
