## Supplemental Table 4 for "Large scale analyses of genotype-phenotype relationships of glycine decarboxylase mutations and neurological disease severity"

| Patient | Gender | Allele 1 | Allele 2 | Protein 1 | Protein 2 |
|---|---|---|---|---|---|
| 1 | F | c.1525C>G | c.1789G>A | P509A | E597K |
| 2 | F | c.605C>T | c.2280C>G | A202V | H760Q |
| 3 | M | c.883G>T | c.1607G>A | D295Y | R536Q |
| 4 | M | c.673T>C | c.673T>C | C225R | C225R |
| 5 | M | c.1114A>T | c.1114A>T | I372F | I372F |
| 6 | F | c.2311G>A | c.2720C>T | G771R | P907L |
| 7 | M | c.872G>A | c.872G>A | C291Y | C291Y |
| 8 | M | c.872G>A | c.872G>A | C291Y | C291Y |
| 9 | F | c.605C>T | c.2665+1G>C | A202V | IVS22+1G>C |
| 10 | M | c.605C>T | c.2665+1G>C | A202V | IVS22+1G>C |
| 11 | M | c.2405C>T | c.2665+1G>A | A802V | IVS22+1G>A |
| 12 | M | c.2405C>T | c.2665+1G>A | A802V | IVS22+1G>A |
| 13 | F | c.1166C>T | c.1580+2T>G | A389V | IVS12+2T>G |
| 14 | F | c.1166C>T | c.1580+2T>G | A389V | IVS12+2T>G |
| 15 | M | c.2111G>T | c.2111G>T | G704V | G704V |
| 16 | F | c.2798T>C | c.2798T>C | I933T | I933T |
| 17 | M | c.1084C>T | c.1084C>T | R362C | R362C |
| 18 | F | c.1742C>G | c.437C>A | P581R | T146K |
| 19 | M | c.2980G>A | c.600delG | G994R | G200fs_230 |
| 20 | F | c.1738C>T | c.(?_-1)_(470+1_471-1) | H580Y | delGLDC |
| 21 | M | c.2288G>A | c.1381C>T | G763D | R461W |
| 22 | F | c.2596G>C | c.799C>G | D866H | P267A |
| 23 | M | c.2303G>A | c.(?_-1)_(1707+1_1708-1)del | G768E | del |
| 24 | F | c.1643T>C | c.(?_-1)_(470+1_471-1) | L548P | del |
| 25 | M | c.1742C>G | c.2368C>T | P581R | R790W |
| 26 | F | c.799C>G | c.1126A>G | P267A | K376E |
| 27 | F | c.1117C>T | c.(?_-1)_(334+1_335-1) | R373W | del |
| 28 | F | c.1117C>T | c.3G>T | R373W | M1I |
| 29 | F | c.2714T>G | c.1666G>T | V905G | G556X |
| 30 | F | c.518T>C | c.924delT | L173P | stop |
| 31 | M | c.1111C>G | delGLDC | H371D | del GLDC |
| 32 | M | c.1952A>G | c.1952A>G | H651R | H651R |
| 33 | F | c.560C>A | c.560C>A | T187K | T187K |
| 34 | M | c.413A>T | c.457G>T | Y138F | E153X |
| 35 | M | c.1597A>T | c.2639A>T | N533Y | D880V |
| 36 | F | c.512G>C | c.1285_1286insCAAA | G171A | L429fs |
| 37 | F | c.395C>G | c.1382G>A | S132W | R461Q |
| 38 | M | c.1931G>T | c.2315-1G>A | C644F | missplice |
| 39 | F | c.2521G>C | c.2422delA | A841P | S808fs |
| 40 | F | c.1319T>A | c.1175delC | I440N | A392fs |
| 41 | M | c.2324A>G | c.1009C>T | H775R | R337X |
| 42 | F | c.1117C>T | del Exon 1 | R373W | delExon1 |
| 43 | F | c.2196T>A | c.2838+5G>A | N732K | missplice |

| | | | | | |
|---|---|---|---|---|---|
| 44 | M | c.1166C>T | c.1166C>T | A389V | A389V |
| 45 | M | c.1166C>T | c.1166C>T | A389V | A389V |
| 46 | F | c.2405C>T | c.2405C>T | A802V | A802V |
| 47 | F | c.2405C>T | c.2405C>T | A802V | A802V |
| 48 | F | c.2405C>T | c.2405C>T | A802V | A802V |
| 49 | F | c.2405C>T | c.2405C>T | A802V | A802V |
| 50 | M | c.2216G>A | c.2216G>A | R739H | R739H |
| 51 | F | c.449A>C | c.2368C>T | N150T | R790W |
| 52 | F | c.245T>G | c.1821_1831del | L82W | 607fs |
| 53 | ? | c.1867T>C | c.1867T>C | Y623H | Y623H |
| 54 | M | c.3006C>G | c.1256C>G | C1002W | S419X |
| 55 | F | c.3006C>G | c.1256C>G | C1002W | S419X |
| 56 | M | c.1859A>G | del Exon 3-9 | Q620R | del Exon 3-9 |
| 57 | M | c.395C>T | c.256-?_334þ?del | S132L | S86Vfs_119 |
| 58 | M | c.490T>C | c.490T>C | Y164H | Y164H |
| 59 | F | c.490T>C | c.490T>C | Y164H | Y164H |
| 60 | F | c.490T>C | c.490T>C | Y164H | Y164H |
| 61 | F | c.482A>G | c.482A>G | Y161C | Y161C |
| 62 | M | c.482A>G | c.482A>G | Y161C | Y161C |
| 63 | F | c.2963G>A | c.2963G>A | R988Q | R988Q |
| 64 | M | c.2963G>A | c.2963G>A | R988Q | R988Q |
| 65 | M | c.2963G>A | c.2963G>A | R988Q | R988Q |
| 66 | M | c.2680A>G | del Exon3 | T894A | del Exon3 |
| 67 | M | c.2680A>G | del Exon3 | T894A | del Exon3 |
| 68 | F | c.2680A>G | del Exon3 | T894A | del Exon3 |
| 69 | M | c.2516A>G | c.2457+2T>A | Y839C | splice |
| 70 | F | c.2296G>T | c.2296G>T | G766C | G766C |
| 71 | M | c.2296G>T | c.2296G>T | G766C | G766C |
| 72 | M | c.2846C>T | c.2846C>T | P949L | P949L |
| 73 | F | c.2311G>A | c.1654A>G | G771R | M552V |
| 74 | F | c.2216G>A | del Exon 1-2 | R739H | del Exon 1-2 |
| 75 | M | c.1742C>G | c.1742C>G | P581R | P581R |
| 76 | M | c.1382G>A | del GLDC | R461Q | del GLDC |
| 77 | M | c.605C>T | c.605C>T | A202V | A202V |
| 78 | M | c.2849A>G | c.2849A>G | H950R | H950R |
| 79 | M | c.1097A>G | c.1097A>G | Q366R | Q366R |
| 80 | M | c.1381C>T | c.1381C>T | R461W | R461W |
| 81 | F | c.1545G>C | c.1545G>C | R515S | R515S |
| 82 | F | c.395C>T | c.395C>T | S132L | S132L |
| 83 | M | c.806C>T | c.806C>T | T269M | T269M |
| 84 | F | c.2422A>C | c.952C>G | S808R | Q318E |
| 85 | F | c.2281G>C | c.1869T>G | G761R | Y632X |
| 86 | M | c.1545G>C | c.2316-1G>A | R515S | IVS19-1G>A |
| 87 | M | c.395C>T | c.499G>T | S132L | E167X |
| 88 | F | c.1969A>C | c.2405C>T | S657R | A802V |
| 89 | F | c.2720C>T | del Exon 1-24 | P907L | del Exon 1-24 |
| 90 | F | c.1545G>C | c.1852G>A | R515S | G618R |

| | | | | | |
|---|---|---|---|---|---|
| 91 | F | c.2198C>T | c.2316-1G>A | A733V | IVS19-1G>A |
| 92 | M | c.395C>T | c.2423G>T | S132L | S808I |
| 93 | F | c.2416G>C | c.2481_2484delACAA | G806R | K827Kfs*7 |
| 94 | M | c.2980G>A | c.2584G>A | G994R | E862K |
| 95 | M | c.1545G>C | c.2316-1G>A | R515S | IVS19-1G>A |
| 96 | F | c.800C>T | c.2415G>A | P267L | W805X |
| 97 | F | c.2654T>C | c.2019T>A | L885P | Y637X |
| 98 | M | c.2311G>A | c.2316-1G>A | G771R | IVS19-1G>A |
| 99 | F | c.395C>T | c.1310C>T | S132L | T437I |
| 100 | F | c.1000T>C | del Exon 1-25 | F334L | del GLDC |
| 101 | F | c.1166C>T | c.1545G>C | A389V | R515S |
| 102 | M | c.1642C>G | del Exon 1-25 | L548V | del GLDC |
| 103 | M | c.2405C>T | c.2665+1G>C | A802V | IVS22+1G>C |
| 104 | F | c.2405C>T | c.1545G>C | A802V | R515S |
| 105 | M | c.1166C>T | c.1545G>C | A389V | R515S |
| 106 | F | c.1142T>C | c.1382G>A | I381T | R461Q |
| 107 | M | c.847G>C | c.1382G>A | A283P | R461Q |
| 108 | M | c.847G>C | c.1382G>A | A283P | R461Q |
| 109 | M | c.1382G>A | c.1929-2A>G | R461Q | IVS12+2T>G |
| 110 | M | c.482A>G | c.1041A>C | Y161C | R347S |
| 111 | M | c.466G>C | c.2183G>A | G156R | G728E |
| 112 | M | c.1889G>C | c.1642C>G | R630P | L548V |
| 113 | F | c.2405C>A | c.2315+2T>G | A802E | IVS19+2T>G |
| 114 | M | c.2714T>G | c.2183G>A | V905G | G728E |
| 115 | F | c.1955G>A | c.1118G>A | G652E | R373Q |
| 116 | M | c.2498C>T | delExon1-2 | A833V | del Exon 1-2 |
| 117 | M | c.2422A>C | c.538C>T | S808R | Q180X |
| 118 | M | c.1545G>C | delExon1-2 | R515S | del Exon 1-2 |
| 119 | F | c.1545G>C | delExon1-2 | R515S | del Exon 1-2 |
| 120 | F | c.806C>T | delExon1-2 | T269M | del Exon 1-2 |
| 121 | F | c.806C>T | delExon1-2 | T269M | del Exon 1-2 |
| 122 | F | c.2183G>A | c.1545G>C | G728E | R515S |
| 123 | F | c.1317G>T | delExon1-21 | K439N | del Exon 1-21 |
| 124 | M | c.1819G>A | delExon3-9 | G607S | del Exon 3-9 |
| 125 | F | c.1545G>C | c.2665+1G>C | R515S | IVS22+1G>C |
| 126 | F | c.2450A>G | delExon1 | Y817C | del Exon 1 |
| 127 | F | c.2678C>T | c.1545G>C | P893L | R515S |
| 128 | F | c.1678A>G | c.499G>T | M560V | E167X |
| 129 | F | c.2654T>C | c.2691G>T | L885P | W897C |
| 130 | M | c.593A>T | c.593A>T | D198V | D198V |
| 131 | M | c.1130C>T | c.2081_2088del | A377V | A694Dfs |

| Reference | Death? | Seizures | Cognition | Brain Malformations | Muscle/ Movement Control | COS |
|---|---|---|---|---|---|---|
| Azize 2014 | no | 1 | - | 3 | 2 | **6** |
| Azize 2014 | yes | N/A | N/A | N/A | N/A | **N/A** |
| Azize 2014 | no | 3 | - | - | 3 | **6** |
| Azize 2014 | yes | N/A | N/A | N/A | N/A | **N/A** |
| Azize 2014 | no | 3 | - | 3 | 3 | **9** |
| Beijer 2012 | yes | N/A | N/A | N/A | N/A | **N/A** |
| Bjoraker 2016 | yes | N/A | N/A | N/A | N/A | **N/A** |
| Bjoraker 2016 | no | 1 | 1 | - | - | **2** |
| Bjoraker 2016 | no | 1 | 1 | - | - | **2** |
| Bjoraker 2016 | no | 1 | 1 | - | - | **2** |
| Bjoraker 2016 | no | 1 | 1 | - | - | **2** |
| Bjoraker 2016 | no | 1 | 1 | - | - | **2** |
| Bjoraker 2016 | no | 1 | 1 | - | - | **2** |
| Bjoraker 2016 | no | 1 | 1 | - | - | **2** |
| Bravo-Alonso 2018 | yes | N/A | N/A | N/A | N/A | **N/A** |
| Bravo-Alonso 2018 | yes | N/A | N/A | N/A | N/A | **N/A** |
| Bravo-Alonso 2018 | no | 1 | 1 | - | - | **2** |
| Bravo-Alonso 2018 | yes | N/A | N/A | N/A | N/A | **N/A** |
| Bravo-Alonso 2018 | yes | N/A | N/A | N/A | N/A | **N/A** |
| Bravo-Alonso 2018 | yes | N/A | N/A | N/A | N/A | **N/A** |
| Bravo-Alonso 2018 | yes | N/A | N/A | N/A | N/A | **N/A** |
| Bravo-Alonso 2018 | yes | N/A | N/A | N/A | N/A | **N/A** |
| Bravo-Alonso 2018 | yes | N/A | N/A | N/A | N/A | **N/A** |
| Bravo-Alonso 2018 | yes | N/A | N/A | N/A | N/A | **N/A** |
| Bravo-Alonso 2018 | yes | N/A | N/A | N/A | N/A | **N/A** |
| Bravo-Alonso 2018 | no | 1 | 2 | - | - | **3** |
| Bravo-Alonso 2018 | no | 1 | 1 | - | - | **2** |
| Bravo-Alonso 2018 | no | 1 | 1 | - | - | **2** |
| Bravo-Alonso 2018 | yes | N/A | N/A | N/A | N/A | **N/A** |
| Bravo-Alonso 2018 | yes | N/A | N/A | N/A | N/A | **N/A** |
| Chang 2008 | no | 1 | - | 3 | 3 | **7** |
| Conter 2006 | yes | N/A | N/A | N/A | N/A | **N/A** |
| Conter 2006 | no | - | 3 | - | - | **INS** |
| Conter 2006 | yes | N/A | N/A | N/A | N/A | **N/A** |
| Conter 2006 | no | - | 2 | - | - | **INS** |
| Conter 2006 | yes | N/A | N/A | N/A | N/A | **N/A** |
| Conter 2006 | yes | N/A | N/A | N/A | N/A | **N/A** |
| Conter 2006 | yes | N/A | N/A | N/A | N/A | **N/A** |
| Conter 2006 | yes | N/A | N/A | N/A | N/A | **N/A** |
| Conter 2006 | no | - | - | - | - | **INS** |
| Conter 2006 | no | - | - | - | - | **INS** |
| Conter 2006 | yes | N/A | N/A | N/A | N/A | **N/A** |
| Conter 2006 | yes | N/A | N/A | N/A | N/A | **N/A** |

| | | | | | | |
|---|---|---|---|---|---|---|
| Dinopoulos 2005 | no | - | 1 | - | 2 | **3** |
| Dinopoulos 2005 | no | 1 | 2 | - | 2 | **5** |
| Dinopoulos 2005 | no | 0 | 0 | - | 2 | **2** |
| Dinopoulos 2005 | no | 0 | 0 | - | 0 | **0** |
| Dinopoulos 2005 | no | 0 | 0 | - | 0 | **0** |
| Dinopoulos 2005 | no | 0 | 1 | - | 2 | **3** |
| Dinopoulos 2005 | no | 1 | 1 | - | 2 | **4** |
| Dinopoulos 2005 | no | 1 | 3 | 0 | 2 | **6** |
| Dinopoulos 2005 | no | 1 | 1 | 0 | 2 | **4** |
| Genc 2018 | no | 1 | 1 | - | 1 | **3** |
| Jiang 2017 | no | 3 | 2 | - | - | **5** |
| Jiang 2017 | no | - | 2 | - | 2 | **4** |
| Kanekar 2013 | no | 3 | - | 0 | 3 | **6** |
| Kava 2018 | no | 3 | - | 3 | 2 | **8** |
| Khraim 2017 | no | 1 | 2 | 0 | 2 | **5** |
| Khraim 2017 | no | 1 | 2 | - | 1 | **4** |
| Khraim 2017 | no | 1 | 1 | - | 0 | **2** |
| Korman 2006 | yes | N/A | N/A | N/A | N/A | **N/A** |
| Korman 2006 | no | 3 | 3 | 3 | 3 | **12** |
| Kose 2017 | no | 1 | - | 3 | 3 | **7** |
| Kose 2017 | yes | N/A | N/A | N/A | N/A | **N/A** |
| Kose 2017 | yes | N/A | N/A | N/A | N/A | **N/A** |
| Lin 2018 | yes | N/A | N/A | N/A | N/A | **N/A** |
| Lin 2018 | yes | N/A | N/A | N/A | N/A | **N/A** |
| Lin 2018 | no | 3 | 3 | - | 2 | **8** |
| Liu 2017 | no | 3 | - | 3 | 3 | **9** |
| Love 2014 | yes | N/A | N/A | N/A | N/A | **N/A** |
| Love 2014 | yes | N/A | N/A | N/A | N/A | **N/A** |
| McAdams 2009 | no | 3 | 3 | 3 | - | **9** |
| Meyer 2010 | no | 1 | - | - | 2 | **3** |
| Morrison 2006 | no | 1 | 1 | 0 | 1 | **3** |
| Rodríguez-Benítez 2014 | no | 1 | 3 | 3 | - | **7** |
| Suzuki 2010 | no | 1 | 2 | 3 | 2 | **8** |
| Swanson 2015 | no | 1 | 1 | - | - | **2** |
| Swanson 2015 | no | 3 | 3 | 3 | - | **9** |
| Swanson 2015 | no | 1 | 2 | - | - | **3** |
| Swanson 2015 | no | - | 3 | - | - | **INS** |
| Swanson 2015 | no | 3 | 3 | 3 | - | **9** |
| Swanson 2015 | no | - | 3 | - | - | **INS** |
| Swanson 2015 | no | 1 | 1 | - | - | **2** |
| Swanson 2015 | yes | N/A | N/A | N/A | N/A | **N/A** |
| Swanson 2015 | no | 3 | 3 | 3 | - | **9** |
| Swanson 2015 | no | 1 | 3 | 3 | - | **7** |
| Swanson 2015 | no | 3 | 3 | 3 | - | **9** |
| Swanson 2015 | no | - | 3 | - | - | **INS** |
| Swanson 2015 | no | - | 3 | 3 | - | **6** |
| Swanson 2015 | no | - | 3 | 3 | - | **6** |

| | | | | | | |
|---|---|---|---|---|---|---|
| Swanson 2015 | no | 1 | 3 | - | - | **4** |
| Swanson 2015 | no | - | 3 | - | - | **INS** |
| Swanson 2015 | no | - | 3 | - | - | **INS** |
| Swanson 2015 | no | - | 3 | - | - | **INS** |
| Swanson 2015 | no | - | 3 | 3 | - | **6** |
| Swanson 2015 | no | - | 3 | - | - | **INS** |
| Swanson 2015 | no | - | 3 | 3 | - | **6** |
| Swanson 2015 | no | - | 3 | 3 | - | **6** |
| Swanson 2015 | no | - | 3 | - | - | **INS** |
| Swanson 2015 | no | - | 3 | 3 | - | **6** |
| Swanson 2015 | no | - | 3 | 3 | - | **6** |
| Swanson 2015 | no | 1 | 2 | - | - | **3** |
| Swanson 2015 | no | 1 | 2 | - | - | **3** |
| Swanson 2015 | no | 1 | 2 | - | - | **3** |
| Swanson 2015 | no | 3 | 1 | - | - | **4** |
| Swanson 2015 | no | 1 | 1 | - | - | **2** |
| Swanson 2015 | no | 3 | 1 | - | - | **4** |
| Swanson 2015 | no | 1 | 1 | - | - | **2** |
| Swanson 2015 | no | 1 | 1 | - | - | **2** |
| Swanson 2015 | no | 1 | 1 | - | - | **2** |
| Swanson 2015 | no | 1 | 1 | - | - | **2** |
| Swanson 2015 | no | 1 | 1 | - | - | **2** |
| Swanson 2015 | no | 1 | 1 | - | - | **2** |
| Swanson 2015 | no | 1 | 1 | - | - | **2** |
| Swanson 2015 | no | 1 | 1 | - | - | **2** |
| Swanson 2015 | yes | N/A | N/A | N/A | N/A | **N/A** |
| Swanson 2015 | yes | N/A | N/A | N/A | N/A | **N/A** |
| Swanson 2015 | yes | N/A | N/A | N/A | N/A | **N/A** |
| Swanson 2015 | yes | N/A | N/A | N/A | N/A | **N/A** |
| Swanson 2015 | yes | N/A | N/A | N/A | N/A | **N/A** |
| Swanson 2015 | yes | N/A | N/A | N/A | N/A | **N/A** |
| Swanson 2015 | yes | N/A | N/A | N/A | N/A | **N/A** |
| Swanson 2015 | yes | N/A | N/A | N/A | N/A | **N/A** |
| Swanson 2015 | yes | N/A | N/A | N/A | N/A | **N/A** |
| Swanson 2015 | yes | N/A | N/A | N/A | N/A | **N/A** |
| Swanson 2015 | yes | N/A | N/A | N/A | N/A | **N/A** |
| Swanson 2015 | yes | N/A | N/A | N/A | N/A | **N/A** |
| Swanson 2015 | yes | N/A | N/A | N/A | N/A | **N/A** |
| Tsuyusaki 2012 | no | 3 | 3 | 3 | 3 | **12** |
| Yilmaz 2015 | no | 3 | - | 3 | 3 | **9** |
| Yoon 2012 | no | 3 | 3 | 0 | 2 | **8** |
