## Supplemental Table 5 for "Large scale analyses of genotype-phenotype relationships of glycine decarboxylase mutations and neurological disease severity"

| MUT | Conservation of Amino Acid | Helix Mutation | Sheet Mutation | Destabilizing | Δ Proline | C-terminal mutation | Active Site Mutation | Δ Polarity | Δ Aromaticity | Δ Charge | Conservation of Substition | Δ Codon | Δ Size | H-interface Mutation | N-PLP Mutation | Active Region | Dimerazation Interface | Stabilizing Mutation | MMS |
|---|---|---|---|---|---|---|---|---|---|---|---|---|---|---|---|---|---|---|---|
| **Pathogenic Mutations** | | | | | | | | | | | | | | | | | | | |
| D36H | 0 | 0 | 0 | 0 | 0 | 0 | 0 | 0 | 1 | 1 | 0 | 0 | 0 | 0 | 0 | 0 | 0 | 0 | 2 |
| S37N | 0 | 0 | 0 | 0 | 0 | 0 | 0 | 0 | 0 | 0 | -1 | 0 | 0 | 0 | 0 | 0 | 0 | 0 | 0 |
| G48W | 0 | 0 | 0 | 0 | 0 | 0 | 0 | 1 | 1 | 0 | 0 | 0 | 1 | 0 | 0 | 0 | 0 | 0 | 3 |
| R59T | 0 | 0 | 0 | 1 | 0 | 0 | 0 | 0 | 0 | 1 | 0 | 0 | 1 | 0 | 0 | 0 | 1 | 0 | 4 |
| P70R | 0 | 0 | 0 | 0 | 1 | 0 | 0 | 1 | 0 | 1 | 0 | 0 | 1 | 0 | 0 | 0 | 0 | 1 | 5 |
| L79P | 1 | 1 | 0 | 1 | 1 | 0 | 0 | 0 | 0 | 0 | 0 | 0 | 1 | 0 | 0 | 0 | 0 | 0 | 5 |
| L82S | 0 | 1 | 0 | 1 | 0 | 0 | 0 | 1 | 0 | 0 | 0 | 0 | 1 | 0 | 0 | 0 | 0 | 0 | 4 |
| L82W | 0 | 1 | 0 | 0 | 0 | 0 | 0 | 1 | 1 | 0 | 0 | 0 | 1 | 0 | 0 | 0 | 0 | 0 | 4 |
| D88N | 0 | 1 | 0 | 0 | 0 | 0 | 0 | 0 | 0 | 1 | -1 | 0 | 0 | 0 | 0 | 0 | 0 | 0 | 1 |
| L90F | 1 | 1 | 0 | 1 | 0 | 0 | 0 | 0 | 1 | 0 | -1 | 0 | 0 | 0 | 0 | 0 | 0 | 0 | 3 |
| P110H | 0 | 0 | 0 | 1 | 1 | 0 | 0 | 1 | 1 | 1 | 0 | 1 | 0 | 0 | 0 | 0 | 0 | 0 | 6 |
| S132L | 1 | 0 | 1 | 0 | 0 | 0 | 0 | 1 | 0 | 0 | 0 | 0 | 1 | 0 | 0 | 0 | 0 | 0 | 4 |
| S132W | 1 | 0 | 1 | 0 | 0 | 0 | 0 | 0 | 1 | 0 | 0 | 0 | 1 | 0 | 0 | 0 | 0 | 1 | 5 |
| G135A | 1 | 0 | 0 | 1 | 0 | 0 | 0 | 0 | 0 | 0 | -1 | 1 | 0 | 0 | 0 | 0 | 0 | 0 | 2 |
| Y138D | 1 | 0 | 0 | 0 | 0 | 0 | 0 | 0 | 1 | 1 | 0 | 0 | 1 | 0 | 1 | 0 | 0 | 0 | 5 |
| Y138F | 1 | 0 | 0 | 0 | 0 | 0 | 0 | 1 | 0 | 0 | -1 | 0 | 0 | 0 | 1 | 0 | 0 | 0 | 2 |
| T146K | 1 | 0 | 0 | 1 | 0 | 0 | 0 | 0 | 0 | 1 | 0 | 0 | 1 | 0 | 0 | 0 | 1 | 0 | 5 |
| N150T | 1 | 1 | 0 | 1 | 0 | 0 | 0 | 0 | 0 | 0 | -1 | 1 | 0 | 0 | 0 | 0 | 1 | 0 | 4 |
| G156R | 0 | 1 | 0 | 1 | 0 | 0 | 0 | 1 | 0 | 1 | 0 | 0 | 1 | 0 | 0 | 0 | 0 | 0 | 5 |
| Y161C | 1 | 0 | 0 | 0 | 0 | 0 | 1 | 0 | 1 | 0 | 0 | 0 | 1 | 0 | 0 | 0 | 0 | 0 | 4 |
| Y164H | 1 | 0 | 0 | 0 | 0 | 0 | 1 | 0 | 0 | 1 | -1 | 0 | 0 | 0 | 0 | 0 | 0 | 0 | 2 |
| S169P | 1 | 0 | 0 | 1 | 1 | 0 | 0 | 1 | 0 | 0 | 0 | 0 | 0 | 0 | 0 | 0 | 0 | 0 | 4 |
| G171A | 1 | 1 | 0 | 0 | 0 | 0 | 0 | 0 | 0 | 0 | -1 | 0 | 0 | 0 | 0 | 0 | 0 | 0 | 1 |
| G171W | 1 | 1 | 0 | 0 | 0 | 0 | 0 | 1 | 1 | 0 | 0 | 0 | 1 | 0 | 0 | 0 | 0 | 1 | 6 |
| L173P | 1 | 1 | 0 | 1 | 1 | 0 | 0 | 0 | 0 | 0 | 0 | 0 | 1 | 0 | 0 | 0 | 0 | 0 | 5 |
| Y179N | 0 | 1 | 0 | 1 | 0 | 0 | 0 | 0 | 1 | 0 | 0 | 0 | 1 | 0 | 0 | 0 | 0 | 0 | 4 |
| M182V | 0 | 1 | 0 | 1 | 0 | 0 | 0 | 0 | 0 | 0 | -1 | 0 | 0 | 0 | 0 | 0 | 0 | 0 | 1 |
| T187K | 1 | 1 | 0 | 1 | 0 | 0 | 0 | 0 | 0 | 1 | 0 | 0 | 1 | 0 | 0 | 0 | 0 | 0 | 5 |
| L197V | 1 | 0 | 0 | 1 | 0 | 0 | 0 | 0 | 0 | 0 | -1 | 0 | 0 | 0 | 0 | 0 | 0 | 0 | 1 |
| D198E | 1 | 0 | 0 | 1 | 0 | 0 | 0 | 0 | 0 | 0 | -1 | 1 | 0 | 0 | 1 | 0 | 0 | 0 | 3 |
| D198V | 1 | 0 | 0 | 0 | 0 | 0 | 0 | 1 | 0 | 1 | 0 | 1 | 0 | 0 | 1 | 0 | 0 | 0 | 5 |
| A202V | 1 | 1 | 0 | 0 | 0 | 0 | 0 | 0 | 0 | 0 | -1 | 0 | 1 | 0 | 0 | 0 | 0 | 0 | 2 |
| A203V | 1 | 1 | 0 | 0 | 0 | 0 | 0 | 0 | 0 | 0 | -1 | 0 | 1 | 0 | 0 | 0 | 0 | 0 | 2 |
| L207V | 1 | 1 | 0 | 0 | 0 | 0 | 0 | 0 | 0 | 0 | -1 | 0 | 0 | 0 | 0 | 0 | 0 | 0 | 1 |
| R212K | 0 | 0 | 0 | 0 | 0 | 0 | 0 | 0 | 0 | 0 | -1 | 0 | 0 | 1 | 0 | 0 | 0 | 0 | 0 |
| H213N | 0 | 0 | 0 | 0 | 0 | 0 | 0 | 0 | 1 | 1 | -1 | 0 | 0 | 0 | 0 | 0 | 0 | 0 | 1 |
| R216G | 0 | 0 | 0 | 0 | 0 | 0 | 0 | 1 | 0 | 1 | 0 | 0 | 1 | 0 | 0 | 0 | 0 | 0 | 3 |
| C225R | 0 | 0 | 0 | 0 | 0 | 0 | 0 | 0 | 0 | 1 | 0 | 1 | 1 | 0 | 1 | 0 | 0 | 1 | 5 |
| V233A | 0 | 1 | 0 | 1 | 0 | 0 | 0 | 1 | 0 | 0 | -1 | 0 | 1 | 0 | 0 | 0 | 0 | 0 | 3 |
| Q234R | 0 | 1 | 0 | 1 | 0 | 0 | 0 | 0 | 0 | 1 | -1 | 0 | 0 | 0 | 0 | 0 | 0 | 0 | 2 |
| R236P | 1 | 1 | 0 | 0 | 1 | 0 | 0 | 1 | 0 | 1 | 0 | 0 | 1 | 0 | 0 | 0 | 0 | 0 | 6 |
| Y266D | 0 | 0 | 1 | 1 | 0 | 0 | 0 | 0 | 1 | 1 | 0 | 0 | 1 | 0 | 1 | 0 | 0 | 0 | 6 |
| P267A | 1 | 0 | 0 | 1 | 1 | 0 | 0 | 0 | 0 | 0 | 0 | 0 | 0 | 0 | 1 | 0 | 0 | 0 | 4 |
| P267L | 1 | 0 | 0 | 0 | 1 | 0 | 0 | 0 | 0 | 0 | 0 | 0 | 1 | 0 | 1 | 0 | 0 | 0 | 4 |
| P267S | 1 | 0 | 0 | 1 | 1 | 0 | 0 | 1 | 0 | 0 | 0 | 0 | 0 | 0 | 1 | 0 | 0 | 0 | 5 |
| T269M | 0 | 0 | 0 | 0 | 0 | 0 | 0 | 1 | 0 | 0 | 0 | 0 | 0 | 0 | 0 | 0 | 0 | 0 | 1 |
| G271R | 1 | 0 | 0 | 1 | 0 | 0 | 0 | 1 | 0 | 1 | 0 | 0 | 1 | 0 | 0 | 0 | 0 | 0 | 5 |
| A283P | 0 | 1 | 0 | 0 | 1 | 0 | 0 | 0 | 0 | 0 | 0 | 0 | 0 | 0 | 0 | 0 | 0 | 0 | 2 |
| H284P | 0 | 1 | 0 | 1 | 1 | 0 | 0 | 1 | 1 | 1 | 0 | 1 | 0 | 0 | 0 | 0 | 0 | 0 | 7 |
| C291R | 0 | 0 | 1 | 1 | 0 | 0 | 0 | 0 | 0 | 1 | 0 | 1 | 1 | 0 | 0 | 0 | 0 | 0 | 5 |
| C291Y | 0 | 0 | 1 | 1 | 0 | 0 | 0 | 0 | 1 | 0 | 0 | 0 | 1 | 0 | 0 | 0 | 0 | 0 | 4 |
| D295Y | 1 | 0 | 0 | 0 | 0 | 0 | 0 | 0 | 1 | 1 | 0 | 0 | 1 | 0 | 0 | 0 | 0 | 1 | 5 |
| L296F | 0 | 0 | 0 | 1 | 0 | 0 | 0 | 0 | 1 | 0 | -1 | 1 | 0 | 0 | 0 | 0 | 0 | 0 | 2 |
| L296R | 0 | 0 | 0 | 0 | 0 | 0 | 0 | 1 | 0 | 1 | 0 | 0 | 0 | 0 | 0 | 0 | 0 | 0 | 2 |
| P304L | 0 | 0 | 0 | 0 | 1 | 0 | 0 | 0 | 0 | 0 | 0 | 0 | 1 | 0 | 0 | 0 | 0 | 0 | 2 |
| P305T | 1 | 1 | 0 | 0 | 1 | 0 | 0 | 1 | 0 | 0 | 0 | 0 | 0 | 0 | 0 | 0 | 0 | 0 | 4 |
| G306R | 1 | 1 | 0 | 0 | 0 | 0 | 0 | 1 | 0 | 1 | 0 | 0 | 1 | 0 | 0 | 0 | 0 | 0 | 5 |
| A313P | 0 | 0 | 1 | 1 | 1 | 0 | 0 | 0 | 0 | 0 | 0 | 0 | 0 | 0 | 0 | 0 | 0 | 0 | 3 |
| Q318E | 1 | 0 | 0 | 0 | 0 | 0 | 0 | 0 | 0 | 1 | -1 | 0 | 0 | 0 | 0 | 0 | 0 | 0 | 1 |
| Y326D | 0 | 1 | 0 | 1 | 0 | 0 | 0 | 0 | 1 | 1 | 0 | 0 | 1 | 0 | 0 | 0 | 0 | 0 | 5 |
| G327W | 1 | 1 | 0 | 0 | 0 | 0 | 0 | 1 | 1 | 0 | 0 | 0 | 1 | 0 | 0 | 0 | 0 | 0 | 5 |
| P329L | 1 | 0 | 0 | 0 | 1 | 0 | 0 | 0 | 0 | 0 | 0 | 0 | 1 | 0 | 0 | 0 | 0 | 0 | 3 |
| P329T | 1 | 0 | 0 | 0 | 1 | 0 | 0 | 1 | 0 | 0 | 0 | 0 | 0 | 0 | 0 | 0 | 0 | 0 | 3 |
| F334L | 0 | 0 | 1 | 0 | 0 | 0 | 0 | 0 | 1 | 0 | -1 | 1 | 0 | 0 | 0 | 0 | 0 | 0 | 2 |
| R337Q | 0 | 0 | 0 | 0 | 0 | 0 | 0 | 0 | 0 | 1 | -1 | 0 | 0 | 0 | 0 | 0 | 0 | 0 | 0 |
| P345T | 1 | 0 | 0 | 1 | 1 | 0 | 0 | 1 | 0 | 0 | 0 | 0 | 0 | 0 | 0 | 0 | 0 | 0 | 4 |
| R347S | 1 | 0 | 0 | 0 | 0 | 0 | 0 | 0 | 0 | 1 | 0 | 0 | 1 | 0 | 0 | 0 | 0 | 0 | 3 |
| T352R | 1 | 0 | 1 | 1 | 0 | 0 | 0 | 0 | 0 | 1 | 0 | 0 | 1 | 0 | 0 | 0 | 0 | 0 | 5 |
| V360L | 1 | 0 | 1 | 0 | 0 | 0 | 0 | 0 | 0 | 0 | -1 | 0 | 0 | 0 | 0 | 0 | 0 | 0 | 1 |
| R362C | 1 | 0 | 0 | 1 | 0 | 0 | 0 | 0 | 0 | 1 | 0 | 1 | 1 | 0 | 0 | 0 | 0 | 0 | 5 |
| R362P | 1 | 0 | 0 | 0 | 1 | 0 | 0 | 1 | 0 | 1 | 0 | 0 | 1 | 0 | 0 | 0 | 0 | 1 | 6 |
| Q366R | 1 | 0 | 0 | 0 | 0 | 0 | 0 | 0 | 0 | 1 | -1 | 0 | 0 | 1 | 0 | 0 | 0 | 0 | 2 |
| T367I | 1 | 0 | 0 | 1 | 0 | 0 | 0 | 1 | 0 | 0 | 0 | 1 | 1 | 1 | 0 | 0 | 0 | 0 | 6 |
| Q370K | 1 | 0 | 0 | 1 | 0 | 0 | 0 | 0 | 0 | 1 | -1 | 0 | 0 | 0 | 0 | 0 | 0 | 0 | 2 |
| H371D | 1 | 0 | 0 | 0 | 0 | 0 | 0 | 0 | 1 | 1 | 0 | 0 | 0 | 0 | 0 | 0 | 0 | 0 | 3 |
| I372F | 1 | 0 | 0 | 1 | 0 | 0 | 0 | 0 | 1 | 0 | -1 | 1 | 0 | 1 | 0 | 0 | 0 | 0 | 4 |
| R373Q | 1 | 0 | 0 | 0 | 0 | 0 | 0 | 0 | 0 | 1 | -1 | 0 | 0 | 1 | 0 | 0 | 0 | 0 | 2 |
| R373W | 1 | 0 | 0 | 0 | 0 | 0 | 0 | 0 | 1 | 1 | 0 | 0 | 1 | 1 | 0 | 0 | 0 | 0 | 5 |
| K376E | 1 | 0 | 0 | 0 | 0 | 0 | 0 | 0 | 0 | 1 | -1 | 0 | 0 | 1 | 0 | 0 | 0 | 0 | 2 |
| A377V | 1 | 0 | 0 | 0 | 0 | 0 | 0 | 0 | 0 | 0 | -1 | 0 | 1 | 0 | 0 | 0 | 0 | 0 | 1 |
| I381T | 1 | 0 | 0 | 1 | 0 | 0 | 0 | 1 | 0 | 0 | 0 | 1 | 1 | 0 | 0 | 0 | 0 | 0 | 5 |
| C382Y | 1 | 0 | 0 | 0 | 0 | 0 | 1 | 0 | 1 | 0 | 0 | 0 | 1 | 0 | 0 | 0 | 0 | 1 | 5 |
| T383I | 1 | 0 | 0 | 0 | 0 | 0 | 0 | 1 | 0 | 0 | 0 | 0 | 1 | 0 | 0 | 1 | 0 | 0 | 4 |
| A389V | 1 | 1 | 0 | 0 | 0 | 0 | 0 | 0 | 0 | 0 | -1 | 0 | 1 | 0 | 0 | 0 | 0 | 0 | 2 |
| F395I | 1 | 1 | 0 | 1 | 0 | 0 | 0 | 0 | 1 | 0 | -1 | 1 | 0 | 0 | 0 | 0 | 0 | 0 | 4 |
| F395L | 1 | 1 | 0 | 1 | 0 | 0 | 0 | 0 | 1 | 0 | -1 | 1 | 0 | 0 | 0 | 0 | 0 | 0 | 4 |
| G421V | 0 | 1 | 0 | 1 | 0 | 0 | 0 | 0 | 0 | 0 | 0 | 1 | 1 | 0 | 0 | 0 | 0 | 0 | 4 |
| L422I | 0 | 1 | 0 | 0 | 0 | 0 | 0 | 0 | 0 | 0 | -1 | 1 | 0 | 0 | 0 | 0 | 0 | 0 | 1 |
| T437I | 1 | 0 | 1 | 0 | 0 | 0 | 0 | 1 | 0 | 0 | 0 | 1 | 1 | 0 | 0 | 0 | 0 | 1 | 6 |
| K439N | 0 | 0 | 1 | 0 | 0 | 0 | 0 | 0 | 0 | 1 | -1 | 1 | 1 | 0 | 0 | 0 | 0 | 0 | 3 |
| I440N | 0 | 0 | 1 | 1 | 0 | 0 | 0 | 1 | 0 | 0 | 0 | 1 | 1 | 0 | 0 | 0 | 0 | 0 | 5 |
| I440T | 0 | 0 | 1 | 1 | 0 | 0 | 0 | 1 | 0 | 0 | 0 | 0 | 1 | 0 | 0 | 0 | 0 | 0 | 4 |
| A453S | 1 | 1 | 0 | 0 | 0 | 0 | 0 | 1 | 0 | 0 | -1 | 0 | 0 | 0 | 0 | 0 | 0 | 0 | 2 |
| N459D | 1 | 0 | 1 | 0 | 0 | 0 | 0 | 0 | 0 | 1 | -1 | 0 | 0 | 0 | 0 | 0 | 0 | 0 | 2 |
| R461Q | 0 | 0 | 1 | 1 | 0 | 0 | 0 | 0 | 0 | 1 | -1 | 0 | 0 | 0 | 0 | 0 | 0 | 0 | 2 |
| R461W | 0 | 0 | 1 | 0 | 0 | 0 | 0 | 0 | 1 | 1 | 0 | 0 | 1 | 0 | 0 | 0 | 0 | 0 | 4 |
| G469D | 0 | 0 | 1 | 0 | 0 | 0 | 0 | 1 | 0 | 1 | -1 | 0 | 1 | 0 | 0 | 0 | 0 | 1 | 4 |
| S471P | 0 | 0 | 1 | 0 | 1 | 0 | 0 | 1 | 0 | 0 | 0 | 0 | 0 | 0 | 0 | 0 | 0 | 0 | 3 |
| D473H | 1 | 0 | 0 | 0 | 0 | 0 | 0 | 0 | 1 | 1 | 0 | 0 | 0 | 0 | 0 | 0 | 0 | 0 | 3 |
| T475I | 0 | 0 | 0 | 1 | 0 | 0 | 0 | 1 | 0 | 0 | 0 | 0 | 1 | 0 | 0 | 0 | 1 | 0 | 4 |
| E478A | 0 | 1 | 0 | 0 | 0 | 0 | 0 | 1 | 0 | 1 | 0 | 0 | 0 | 0 | 0 | 0 | 1 | 0 | 4 |
| E495Q | 0 | 0 | 0 | 0 | 0 | 0 | 0 | 0 | 0 | 1 | -1 | 0 | 0 | 0 | 0 | 0 | 0 | 0 | 0 |
| R506T | 0 | 0 | 0 | 1 | 0 | 0 | 0 | 0 | 0 | 1 | 0 | 0 | 1 | 0 | 0 | 0 | 0 | 0 | 3 |
| P509A | 0 | 0 | 0 | 1 | 1 | 0 | 0 | 0 | 0 | 0 | 0 | 0 | 0 | 0 | 0 | 0 | 0 | 0 | 2 |
| R515S | 1 | 0 | 0 | 1 | 0 | 1 | 0 | 0 | 0 | 1 | 0 | 0 | 1 | 0 | 0 | 0 | 0 | 0 | 5 |
| P518Q | 0 | 0 | 0 | 1 | 1 | 1 | 0 | 1 | 0 | 0 | 0 | 0 | 0 | 0 | 0 | 0 | 0 | 0 | 4 |
| V524L | 1 | 1 | 0 | 0 | 0 | 1 | 0 | 0 | 0 | 0 | -1 | 0 | 0 | 0 | 0 | 0 | 0 | 0 | 2 |
| S527T | 0 | 0 | 0 | 1 | 0 | 1 | 0 | 0 | 0 | 0 | -1 | 1 | 0 | 0 | 0 | 0 | 0 | 0 | 2 |

| | | | | | | | | | | | | | | | | | | |
|---|---|---|---|---|---|---|---|---|---|---|---|---|---|---|---|---|---|---|
| T532R | 1 | 1 | 0 | 0 | 0 | 1 | 0 | 0 | 0 | 1 | 0 | 0 | 1 | 0 | 0 | 0 | 1 | 0 | 6 |
| N533Y | 0 | 1 | 0 | 0 | 0 | 1 | 0 | 0 | 1 | 0 | 0 | 0 | 1 | 0 | 0 | 0 | 1 | 0 | 5 |
| R536Q | 1 | 1 | 0 | 0 | 0 | 1 | 0 | 0 | 0 | 1 | -1 | 0 | 0 | 0 | 0 | 0 | 1 | 0 | 4 |
| R536W | 1 | 1 | 0 | 0 | 0 | 1 | 0 | 0 | 1 | 1 | 0 | 0 | 1 | 0 | 0 | 0 | 0 | 0 | 6 |
| D545H | 1 | 0 | 0 | 0 | 0 | 1 | 0 | 0 | 1 | 1 | 0 | 0 | 0 | 0 | 0 | 0 | 0 | 1 | 5 |
| S547C | 1 | 0 | 0 | 1 | 0 | 1 | 0 | 0 | 0 | 0 | 0 | 1 | 0 | 0 | 0 | 0 | 0 | 0 | 4 |
| L548P | 1 | 0 | 0 | 0 | 1 | 1 | 0 | 0 | 0 | 0 | 0 | 0 | 1 | 0 | 0 | 0 | 1 | 0 | 5 |
| L548V | 1 | 0 | 0 | 1 | 0 | 1 | 0 | 0 | 0 | 0 | -1 | 0 | 0 | 0 | 0 | 0 | 1 | 0 | 3 |
| S551I | 0 | 0 | 0 | 0 | 0 | 1 | 0 | 1 | 0 | 0 | 0 | 0 | 1 | 0 | 0 | 0 | 1 | 0 | 4 |
| M552V | 1 | 0 | 0 | 0 | 0 | 1 | 0 | 0 | 0 | 0 | -1 | 0 | 0 | 0 | 0 | 0 | 1 | 1 | 3 |
| L555R | 1 | 0 | 0 | 1 | 0 | 1 | 0 | 1 | 0 | 1 | 0 | 0 | 0 | 0 | 0 | 1 | 0 | 0 | 6 |
| T559N | 1 | 0 | 0 | 1 | 0 | 1 | 1 | 0 | 0 | 0 | -1 | 1 | 0 | 0 | 0 | 0 | 0 | 0 | 4 |
| M560V | 1 | 0 | 0 | 0 | 0 | 1 | 0 | 0 | 0 | 0 | -1 | 0 | 0 | 0 | 0 | 1 | 0 | 0 | 2 |
| L562P | 1 | 0 | 0 | 1 | 1 | 1 | 0 | 0 | 0 | 0 | 0 | 0 | 1 | 0 | 0 | 1 | 0 | 0 | 6 |
| S564I | 1 | 0 | 0 | 1 | 0 | 1 | 0 | 1 | 0 | 0 | 0 | 1 | 1 | 0 | 0 | 0 | 1 | 0 | 7 |
| K574N | 0 | 1 | 0 | 0 | 0 | 1 | 0 | 0 | 0 | 1 | -1 | 0 | 1 | 0 | 0 | 0 | 1 | 0 | 4 |
| H580D | 1 | 0 | 0 | 1 | 0 | 1 | 0 | 0 | 1 | 1 | 0 | 0 | 0 | 0 | 0 | 0 | 0 | 0 | 5 |
| H580Y | 1 | 0 | 0 | 1 | 0 | 1 | 0 | 0 | 0 | 1 | -1 | 0 | 0 | 0 | 0 | 0 | 0 | 0 | 3 |
| P581R | 1 | 0 | 0 | 1 | 1 | 1 | 0 | 1 | 0 | 1 | 0 | 0 | 1 | 0 | 0 | 0 | 0 | 0 | 7 |
| E597K | 0 | 1 | 0 | 0 | 0 | 1 | 0 | 0 | 0 | 1 | -1 | 0 | 0 | 0 | 0 | 0 | 0 | 0 | 2 |
| G607S | 1 | 0 | 0 | 0 | 0 | 1 | 0 | 1 | 0 | 0 | -1 | 0 | 0 | 0 | 0 | 0 | 0 | 0 | 2 |
| Y608D | 1 | 0 | 0 | 0 | 0 | 1 | 0 | 0 | 1 | 1 | 0 | 0 | 1 | 0 | 0 | 0 | 0 | 1 | 6 |
| V611G | 0 | 0 | 1 | 1 | 0 | 1 | 0 | 0 | 0 | 0 | 0 | 1 | 1 | 0 | 0 | 0 | 0 | 0 | 5 |
| P615L | 1 | 0 | 0 | 0 | 1 | 1 | 0 | 0 | 0 | 0 | 0 | 0 | 1 | 0 | 0 | 0 | 0 | 0 | 4 |
| N616D | 1 | 0 | 0 | 1 | 0 | 1 | 0 | 0 | 0 | 1 | -1 | 0 | 0 | 0 | 0 | 0 | 0 | 0 | 3 |
| N616Y | 1 | 0 | 0 | 0 | 0 | 1 | 0 | 0 | 1 | 0 | 0 | 0 | 1 | 0 | 0 | 0 | 0 | 0 | 4 |
| G618R | 1 | 1 | 0 | 0 | 0 | 1 | 1 | 1 | 0 | 1 | 0 | 0 | 1 | 0 | 0 | 0 | 0 | 0 | 7 |
| Q620R | 1 | 1 | 0 | 0 | 0 | 1 | 0 | 0 | 0 | 1 | -1 | 0 | 0 | 0 | 0 | 0 | 0 | 0 | 3 |
| Y623C | 0 | 1 | 0 | 0 | 0 | 1 | 0 | 0 | 1 | 0 | 0 | 0 | 1 | 0 | 0 | 0 | 0 | 0 | 4 |
| Y623H | 0 | 1 | 0 | 1 | 0 | 1 | 0 | 0 | 0 | 1 | -1 | 0 | 0 | 0 | 0 | 0 | 0 | 0 | 3 |
| A624D | 1 | 1 | 0 | 1 | 0 | 1 | 0 | 1 | 0 | 1 | 0 | 1 | 0 | 0 | 0 | 0 | 0 | 0 | 7 |
| I629T | 1 | 1 | 0 | 1 | 0 | 1 | 0 | 1 | 0 | 0 | 0 | 1 | 1 | 0 | 0 | 0 | 0 | 0 | 7 |
| R630P | 0 | 1 | 0 | 0 | 1 | 1 | 0 | 1 | 0 | 1 | 0 | 0 | 1 | 0 | 0 | 0 | 0 | 0 | 6 |
| C644F | 0 | 0 | 1 | 1 | 0 | 1 | 0 | 1 | 1 | 0 | 0 | 0 | 1 | 0 | 0 | 0 | 0 | 0 | 6 |
| P647L | 1 | 0 | 1 | 1 | 1 | 1 | 0 | 0 | 0 | 0 | 0 | 0 | 1 | 0 | 0 | 0 | 0 | 0 | 6 |
| P647R | 1 | 0 | 1 | 1 | 1 | 1 | 0 | 1 | 0 | 1 | 0 | 0 | 1 | 0 | 0 | 0 | 0 | 0 | 8 |
| H651R | 1 | 0 | 0 | 1 | 0 | 1 | 1 | 0 | 1 | 0 | -1 | 1 | 0 | 0 | 0 | 0 | 0 | 0 | 5 |
| G652E | 1 | 0 | 0 | 0 | 0 | 1 | 0 | 1 | 0 | 1 | 0 | 0 | 1 | 0 | 0 | 0 | 0 | 0 | 5 |
| S657N | 1 | 1 | 0 | 0 | 0 | 1 | 0 | 0 | 0 | 0 | -1 | 0 | 0 | 0 | 0 | 0 | 0 | 0 | 2 |
| S657R | 1 | 1 | 0 | 0 | 0 | 1 | 0 | 0 | 0 | 1 | 0 | 1 | 1 | 0 | 0 | 0 | 0 | 0 | 6 |
| A661P | 0 | 0 | 0 | 0 | 1 | 1 | 0 | 0 | 0 | 0 | 0 | 0 | 0 | 0 | 0 | 0 | 0 | 0 | 2 |
| M663K | 1 | 0 | 0 | 1 | 0 | 1 | 0 | 1 | 0 | 1 | 0 | 0 | 0 | 0 | 0 | 0 | 0 | 0 | 5 |
| A694P | 0 | 0 | 1 | 1 | 1 | 1 | 0 | 0 | 0 | 0 | 0 | 0 | 0 | 0 | 0 | 0 | 0 | 0 | 4 |
| T698I | 1 | 0 | 1 | 0 | 0 | 1 | 1 | 1 | 0 | 0 | 0 | 0 | 1 | 0 | 0 | 0 | 0 | 1 | 7 |
| P700A | 1 | 0 | 0 | 1 | 1 | 1 | 0 | 0 | 0 | 0 | 0 | 0 | 0 | 0 | 0 | 0 | 0 | 0 | 4 |
| S701F | 1 | 0 | 0 | 0 | 0 | 1 | 0 | 1 | 1 | 0 | 0 | 1 | 1 | 0 | 0 | 0 | 0 | 1 | 7 |
| G704V | 1 | 0 | 0 | 1 | 0 | 1 | 0 | 0 | 0 | 0 | 0 | 0 | 1 | 0 | 0 | 0 | 0 | 0 | 4 |
| N709S | 0 | 0 | 0 | 0 | 0 | 1 | 0 | 0 | 0 | 0 | -1 | 0 | 0 | 0 | 0 | 0 | 0 | 0 | 0 |
| V713M | 0 | 1 | 0 | 0 | 0 | 1 | 0 | 0 | 0 | 0 | -1 | 0 | 0 | 0 | 0 | 0 | 0 | 0 | 1 |
| C714R | 1 | 1 | 0 | 1 | 0 | 1 | 0 | 0 | 0 | 1 | 0 | 1 | 1 | 0 | 0 | 0 | 0 | 0 | 7 |
| L716H | 0 | 1 | 0 | 0 | 0 | 1 | 0 | 1 | 1 | 1 | 0 | 1 | 0 | 0 | 0 | 0 | 0 | 0 | 6 |
| I717N | 0 | 1 | 0 | 0 | 0 | 1 | 0 | 1 | 0 | 0 | 0 | 0 | 1 | 0 | 0 | 0 | 0 | 0 | 4 |
| I717V | 0 | 1 | 0 | 1 | 0 | 1 | 0 | 0 | 0 | 0 | -1 | 1 | 0 | 0 | 0 | 0 | 0 | 0 | 3 |
| G722R | 1 | 0 | 0 | 0 | 0 | 1 | 0 | 1 | 0 | 1 | 0 | 0 | 1 | 0 | 0 | 0 | 0 | 0 | 5 |
| G728E | 1 | 0 | 0 | 0 | 0 | 1 | 0 | 1 | 0 | 1 | 0 | 0 | 1 | 0 | 0 | 1 | 0 | 1 | 7 |
| G728R | 1 | 0 | 0 | 0 | 0 | 1 | 0 | 1 | 0 | 1 | 0 | 0 | 1 | 0 | 0 | 1 | 0 | 1 | 7 |
| N732K | 1 | 1 | 0 | 1 | 0 | 1 | 0 | 0 | 0 | 1 | -1 | 1 | 1 | 0 | 0 | 1 | 0 | 0 | 7 |
| A733D | 1 | 1 | 0 | 0 | 0 | 1 | 0 | 1 | 0 | 1 | 0 | 1 | 0 | 0 | 0 | 0 | 0 | 1 | 7 |
| A733V | 1 | 1 | 0 | 1 | 0 | 1 | 0 | 0 | 0 | 0 | -1 | 0 | 1 | 0 | 0 | 0 | 0 | 0 | 4 |
| R739H | 0 | 0 | 0 | 0 | 0 | 1 | 0 | 0 | 1 | 0 | -1 | 1 | 0 | 0 | 0 | 0 | 0 | 1 | 3 |
| D746E | 1 | 0 | 0 | 0 | 0 | 1 | 0 | 0 | 0 | 0 | -1 | 1 | 0 | 0 | 0 | 0 | 0 | 0 | 2 |
| D746H | 1 | 0 | 0 | 0 | 0 | 1 | 0 | 0 | 1 | 1 | 0 | 0 | 0 | 0 | 0 | 0 | 0 | 1 | 5 |
| H753P | 1 | 0 | 0 | 1 | 1 | 1 | 1 | 1 | 1 | 1 | 0 | 1 | 0 | 0 | 0 | 0 | 0 | 0 | 9 |
| P759R | 1 | 0 | 0 | 1 | 1 | 1 | 0 | 0 | 0 | 1 | 0 | 0 | 1 | 0 | 0 | 1 | 0 | 0 | 7 |
| H760P | 1 | 0 | 0 | 1 | 1 | 1 | 1 | 1 | 1 | 1 | 0 | 1 | 0 | 0 | 0 | 0 | 0 | 0 | 9 |
| H760Q | 1 | 0 | 0 | 1 | 0 | 1 | 1 | 0 | 1 | 1 | -1 | 0 | 0 | 0 | 0 | 0 | 0 | 0 | 5 |
| H760R | 1 | 0 | 0 | 1 | 0 | 1 | 1 | 0 | 1 | 0 | -1 | 1 | 0 | 0 | 0 | 0 | 0 | 0 | 5 |
| G761R | 1 | 0 | 0 | 0 | 0 | 1 | 0 | 1 | 0 | 1 | 0 | 0 | 1 | 0 | 0 | 0 | 0 | 1 | 6 |
| G762R | 1 | 0 | 0 | 1 | 0 | 1 | 0 | 1 | 0 | 1 | 0 | 0 | 1 | 0 | 0 | 0 | 0 | 0 | 6 |
| G763R | 1 | 0 | 0 | 1 | 0 | 1 | 0 | 1 | 0 | 1 | 0 | 1 | 1 | 0 | 0 | 0 | 0 | 0 | 7 |
| P765S | 1 | 0 | 0 | 1 | 1 | 1 | 0 | 1 | 0 | 0 | 0 | 0 | 0 | 0 | 0 | 1 | 0 | 0 | 6 |
| G766C | 1 | 0 | 0 | 0 | 0 | 1 | 0 | 1 | 0 | 0 | 0 | 0 | 0 | 0 | 0 | 1 | 0 | 1 | 5 |
| G766V | 1 | 0 | 0 | 0 | 0 | 1 | 0 | 0 | 0 | 0 | 0 | 1 | 1 | 0 | 0 | 1 | 0 | 1 | 6 |
| G768E | 1 | 0 | 0 | 0 | 0 | 1 | 0 | 1 | 0 | 1 | 0 | 0 | 1 | 0 | 0 | 0 | 0 | 1 | 6 |
| P769L | 1 | 0 | 0 | 1 | 1 | 1 | 0 | 0 | 0 | 0 | 0 | 0 | 1 | 0 | 0 | 0 | 0 | 0 | 5 |
| P769T | 1 | 0 | 0 | 0 | 1 | 1 | 0 | 1 | 0 | 0 | 0 | 0 | 0 | 0 | 0 | 0 | 0 | 0 | 4 |
| G771R | 0 | 0 | 1 | 1 | 0 | 1 | 0 | 1 | 0 | 1 | 0 | 0 | 1 | 0 | 0 | 0 | 0 | 0 | 6 |
| H775P | 1 | 1 | 0 | 0 | 1 | 1 | 0 | 1 | 1 | 1 | 0 | 1 | 0 | 0 | 0 | 0 | 0 | 0 | 8 |
| H775R | 1 | 1 | 0 | 0 | 0 | 1 | 0 | 0 | 1 | 0 | -1 | 1 | 0 | 0 | 0 | 0 | 0 | 0 | 4 |
| H775Y | 1 | 1 | 0 | 0 | 0 | 1 | 0 | 0 | 0 | 1 | -1 | 0 | 0 | 0 | 0 | 0 | 0 | 0 | 3 |
| A777P | 0 | 1 | 0 | 0 | 1 | 1 | 0 | 0 | 0 | 0 | 0 | 0 | 0 | 0 | 0 | 0 | 0 | 0 | 3 |
| P778A | 0 | 1 | 0 | 1 | 1 | 1 | 0 | 0 | 0 | 0 | 0 | 0 | 0 | 0 | 0 | 0 | 0 | 0 | 4 |
| R790W | 0 | 1 | 0 | 0 | 0 | 1 | 0 | 0 | 1 | 1 | 0 | 0 | 1 | 0 | 0 | 0 | 0 | 0 | 5 |
| E792D | 0 | 1 | 0 | 0 | 0 | 1 | 0 | 0 | 0 | 0 | -1 | 0 | 0 | 0 | 0 | 0 | 0 | 0 | 1 |
| C795S | 0 | 0 | 0 | 1 | 0 | 1 | 0 | 0 | 0 | 0 | 0 | 0 | 0 | 0 | 0 | 0 | 0 | 0 | 2 |
| A802E | 0 | 0 | 0 | 0 | 0 | 1 | 0 | 1 | 0 | 1 | 0 | 0 | 0 | 0 | 1 | 0 | 0 | 0 | 4 |
| A802V | 0 | 0 | 0 | 0 | 0 | 1 | 0 | 0 | 0 | 0 | -1 | 0 | 1 | 0 | 1 | 0 | 0 | 0 | 2 |
| A803V | 1 | 0 | 0 | 0 | 0 | 1 | 0 | 0 | 0 | 0 | -1 | 0 | 1 | 0 | 0 | 0 | 0 | 0 | 2 |
| G806R | 1 | 0 | 0 | 1 | 0 | 1 | 0 | 1 | 0 | 1 | 0 | 1 | 1 | 0 | 0 | 0 | 0 | 0 | 7 |
| S808I | 1 | 1 | 0 | 1 | 0 | 1 | 0 | 1 | 0 | 0 | 0 | 1 | 1 | 0 | 0 | 0 | 0 | 0 | 7 |
| S808R | 1 | 1 | 0 | 0 | 0 | 1 | 0 | 0 | 0 | 1 | 0 | 1 | 1 | 0 | 0 | 0 | 0 | 1 | 7 |
| A816T | 0 | 1 | 0 | 1 | 0 | 1 | 0 | 1 | 0 | 0 | -1 | 0 | 0 | 0 | 0 | 0 | 0 | 0 | 3 |
| Y817C | 1 | 1 | 0 | 0 | 0 | 1 | 0 | 0 | 1 | 0 | 0 | 0 | 1 | 0 | 0 | 1 | 0 | 0 | 6 |
| K819E | 0 | 1 | 0 | 0 | 0 | 1 | 0 | 0 | 0 | 1 | -1 | 0 | 0 | 0 | 0 | 0 | 0 | 0 | 2 |
| M820V | 1 | 1 | 0 | 0 | 0 | 1 | 0 | 0 | 0 | 0 | -1 | 0 | 0 | 0 | 0 | 0 | 0 | 0 | 2 |
| T830K | 1 | 0 | 0 | 1 | 0 | 1 | 0 | 0 | 0 | 1 | 0 | 0 | 1 | 0 | 0 | 0 | 0 | 0 | 5 |
| T830M | 1 | 0 | 0 | 0 | 0 | 1 | 0 | 1 | 0 | 0 | 0 | 0 | 0 | 0 | 0 | 0 | 0 | 1 | 4 |
| A833V | 1 | 1 | 0 | 0 | 0 | 1 | 0 | 0 | 0 | 0 | -1 | 0 | 1 | 0 | 0 | 0 | 0 | 0 | 3 |
| N838D | 1 | 1 | 0 | 0 | 0 | 1 | 0 | 0 | 0 | 1 | -1 | 0 | 0 | 0 | 0 | 0 | 0 | 0 | 3 |
| Y839C | 1 | 1 | 0 | 0 | 0 | 1 | 0 | 0 | 1 | 0 | 0 | 0 | 1 | 0 | 0 | 0 | 0 | 0 | 5 |
| M840K | 0 | 1 | 0 | 0 | 0 | 1 | 0 | 1 | 0 | 1 | 0 | 0 | 0 | 0 | 0 | 0 | 0 | 0 | 4 |
| M840V | 0 | 1 | 0 | 1 | 0 | 1 | 0 | 0 | 0 | 0 | -1 | 0 | 0 | 0 | 0 | 0 | 0 | 0 | 2 |
| A841P | 1 | 1 | 0 | 1 | 1 | 1 | 0 | 0 | 0 | 0 | 0 | 0 | 0 | 0 | 0 | 0 | 0 | 0 | 5 |
| T846I | 0 | 0 | 0 | 1 | 0 | 1 | 0 | 1 | 0 | 0 | 0 | 0 | 1 | 0 | 0 | 0 | 0 | 0 | 4 |
| R849G | 0 | 0 | 1 | 1 | 0 | 1 | 0 | 1 | 0 | 1 | 0 | 0 | 1 | 0 | 0 | 0 | 0 | 0 | 6 |
| A855V | 0 | 1 | 0 | 0 | 0 | 1 | 0 | 0 | 0 | 0 | -1 | 0 | 1 | 0 | 0 | 0 | 0 | 0 | 2 |
| G860R | 1 | 0 | 0 | 1 | 0 | 1 | 0 | 1 | 0 | 1 | 0 | 1 | 1 | 0 | 0 | 0 | 0 | 0 | 7 |
| E862K | 1 | 0 | 0 | 0 | 0 | 1 | 0 | 0 | 0 | 1 | -1 | 0 | 0 | 0 | 0 | 0 | 0 | 0 | 2 |
| D866H | 1 | 0 | 1 | 0 | 0 | 1 | 0 | 0 | 1 | 1 | 0 | 0 | 0 | 0 | 0 | 0 | 0 | 0 | 5 |
| D880E | 1 | 1 | 0 | 0 | 0 | 1 | 0 | 0 | 0 | 0 | -1 | 1 | 0 | 0 | 0 | 0 | 0 | 0 | 3 |
| D880V | 1 | 1 | 0 | 0 | 0 | 1 | 0 | 1 | 0 | 1 | 0 | 1 | 0 | 0 | 0 | 0 | 0 | 1 | 7 |

| | | | | | | | | | | | | | | | | | | | |
|---|---|---|---|---|---|---|---|---|---|---|---|---|---|---|---|---|---|---|---|
| R884G | 1 | 1 | 0 | 0 | 0 | 1 | 0 | 1 | 0 | 1 | 0 | 0 | 1 | 0 | 0 | 0 | 0 | 0 | 6 |
| L885P | 1 | 1 | 0 | 1 | 1 | 1 | 0 | 0 | 0 | 0 | 0 | 0 | 1 | 0 | 0 | 0 | 0 | 0 | 6 |
| G889V | 1 | 0 | 0 | 1 | 0 | 1 | 0 | 0 | 0 | 0 | 0 | 0 | 1 | 0 | 0 | 0 | 0 | 0 | 4 |
| P893L | 1 | 0 | 0 | 1 | 1 | 1 | 0 | 0 | 0 | 0 | 0 | 0 | 1 | 0 | 0 | 0 | 0 | 0 | 5 |
| T894A | 1 | 0 | 1 | 0 | 0 | 1 | 1 | 1 | 0 | 0 | -1 | 0 | 0 | 0 | 0 | 0 | 0 | 0 | 4 |
| M895T | 1 | 0 | 1 | 0 | 0 | 1 | 0 | 1 | 0 | 0 | 0 | 0 | 0 | 0 | 0 | 0 | 0 | 0 | 4 |
| W897C | 0 | 0 | 0 | 0 | 0 | 1 | 1 | 0 | 1 | 0 | 0 | 1 | 1 | 0 | 0 | 0 | 0 | 0 | 5 |
| V905G | 0 | 0 | 1 | 1 | 0 | 1 | 0 | 0 | 0 | 0 | 0 | 0 | 1 | 0 | 0 | 0 | 0 | 0 | 4 |
| P907L | 1 | 0 | 0 | 0 | 1 | 1 | 0 | 0 | 0 | 0 | 0 | 0 | 1 | 0 | 0 | 0 | 0 | 0 | 4 |
| T908P | 1 | 0 | 0 | 0 | 1 | 1 | 0 | 1 | 0 | 0 | 0 | 0 | 0 | 0 | 0 | 1 | 0 | 0 | 5 |
| S910L | 1 | 0 | 0 | 0 | 0 | 1 | 0 | 1 | 0 | 0 | 0 | 0 | 1 | 0 | 0 | 0 | 1 | 1 | 6 |
| D917E | 1 | 1 | 0 | 0 | 0 | 1 | 0 | 0 | 0 | 0 | -1 | 0 | 0 | 0 | 0 | 0 | 0 | 0 | 2 |
| F919I | 1 | 1 | 0 | 0 | 0 | 1 | 0 | 0 | 1 | 0 | -1 | 0 | 0 | 0 | 0 | 0 | 0 | 0 | 3 |
| I933F | 0 | 1 | 0 | 1 | 0 | 1 | 0 | 0 | 1 | 0 | -1 | 1 | 0 | 0 | 0 | 0 | 0 | 0 | 4 |
| I933N | 0 | 1 | 0 | 0 | 0 | 1 | 0 | 1 | 0 | 0 | 0 | 1 | 1 | 0 | 0 | 0 | 0 | 0 | 5 |
| I933T | 0 | 1 | 0 | 1 | 0 | 1 | 0 | 1 | 0 | 0 | 0 | 0 | 1 | 0 | 0 | 0 | 0 | 0 | 5 |
| M947I | 0 | 1 | 0 | 0 | 0 | 1 | 0 | 0 | 0 | 0 | -1 | 0 | 0 | 0 | 0 | 0 | 0 | 0 | 1 |
| P949L | 1 | 0 | 0 | 1 | 1 | 1 | 0 | 0 | 0 | 0 | 0 | 0 | 1 | 0 | 0 | 0 | 0 | 0 | 5 |
| H950R | 1 | 0 | 0 | 1 | 0 | 1 | 0 | 0 | 1 | 0 | -1 | 1 | 0 | 0 | 0 | 0 | 0 | 0 | 4 |
| S951Y | 1 | 0 | 0 | 0 | 0 | 1 | 0 | 0 | 1 | 0 | 0 | 1 | 1 | 0 | 0 | 0 | 0 | 0 | 5 |
| S957P | 0 | 0 | 0 | 0 | 1 | 1 | 0 | 1 | 0 | 0 | 0 | 0 | 0 | 0 | 0 | 0 | 1 | 0 | 4 |
| W960C | 1 | 0 | 0 | 0 | 0 | 1 | 0 | 0 | 1 | 0 | 0 | 0 | 1 | 0 | 0 | 0 | 0 | 0 | 4 |
| R962P | 0 | 0 | 0 | 0 | 1 | 1 | 0 | 1 | 0 | 1 | 0 | 0 | 1 | 0 | 0 | 0 | 0 | 0 | 5 |
| R962Q | 0 | 0 | 0 | 1 | 0 | 1 | 0 | 0 | 0 | 1 | -1 | 0 | 0 | 0 | 0 | 0 | 0 | 0 | 2 |
| R966G | 1 | 1 | 0 | 1 | 0 | 1 | 0 | 1 | 0 | 1 | 0 | 0 | 1 | 0 | 0 | 0 | 1 | 0 | 8 |
| E967D | 0 | 1 | 0 | 0 | 0 | 1 | 0 | 0 | 0 | 0 | -1 | 0 | 0 | 0 | 0 | 0 | 1 | 0 | 2 |
| E979A | 0 | 0 | 0 | 0 | 0 | 1 | 0 | 1 | 0 | 1 | 0 | 0 | 0 | 0 | 0 | 0 | 1 | 0 | 4 |
| N980D | 0 | 0 | 0 | 0 | 0 | 1 | 0 | 0 | 0 | 1 | -1 | 0 | 0 | 0 | 0 | 0 | 1 | 0 | 2 |
| W983R | 1 | 0 | 0 | 0 | 0 | 1 | 0 | 0 | 1 | 1 | 0 | 0 | 1 | 0 | 0 | 0 | 1 | 1 | 7 |
| R988Q | 1 | 0 | 0 | 0 | 0 | 1 | 0 | 0 | 0 | 1 | -1 | 0 | 0 | 0 | 0 | 0 | 0 | 0 | 2 |
| R988W | 1 | 0 | 0 | 0 | 0 | 1 | 0 | 0 | 1 | 1 | 0 | 0 | 1 | 0 | 0 | 0 | 0 | 0 | 5 |
| G994R | 1 | 1 | 0 | 1 | 0 | 1 | 0 | 1 | 0 | 1 | 0 | 0 | 1 | 0 | 0 | 0 | 0 | 0 | 7 |
| D995N | 1 | 1 | 0 | 0 | 0 | 1 | 0 | 0 | 0 | 1 | -1 | 0 | 0 | 0 | 0 | 0 | 0 | 0 | 3 |
| Q996H | 1 | 1 | 0 | 0 | 0 | 1 | 0 | 0 | 1 | 1 | -1 | 0 | 0 | 0 | 0 | 0 | 0 | 0 | 4 |
| H997Q | 1 | 0 | 0 | 0 | 0 | 1 | 0 | 0 | 1 | 1 | -1 | 0 | 0 | 0 | 0 | 0 | 0 | 0 | 3 |
| C1002W | 1 | 0 | 0 | 0 | 0 | 1 | 0 | 0 | 1 | 0 | 0 | 0 | 1 | 0 | 0 | 0 | 0 | 0 | 4 |
| **Non-Pathogenic Mutations** | | | | | | | | | | | | | | | | | | | |
| A64S | 0 | 1 | 0 | 1 | 0 | 0 | 0 | 1 | 0 | 0 | -1 | 0 | 0 | 0 | 0 | 0 | 0 | 0 | 2 |
| A64T | 0 | 1 | 0 | 1 | 0 | 0 | 0 | 1 | 0 | 0 | -1 | 0 | 0 | 0 | 0 | 0 | 0 | 0 | 2 |
| R66K | 0 | 1 | 0 | 1 | 0 | 0 | 0 | 0 | 0 | 0 | -1 | 0 | 0 | 0 | 0 | 0 | 1 | 0 | 2 |
| M107V | 0 | 0 | 0 | 1 | 0 | 0 | 0 | 0 | 0 | 0 | -1 | 0 | 0 | 0 | 0 | 0 | 0 | 0 | 0 |
| G137S | 1 | 0 | 0 | 0 | 0 | 0 | 0 | 1 | 0 | 0 | -1 | 0 | 0 | 0 | 1 | 0 | 0 | 0 | 2 |
| N193S | 1 | 0 | 0 | 0 | 0 | 0 | 0 | 0 | 0 | 0 | -1 | 0 | 0 | 0 | 0 | 0 | 0 | 0 | 0 |
| L207V | 1 | 1 | 0 | 0 | 0 | 0 | 0 | 0 | 0 | 0 | -1 | 0 | 0 | 0 | 0 | 0 | 0 | 0 | 1 |
| R224H | 0 | 0 | 0 | 1 | 0 | 0 | 0 | 0 | 1 | 0 | -1 | 1 | 0 | 0 | 0 | 0 | 0 | 0 | 2 |
| V233A | 0 | 1 | 0 | 1 | 0 | 0 | 0 | 0 | 0 | 0 | -1 | 0 | 1 | 0 | 0 | 0 | 0 | 0 | 2 |
| R236Q | 1 | 1 | 0 | 0 | 0 | 0 | 0 | 0 | 0 | 1 | -1 | 0 | 0 | 0 | 0 | 0 | 0 | 0 | 2 |
| E278K | 0 | 1 | 0 | 0 | 0 | 0 | 0 | 0 | 0 | 1 | -1 | 0 | 0 | 0 | 0 | 0 | 0 | 0 | 1 |
| C291G | 0 | 0 | 1 | 1 | 0 | 0 | 0 | 0 | 0 | 0 | 0 | 0 | 0 | 0 | 0 | 0 | 0 | 0 | 2 |
| I301M | 0 | 0 | 0 | 1 | 0 | 0 | 0 | 0 | 0 | 0 | -1 | 0 | 0 | 0 | 0 | 0 | 0 | 0 | 0 |
| R410K | 0 | 1 | 0 | 0 | 0 | 0 | 0 | 0 | 0 | 0 | -1 | 0 | 0 | 0 | 0 | 0 | 0 | 0 | 0 |
| N413Y | 0 | 1 | 0 | 0 | 0 | 0 | 0 | 0 | 1 | 0 | 0 | 0 | 1 | 0 | 0 | 0 | 1 | 0 | 4 |
| A414T | 0 | 1 | 0 | 0 | 0 | 0 | 0 | 1 | 0 | 0 | -1 | 0 | 0 | 0 | 0 | 0 | 1 | 0 | 2 |
| L462V | 0 | 0 | 1 | 0 | 0 | 0 | 0 | 0 | 0 | 0 | -1 | 0 | 0 | 0 | 0 | 0 | 0 | 0 | 0 |
| E503A | 0 | 0 | 0 | 0 | 0 | 0 | 0 | 1 | 0 | 1 | 0 | 0 | 0 | 0 | 0 | 0 | 0 | 0 | 2 |
| P509A | 0 | 0 | 0 | 1 | 1 | 0 | 0 | 0 | 0 | 0 | 0 | 0 | 0 | 0 | 0 | 0 | 0 | 0 | 2 |
| N533S | 0 | 1 | 0 | 1 | 0 | 1 | 0 | 0 | 0 | 0 | -1 | 0 | 0 | 0 | 0 | 0 | 1 | 0 | 3 |
| A569T | 0 | 1 | 0 | 0 | 0 | 1 | 0 | 1 | 0 | 0 | -1 | 0 | 0 | 0 | 0 | 0 | 1 | 0 | 3 |
| R596Q | 0 | 1 | 0 | 0 | 0 | 1 | 0 | 0 | 0 | 1 | -1 | 0 | 0 | 0 | 0 | 0 | 0 | 0 | 2 |
| E669K | 0 | 0 | 0 | 0 | 0 | 1 | 0 | 0 | 0 | 1 | -1 | 0 | 0 | 0 | 0 | 0 | 0 | 0 | 1 |
| V705M | 1 | 0 | 0 | 0 | 0 | 1 | 0 | 0 | 0 | 0 | -1 | 0 | 0 | 0 | 0 | 0 | 0 | 0 | 1 |
| V735L | 1 | 0 | 0 | 1 | 0 | 1 | 0 | 0 | 0 | 0 | -1 | 0 | 0 | 0 | 0 | 0 | 0 | 0 | 2 |
| V747I | 1 | 0 | 1 | 1 | 0 | 1 | 0 | 0 | 0 | 0 | -1 | 1 | 0 | 0 | 0 | 0 | 0 | 0 | 4 |
| A794T | 0 | 0 | 0 | 0 | 0 | 1 | 0 | 1 | 0 | 0 | -1 | 0 | 0 | 0 | 0 | 0 | 0 | 0 | 1 |
| T799S | 0 | 0 | 0 | 1 | 0 | 1 | 0 | 0 | 0 | 0 | -1 | 1 | 0 | 0 | 0 | 0 | 0 | 0 | 2 |
| V800I | 0 | 0 | 0 | 1 | 0 | 1 | 0 | 0 | 0 | 0 | -1 | 1 | 0 | 0 | 0 | 0 | 0 | 0 | 2 |
| S814F | 1 | 1 | 0 | 0 | 0 | 1 | 0 | 1 | 1 | 0 | 0 | 1 | 1 | 0 | 0 | 0 | 0 | 1 | 8 |
| R868T | 1 | 1 | 0 | 0 | 0 | 1 | 0 | 0 | 0 | 1 | 0 | 0 | 1 | 0 | 0 | 0 | 0 | 0 | 5 |
| M895V | 1 | 0 | 1 | 0 | 0 | 1 | 0 | 0 | 0 | 0 | -1 | 0 | 0 | 0 | 0 | 0 | 0 | 1 | 3 |
| E979D | 0 | 0 | 0 | 0 | 0 | 1 | 0 | 0 | 0 | 0 | -1 | 0 | 0 | 0 | 0 | 0 | 1 | 0 | 1 |
| T1001N | 0 | 0 | 0 | 1 | 0 | 1 | 0 | 0 | 0 | 0 | -1 | 1 | 0 | 0 | 0 | 0 | 0 | 0 | 2 |
