## Supplemental Table 6 for "Large scale analyses of genotype-phenotype relationships of glycine decarboxylase mutations and neurological disease severity"

| WT | Position | Mutant | COS | Conservation of Amino Acid | Helix Mutation | Sheet Mutation | Destabilizing | Δ Proline | C-terminal mutation | Active Site Mutation | Δ Polarity | Δ Aromaticity | Δ Charge | Conservation of Substition | Δ Codon | Δ Size | H-interface Mutation | N-PLP Mutation | Active Region | Dimerazation Interface | Stabilizing Mutation | SUM |
|---|---|---|---|---|---|---|---|---|---|---|---|---|---|---|---|---|---|---|---|---|---|---|
| **Pathogenic Mutations** | | | | | | | | | | | | | | | | | | | | | | |
| Y | 161 | C | 12.0 | 4.1 | 0.0 | 0.0 | 0.0 | 0.0 | 0.0 | 4.2 | 0.0 | 2.9 | 0.0 | 0.0 | 0.0 | 0.7 | 0.0 | 0.0 | 0.0 | 0.0 | 0.0 | 11.9 |
| Y | 164 | H | 3.3 | 4.1 | 0.0 | 0.0 | 0.0 | 0.0 | 0.0 | 4.2 | 0.0 | 0.0 | 1.1 | -5.0 | 0.0 | 0.0 | 0.0 | 0.0 | 0.0 | 0.0 | 0.0 | 4.4 |
| D | 198 | V | 9.0 | 4.1 | 0.0 | 0.0 | 0.0 | 0.0 | 0.0 | 0.0 | 0.0 | 0.0 | 1.1 | 0.0 | 3.1 | 0.0 | 0.0 | 1.8 | 0.0 | 0.0 | 0.0 | 10.1 |
| A | 202 | V | 2.0 | 4.1 | 3.1 | 0.0 | 0.0 | 0.0 | 0.0 | 0.0 | 0.0 | 0.0 | 0.0 | -5.0 | 0.0 | 0.7 | 0.0 | 0.0 | 0.0 | 0.0 | 0.0 | 2.9 |
| T | 269 | M | 2.0 | 0.0 | 0.0 | 0.0 | 0.0 | 0.0 | 0.0 | 0.0 | 0.0 | 0.0 | 0.0 | 0.0 | 0.0 | 0.0 | 0.0 | 0.0 | 0.0 | 0.0 | 0.0 | 0.0 |
| C | 291 | Y | 2.0 | 0.0 | 0.0 | 0.0 | 0.0 | 0.0 | 0.0 | 0.0 | 0.0 | 2.9 | 0.0 | 0.0 | 0.0 | 0.7 | 0.0 | 0.0 | 0.0 | 0.0 | 0.0 | 3.6 |
| R | 362 | C | 2.0 | 4.1 | 0.0 | 0.0 | 0.0 | 0.0 | 0.0 | 0.0 | 0.0 | 0.0 | 1.1 | 0.0 | 3.1 | 0.7 | 0.0 | 0.0 | 0.0 | 0.0 | 0.0 | 9.0 |
| Q | 366 | R | 3.0 | 4.1 | 0.0 | 0.0 | 0.0 | 0.0 | 0.0 | 0.0 | 0.0 | 0.0 | 1.1 | -5.0 | 0.0 | 0.0 | 3.6 | 0.0 | 0.0 | 0.0 | 0.0 | 3.8 |
| I | 372 | F | 9.0 | 4.1 | 0.0 | 0.0 | 0.0 | 0.0 | 0.0 | 0.0 | 0.0 | 2.9 | 0.0 | -5.0 | 3.1 | 0.0 | 3.6 | 0.0 | 0.0 | 0.0 | 0.0 | 8.7 |
| A | 389 | V | 4.5 | 4.1 | 3.1 | 0.0 | 0.0 | 0.0 | 0.0 | 0.0 | 0.0 | 0.0 | 0.0 | -5.0 | 0.0 | 0.7 | 0.0 | 0.0 | 0.0 | 0.0 | 0.0 | 2.9 |
| R | 515 | S | 9.0 | 4.1 | 0.0 | 0.0 | 0.0 | 0.0 | 3.3 | 0.0 | 0.0 | 0.0 | 1.1 | 0.0 | 0.0 | 0.7 | 0.0 | 0.0 | 0.0 | 0.0 | 0.0 | 9.2 |
| P | 581 | R | 7.0 | 4.1 | 0.0 | 0.0 | 0.0 | 0.0 | 3.3 | 0.0 | 0.0 | 0.0 | 1.1 | 0.0 | 0.0 | 0.7 | 0.0 | 0.0 | 0.0 | 0.0 | 0.0 | 9.2 |
| Y | 623 | H | 3.0 | 0.0 | 3.1 | 0.0 | 0.0 | 0.0 | 3.3 | 0.0 | 0.0 | 0.0 | 1.1 | -5.0 | 0.0 | 0.0 | 0.0 | 0.0 | 0.0 | 0.0 | 0.0 | 2.5 |
| R | 739 | H | 4.0 | 0.0 | 0.0 | 0.0 | 0.0 | 0.0 | 3.3 | 0.0 | 0.0 | 2.9 | 0.0 | -5.0 | 3.1 | 0.0 | 0.0 | 0.0 | 0.0 | 0.0 | 0.0 | 4.3 |
| A | 802 | V | 3.5 | 0.0 | 0.0 | 0.0 | 0.0 | 0.0 | 3.3 | 0.0 | 0.0 | 0.0 | 0.0 | -5.0 | 0.0 | 0.7 | 0.0 | 1.8 | 0.0 | 0.0 | 0.0 | 0.8 |
| P | 949 | L | 9.0 | 4.1 | 0.0 | 0.0 | 0.0 | 0.0 | 3.3 | 0.0 | 0.0 | 0.0 | 0.0 | 0.0 | 0.0 | 0.7 | 0.0 | 0.0 | 0.0 | 0.0 | 0.0 | 8.1 |
| H | 950 | R | 9.0 | 4.1 | 0.0 | 0.0 | 0.0 | 0.0 | 3.3 | 0.0 | 0.0 | 2.9 | 0.0 | -5.0 | 3.1 | 0.0 | 0.0 | 0.0 | 0.0 | 0.0 | 0.0 | 8.4 |
| R | 988 | Q | 7.0 | 4.1 | 0.0 | 0.0 | 0.0 | 0.0 | 3.3 | 0.0 | 0.0 | 0.0 | 1.1 | -5.0 | 0.0 | 0.0 | 0.0 | 0.0 | 0.0 | 0.0 | 0.0 | 3.5 |
| **Non Pathogenic Mutations** | | | | | | | | | | | | | | | | | | | | | | |
| M | 107 | V | 0.0 | 0.0 | 0.0 | 0.0 | 0.0 | 0.0 | 0.0 | 0.0 | 0.0 | 0.0 | 0.0 | -5.0 | 0.0 | 0.0 | 0.0 | 0.0 | 0.0 | 0.0 | 0.0 | 0.0 |
| R | 224 | H | 0.0 | 0.0 | 0.0 | 0.0 | 0.0 | 0.0 | 0.0 | 0.0 | 0.0 | 2.9 | 0.0 | -5.0 | 3.1 | 0.0 | 0.0 | 0.0 | 0.0 | 0.0 | 0.0 | 1.0 |
| C | 291 | G | 0.0 | 0.0 | 0.0 | 0.0 | 0.0 | 0.0 | 0.0 | 0.0 | 0.0 | 0.0 | 0.0 | 0.0 | 0.0 | 0.0 | 0.0 | 0.0 | 0.0 | 0.0 | 0.0 | 0.0 |
| R | 410 | K | 0.0 | 0.0 | 3.1 | 0.0 | 0.0 | 0.0 | 0.0 | 0.0 | 0.0 | 0.0 | 0.0 | -5.0 | 0.0 | 0.0 | 0.0 | 0.0 | 0.0 | 0.0 | 0.0 | 0.0 |
| L | 462 | V | 0.0 | 0.0 | 0.0 | 0.0 | 0.0 | 0.0 | 0.0 | 0.0 | 0.0 | 0.0 | 0.0 | -5.0 | 0.0 | 0.0 | 0.0 | 0.0 | 0.0 | 0.0 | 0.0 | 0.0 |
| E | 503 | A | 0.0 | 0.0 | 0.0 | 0.0 | 0.0 | 0.0 | 0.0 | 0.0 | 0.0 | 0.0 | 1.1 | 0.0 | 0.0 | 0.0 | 0.0 | 0.0 | 0.0 | 0.0 | 0.0 | 1.1 |
| A | 569 | T | 0.0 | 0.0 | 3.1 | 0.0 | 0.0 | 0.0 | 3.3 | 0.0 | 0.0 | 0.0 | 0.0 | -5.0 | 0.0 | 0.0 | 0.0 | 0.0 | 0.0 | 0.0 | 0.0 | 1.4 |
| V | 705 | M | 0.0 | 4.1 | 0.0 | 0.0 | 0.0 | 0.0 | 3.3 | 0.0 | 0.0 | 0.0 | 0.0 | -5.0 | 0.0 | 0.0 | 0.0 | 0.0 | 0.0 | 0.0 | 0.0 | 2.4 |
| V | 735 | L | 0.0 | 4.1 | 0.0 | 0.0 | 0.0 | 0.0 | 3.3 | 0.0 | 0.0 | 0.0 | 0.0 | -5.0 | 0.0 | 0.0 | 0.0 | 0.0 | 0.0 | 0.0 | 0.0 | 2.4 |
| A | 794 | T | 0.0 | 0.0 | 0.0 | 0.0 | 0.0 | 0.0 | 3.3 | 0.0 | 0.0 | 0.0 | 0.0 | -5.0 | 0.0 | 0.0 | 0.0 | 0.0 | 0.0 | 0.0 | 0.0 | 0.0 |
