## Supplemental Table 7 for "Large scale analyses of genotype-phenotype relationships of glycine decarboxylase mutations and neurological disease severity"

| Allele 1 | Allele 2 | COS | Allele 1 Score | Allele 2 Score | Patient Score |
|---|---|---|---|---|---|
| L885P | W897C | 12 | 14.7 | 11.3 | 13.0 |
| Y839C | Intron | 9 | 10.6 | 8.7 | 9.7 |
| G761R | N/FS | 9 | 8.6 | 15.5 | 12.1 |
| S132L | N/FS | 9 | 8.9 | 15.5 | 12.2 |
| S132L | N/FS | 8 | 8.9 | 15.5 | 12.2 |
| A377V | N/FS | 8 | 4.3 | 15.5 | 9.9 |
| T894A | Del | 8 | 13.5 | 14.0 | 13.8 |
| R461Q | Del | 8 | 8.7 | 14.0 | 11.4 |
| H371D | Del | 7 | 7.4 | 14.0 | 10.7 |
| R515S | Intron | 7 | 9.9 | 8.7 | 9.3 |
| Q620R | Del | 6 | 9.3 | 14.0 | 11.7 |
| P509A | E597K | 6 | 6.3 | 6.9 | 6.6 |
| D295Y | R536Q | 6 | 9.3 | 9.3 | 9.3 |
| P907L | Del | 6 | 10.8 | 14.0 | 12.4 |
| R515S | G618R | 6 | 9.9 | 16.2 | 13.1 |
| R515S | Intron | 6 | 9.9 | 8.7 | 9.3 |
| L885P | N/FS | 6 | 14.7 | 15.5 | 15.1 |
| G771R | Intron | 6 | 12.1 | 8.7 | 10.4 |
| F334L | Del | 6 | 7.5 | 14.0 | 10.8 |
| A389V | R515S | 6 | 6.9 | 9.9 | 8.4 |
| N150T | R790W | 6 | 7.0 | 11.0 | 9.0 |
| C1002W | N/FS | 5 | 8.0 | 15.5 | 11.8 |
| L82W | N/FS | 4 | 6.7 | 15.5 | 11.1 |
| C1002W | N/FS | 4 | 8.0 | 15.5 | 11.8 |
| A733V | Intron | 4 | 9.7 | 8.7 | 9.2 |
| A389V | R515S | 4 | 6.9 | 9.9 | 8.4 |
| A283P | R461Q | 4 | 7.6 | 8.7 | 8.2 |
| P267A | K376E | 3 | 10.1 | 5.2 | 7.7 |
| G771R | M552V | 3 | 12.1 | 3.9 | 8.0 |
| R739H | Del | 3 | 4.4 | 14.0 | 9.2 |
| L548V | Del | 3 | 5.2 | 14.0 | 9.6 |
| A802V | Intron | 3 | 4.8 | 8.7 | 6.8 |
| A802V | R515S | 3 | 4.8 | 9.9 | 7.4 |
| A202V | Intron | 2 | 6.9 | 8.7 | 7.8 |
| A202V | Intron | 2 | 6.9 | 8.7 | 7.8 |
| A802V | Intron | 2 | 4.8 | 8.7 | 6.8 |
| A802V | Intron | 2 | 4.8 | 8.7 | 6.8 |
| A389V | Intron | 2 | 6.9 | 8.7 | 7.8 |
| A389V | Intron | 2 | 6.9 | 8.7 | 7.8 |
| R373W | Intron | 2 | 9.3 | 8.7 | 9.0 |
| R373W | MLS | 2 | 9.3 | 4.0 | 6.7 |
| I381T | R461Q | 2 | 6.3 | 8.7 | 7.5 |
| A283P | R461Q | 2 | 7.6 | 8.7 | 8.2 |
| R461Q | Intron | 2 | 8.7 | 8.7 | 8.7 |
| Y161C | R347S | 2 | 11.5 | 7.1 | 9.3 |
| G156R | G728E | 2 | 8.6 | 8.6 | 8.6 |
| R630P | L548V | 2 | 13.8 | 5.2 | 9.5 |
| A802E | Intron | 2 | 5.7 | 8.7 | 7.2 |
| V905G | G728E | 2 | 9.3 | 8.6 | 9.0 |
| G652E | R373Q | 2 | 8.6 | 5.2 | 6.9 |
| E503A | V233A | 0 | 2.8 | 5.8 | 4.3 |
| E503A | T799S | 0 | 2.8 | 3.5 | 3.2 |
| E503A | R236Q | 0 | 2.8 | 7.8 | 5.3 |
| E503A | V705M | 0 | 2.8 | 3.9 | 3.4 |
| T799S | V747I | 0 | 3.5 | 10.5 | 7.0 |
| V800I | A414T | 0 | 3.5 | 2.6 | 3.1 |
| E503A | N533S | 0 | 2.8 | 5.4 | 4.1 |
| P509A | I301M | 0 | 6.3 | 1.3 | 3.8 |
| P509A | G137S | 0 | 6.3 | 3.8 | 5.1 |
| E503A | R596Q | 0 | 2.8 | 6.9 | 4.9 |
| E503A | S814F | 0 | 2.8 | 11.3 | 7.1 |
| V705M | A64S | 0 | 3.9 | 3.9 | 3.9 |
| A794T | L462V | 0 | 1.5 | 4.6 | 3.1 |
| L207V | N193S | 0 | 5.0 | 2.4 | 3.7 |
| E669K | N413Y | 0 | 4.3 | 6.7 | 5.5 |
| V735L | V233A | 0 | 5.2 | 5.8 | 5.5 |
| V735L | R66K | 0 | 5.2 | 3.9 | 4.6 |
| E278K | A64T | 0 | 5.4 | 3.9 | 4.7 |
| V735L | A569T | 0 | 5.2 | 4.1 | 4.7 |

Del = Deletion
N/FS = Nonsense OR frameshift mutation
Intron = Intronic mutation
MLS = Mutation in Mitochondrial Leader Sequence

| | | | | | |
|---|---|---|---|---|---|
| A569T | MLS | 0 | 4.1 | 8.7 | 6.4 |
